# An inherent T cell-activating mRNA delivery platform for *in vivo* CAR T generation

**DOI:** 10.1101/2025.11.19.688592

**Authors:** Qiannan Cao, Yingli Yao, Wenming Zheng, Hongqian Liu, Mingxia Jiang, Dayang Xie, Siting Zhang, Pijun Su, Huilin Yuan, Xiaoyuan Chen, Huapan Fang, Huayu Tian

## Abstract

The clinical success of CAR T cell therapy has underscored the need for scalable, non-invasive strategies for *in vivo* T cell engineering. Although mRNA delivery offers a promising alternative, current lipid-based vectors lack intrinsic T cell tropism and often rely on antibody conjugation, complicating manufacturing and raising safety concerns. Here we present an inherent T cell-activating polymer–lipid nanoparticle (**ERTLNPs**) that enable ligand-free, efficient mRNA transfection and activation of T cells *in vivo*. **ERTLNPs**, composed of p-toluenesulfonyl arginine (RT)-functionalized polyethylenimine (denoted as PEI-RT) and helper lipids, preferentially accumulate in the spleen following systemic administration. Without exogenous stimulation, **ERTLNPs** intrinsically activate T cells, triggering robust mRNA expression and proliferation. Mechanistically, **ERTLNPs** engage the PI3K/AKT/mTOR signaling axis to reprogram T cell metabolism, promoting expansion while restraining exhaustion. Systemic delivery of mRNA encoding fibroblast activation protein chimeric antigen receptor (mFAP CAR) *via* **ERTLNPs** leads to *in situ* generation of functional CAR T cells, which efficiently eliminate pathological fibroblasts in models of cancer and fibrosis, with minimal off-target effects. This ligand-free, metabolically reprogramming mRNA delivery platform provides a clinically translatable approach for *in vivo* CAR T cell generation and broadens the landscape of non-viral immunotherapeutic engineering.

## Introduction

T cell-based immunotherapy has gained considerable attention due to its promising potential in cancer treatments, particularly following the approval of chimeric antigen receptor (CAR) T cell therapy by U. S. Food and Drug Administration in 2017^1,2^. The clinical success of CAR T cell therapy in cancer treatment has facilitated its exploration in other disease areas^3^. Although *ex vivo* engineered CAR T cells have achieved significant breakthroughs in treating diseases such as hematologic malignancies, *ex vivo* CAR T cell therapy involves complex procedures and high costs, greatly limiting its clinical application^4–6^. mRNA technology has rapidly advanced in the treatment and prevention of diseases such as infectious diseases and cancer^7,8^. The *in vivo* generation of CAR T cells *via* mRNA-mediated delivery of CAR-encoding constructs directly into host T cells represents a promising alternative strategy, bypassing the need for *ex vivo* cell manipulation. However, most current mRNA delivery systems based on lipid nanoparticles (LNPs) primarily target the liver^9,10^, and monoclonal antibody-conjugated LNPs, such as anti-CD3 LNPs, have been shown to induce T cell exhaustion after treatment, raising concerns for *in vivo* CAR T therapy^11^. Recent studies have developed extrahepatic mRNA delivery carriers that do not require targeting ligand modifications, such as selective organ-targeting (SORT) LNPs^12^ and charge-altering releasable transporters polymers^13^. Although these delivery systems exhibit good mRNA transfection efficiency *in vivo*, the transfection mechanisms in T cells have not been fully elucidated. Therefore, clarifying the mechanisms of mRNA transfection in T cells *in vivo* will provide essential theoretical guidance for *in vivo* T cell engineering and facilitate the development of more efficient and safer CAR T therapy strategies.

Activation is critical for T cell proliferation, differentiation, and transfection^14^. Upon activation, T cells enter an active proliferative phase, facilitating the integration or efficient expression of foreign genes^15,16^. In recent years, the rapid advancement of biomaterials has driven the development of T cell activation platforms^15,17,18^. These platforms often mimic the T cell activation process by binding to anti-CD3 and anti-CD28 antibodies, such as modifying anti-CD3 and anti-CD28 antibodies on polymers^19^. However, these antibody-based activation platforms involve complex manufacturing processes, exhibit substantial batch-to-batch variability, and often display limited T cell activation efficiency. More critically, their reliance on TCR complex-mediated stimulation can induce excessive T cell activation, which under chronic exposure frequently leads to T cell dysfunction and exhaustion^20,21^. These limitations highlight the need for improved activation strategies.

In this study, we developed an inherent T-activating mRNA delivery platform for *in situ* CAR T cell generation. Specifically, we utilized *p*-toluenesulfonyl arginine (RT)-modified polyethylenimine (denoted as PEI-RT) as an ionizable component, which was co-assembled with helper lipid (DSPC), PEG-lipid (DMG-PEG2000), and cholesterol to form **PEI-RT**-based lipid nanoparticles (**ERTLNPs**) for mRNA delivery and *in vivo* T cell gene editing. In contrast to conventional Dynabeads, **ERTLNPs** circumvent the need for antibody conjugation and achieves superior T cell activation efficiency, thereby facilitating enhanced proliferation and transfection outcomes. Additionally, following **ERTLNPs** treatment, the proliferating T cells exhibit lower levels of exhaustion. Further investigation into the activation mechanism demonstrated that **ERTLNPs** reprograms T cell metabolism and effector functions primarily through modulating PI3K/AKT/mTOR signaling axis, enabling efficient T cell activation and transfection. Gene sequencing results further revealed that significant upregulation of genes associated with T cell activation, metabolic reprogramming, and effector functionality in **ERTLNPs**-treated T cells (**Fig. 1**).

**Fig. 1.**
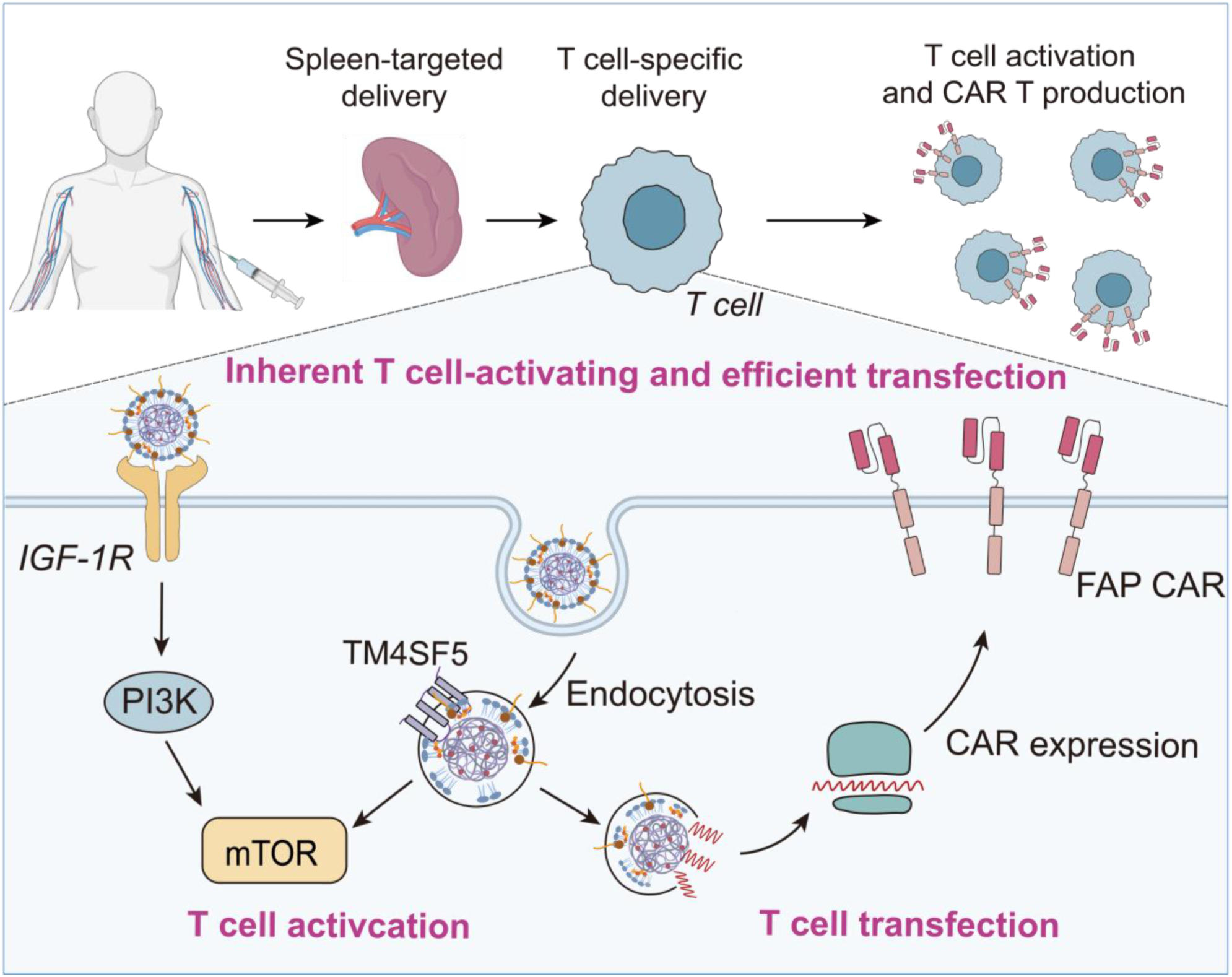
Illustration of ERTLNPs-mediated *in vivo* T cell activation and transfection. Following systemic administration, **ERTLNPs** preferentially accumulate in the spleen and efficiently deliver mRNA into T cells. **ERTLNPs**-mediated activation of naive T cells involves two distinct mechanisms: **i**) **ERTLNPs** bind to IGF-1 receptors (IGF-1R) on naive T cells, triggering the IGF-1R/PI3K/mTOR signaling pathway, thereby initiating T cell activation; **ii**) internalized **ERTLNPs** interact with the lysosomal transmembrane protein TM4SF5, activating the TM4SF5/mTOR signaling pathway, which further promotes T cell activation. Moreover, activated T cells exhibit enhanced metabolic activity and proliferation capability, facilitating efficient mRNA transfection and robust *in vivo* CAR T generation.

Notably, **ERTLNPs** exhibited spleen-selective tropism following systemic administration and achieved robust T lymphocyte transfection. Using mRNA encoding fibroblast activation protein chimeric antigen receptor (mFAP CAR) as the therapeutic gene, **ERTLNPs**-mediated systemic delivery of FAP CAR-encoding mRNA induced *in vivo* generation of potent FAP CAR T cells, demonstrating their translational potential for adoptive immunotherapy. In models of pancreatic cancer, pulmonary fibrosis, and hepatic fibrosis, **ERTLNPs** treatment significantly increased FAP CAR T cell infiltration at lesion sites, effectively eliminated activated fibroblasts in the affected areas, exhibited superior therapeutic efficacy compared to first-line clinical drugs with negligible immune-related side effects. In summary, we developed an “off-the-shelf” T cell activation platform that enables efficient and safe *in vivo* T cell activation and transfection for gene editing to treat various diseases. This activation strategy helps provide a novel paradigm for T cell activation and expands the current toolbox for CAR T cell therapy.

## Results

### Construction of efficient mRNA delivery system for *in vivo* CAR T cell therapy

The delivery efficiency of mRNA carriers is closely related to their chemical structure. In our previous study, we grafted *p*-toluene sulfonyl arginine onto polycations, thereby introducing multiple interactions including electrostatic, hydrogen bonding, and hydrophobic interactions between the carrier and the cell membrane, which significantly enhanced the nucleic acid delivery efficiency of the carrier^22^. To construct an efficient mRNA delivery vehicle, the *p*-toluene sulfonyl arginine-functionalized polyethylenimine (abbreviated as **PEI-RT**) was selected as the ionizable polymer component (**Fig. 2a**). The successful synthesis of PEI-RT was confirmed by hydrogen nuclear magnetic resonance (^1^H NMR) spectra indicated (**Supplementary Fig. 1**). Then, the obtained PEI-RT, 1,2-distearoyl-sn-glycero-3-phosphocholine (DSPC), 1,2-dimyristoyl-snglycerol-methoxypolyethylene glycol 2000 (DMG-PEG), cholesterol, and mRNA were assembled into a polymer-lipid hybrid mRNA delivery system (termed as **ERTLNPs**) through a dialysis process. To identify the roles of each component of **ERTLNPs** in mRNA transfection *in vivo*, we firstly investigated the effect of only three components on mRNA transfection efficiency (**Supplementary Fig. 2**). It was observed that the delivery system composed of only DSPC, DMG-PEG, and cholesterol exhibited negligible mRNA transfection performance after systemic administration, indicating the essential role of the ionizable polymer component in mRNA delivery. (**Fig. 2b**). Moreover, we found that the absence of DMG-PEG significantly reduced mRNA transfection efficiency compared to the omission of cholesterol or DSPC (**Supplementary Fig. 2a&b**), which is consistent with previous reports^23,24^.

**Fig. 2.**
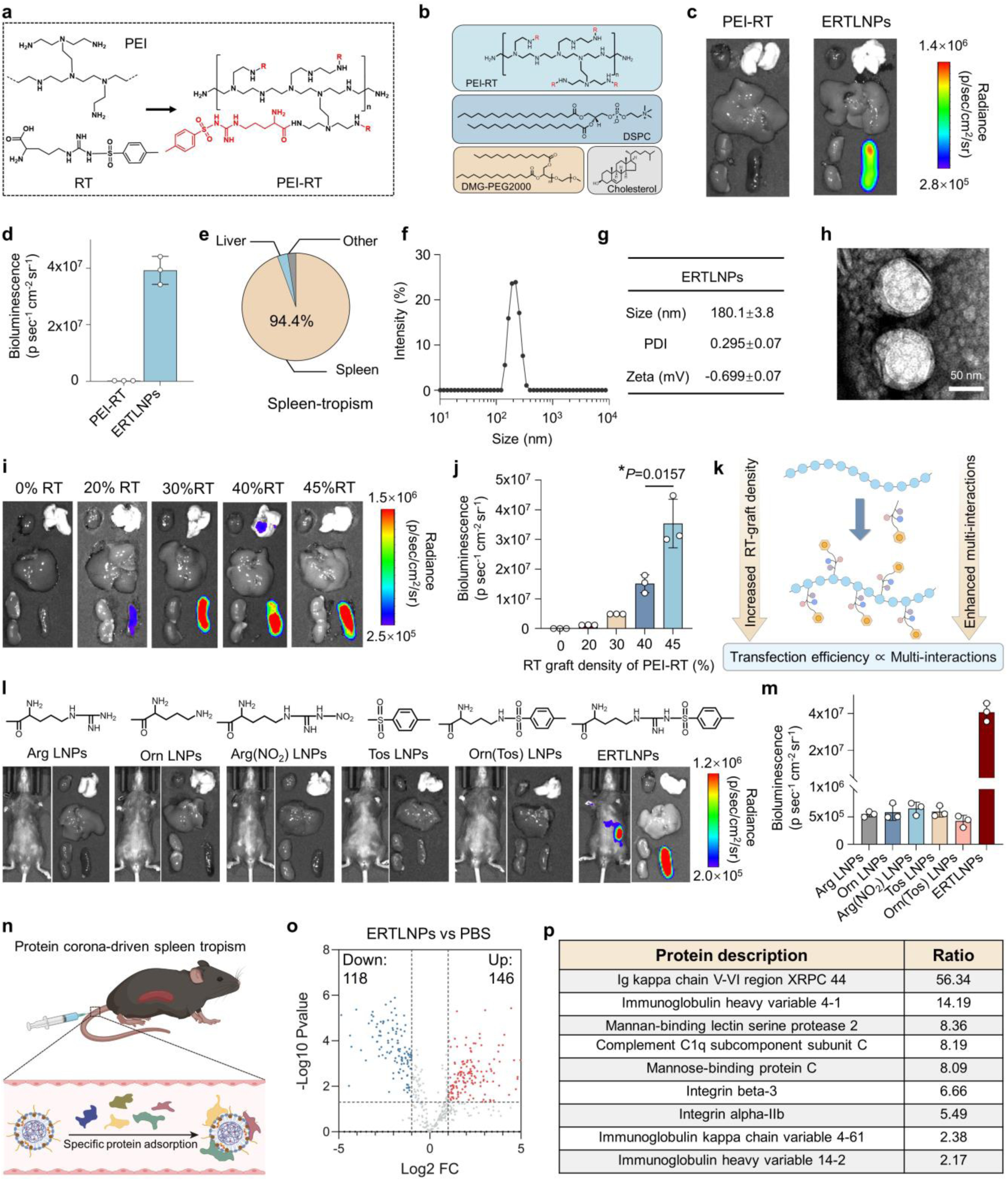
ERTLNPs mediate efficient mRNA delivery *in vivo*. **a,** The synthetic route of the ionizable polymer component PEI-RT. **b,** Composition of **ERTLNPs**. **c,d,** Representative *ex vivo* images (**c**) and quantification of luminescence **(d)** in main organs (heart, liver, spleen, lung, and kidney) from C57 mice 12 hours after systemic administration of Luc mRNA (denoted as mLuc)-loaded PEI-RT and **ERTLNPs** (0.5 mg kg^−1^ mLuc). **e**, Pie chart analysis of transfection efficiency in major organs using **ERTLNPs**. **f-h,** Hydrodynamic particle size, polydispersity index (PDI) and zeta potential **(f,g)** and transmission electron microscopy (TEM) image (**h**) of **ERTLNPs**. Scale bar: 50 nm. **i,j,** Representative *ex vivo* images (**i**) and quantification of bioluminescence **(j)** in main organs (heart, liver, spleen, lung, and kidney) from C57 mice 12 hours after systemic administration of mLuc-loaded **ERTLNPs** in which PEI-RTs contains different numbers of grafted RT (0.5 mg kg^−1^ mLuc).**k,** Illustration of the relationship between RT-mediated multiple interactions and transfection efficiency *in vivo*. **l,m,** Representative *in vivo* and *ex vivo* images (**l**) and quantification of bioluminescence **(m)** in main organs (heart, liver, spleen, lung, and kidney) from C57 mice 12 hours after systemic administration of mLuc-loaded Arg LNPs, Orn LNPs, Arg(NO_2_) LNPs, Tos LNPs, Orn(Tos) LNPs, or **ERTLNPs** (0.5 mg kg^−1^ mLuc). **n**, Schematic of the spleen-tropism mechanism of mRNA-loaded **ERTLNPs** following systemic administration. That is, following systemic administration, **ERTLNPs** bind to endogenous plasma proteins to form a protein corona, facilitating spleen-specific tropism. **o,** Volcano plot of differential protein corona profiles on **ERTLNPs** versus PBS. **p,** Relative quantification analysis of the top 9 spleen-associated proteins bound to **ERTLNPs** in plasma. Data in **d**, **j**, and **m** are shown as mean ± SD, n = 3 biologically independent samples. *P* values were determined by a two-tailed Student’s *t*-test in **j** as indicated in the figures. \**P* < 0.05. A representative image of three independent samples from each group is shown in **c**, **i**, and **l**.

Some studies have shown that PEG-lipid is an essential component of lipid nanoparticle (LNP) formulations, and the amount of PEG-lipid can significantly influence mRNA transfection efficiency while the cholesterol has a relatively minor effect on endosomal release^25^. Therefore, to achieve optimal *in vivo* mRNA transfection with **ERTLNPs**, we adjusted the proportions of DMG-PEG and cholesterol while keeping the amounts of the other two components constant. According to *ex vivo* fluorescence imaging results, the top candidate **ERTLNPs** exhibited selective expression in the spleen after systemic administration, at which point the spleen accounted for 94.4% of the total fluorescence among major organs (heart, liver, spleen, lungs, kidneys), highlighting their tissue-specific delivery capability (**Fig. 2c-e**). This spleen-targeting property minimized the impact on non-splenic organs and reduced potential safety concerns associated with off-target effects^26^. This was markedly different from conventional lipid nanoparticles such as SM102 LNP, which primarily mediate mRNA expression in the liver (**Supplementary Fig. 3**).

The *in viv*o mRNA transfection performance of polymer-lipid nanoparticles was closely associated with physicochemical properties^27–29^. It was observed that **ERTLNPs** exhibited hydrodynamic diameter (180.1 ±3.8 nm), a narrow particle size distribution (polydispersity index, PDI < 0.3), and a nearly neutral surface charge (0.699 ± 0.07 mV) (**Fig. 2f,g**), suggesting that **ERTLNPs** could assemble into nanoparticles both in the presence and absence of mRNA through multiple interactions. Furthermore, transmission electron microscopy (TEM) revealed a clear core–shell structure of the **ERTLNPs** (**Fig. 2h**). **ERTLNPs** formulations achieved mRNA encapsulation efficiencies above 85% (**Supplementary Fig. 4**). In addition, *ex vivo* fluorescence imaging results further demonstrated that **ERTLNPs** loaded with Cy5-mRNA selectively accumulated in spleen tissue following systemic administration (**Supplementary Fig. 5**).

Furthermore, to further investigate the effect of the molar grafting percentages of RT onto polyethyleneimine oligomers on the transfection performance of **ERTLNPs**, we synthesized PEI-RTs with varying RT grafting levels (20%, 30%, 40%, and 45%). The impact of **RT** grating contents on the *in vivo* mRNA transfection of **ERTLNPs** was evaluated at the same mass percentage of PEI-RTs. The bioluminescence imaging results showed that the transfection efficiency of **ERTLNPs** increased significantly with higher RT grafting ratios in PEI-RT (**Fig. 2i&j**). This phenomenon was likely due to the fact that, for polymer–lipid nanoparticles, increasing the RT content strengthened the multiple interactions between PEI-RT and cell membrane—including electrostatic, hydrogen bonding, and hydrophobic interactions^22,30^, and the enhanced multiple interactions facilitated cellular uptake of **ERTLNPs**, thereby promoting their gene transfection (**Fig. 2k**).

To further elucidate the mechanism behind the excellent *in vivo* mRNA transfection performance of **ERTLNPs**, we prepared PEI modified with various functional moieties for *in vivo* mRNA transfection studies (**Fig. 2l&m**, **Supplementary Fig. 6-10**). These moieties included ornithine (Orn), arginine (Arg), p-toluenesulfonyl ornithine (Orn(Tos)), p-toluenesulfonyl (Tos), and nitroarginine (Arg(NO_2_)). Interestingly, among all the tested polymer–lipid nanoparticle formulations, the **ERTLNPs** group exhibited the most efficient mRNA transfection in the spleen. This finding further supported the notion that synergistic electrostatic, hydrogen bonding, and hydrophobic interactions significantly enhanced the *in vivo* mRNA delivery capability of polymer–lipid nanoparticles.

In addition, recent studies had shown that LNPs could form a protein corona by adsorbing various plasma proteins upon systemic administration, which might subsequently influence their tissue-specific distribution^31–33^. Therefore, a protein corona profiling experiment was conducted to analyze the proteins adsorbed onto the surface of **ERTLNPs** in plasma. It was observed that **ERTLNPs** adsorbed a total of 1,942 proteins, with 146 proteins significantly upregulated compared to the control group (**Fig. 2n-p**). Notably, nearly one-third of these proteins were involved in immune receptor-mediated binding processes, including immunoglobulins, complement proteins, and lectin-related proteins. Remarkably, the level of the Ig kappa chain protein in the **ERTLNPs** group was upregulated by 56.34-fold compared to the control group (**Fig. 2p**). The Ig kappa chain could interact with Fc receptors of immune cells *via* its Fc region and activated the complement system^34^. Given the spleen’s rich populations of immune cells, such interactions between protein corona and immune cells likely contributed to the efficient accumulation and transfection of **ERTLNPs** in the spleen after systemic administration.

### ERTLNPs enable robust *in vivo* transfection in T cells following systemic administration

According to the aforementioned results, **ERTLNPs** selectively accumulate and mediate transfection in the spleen after intravenous injection. Given the diverse population of immune cells in the spleen, we investigated the immune cell types transfected by **ERTLNPs** *in vivo* using mTmG transgenic mice. Cre mRNA-loaded **ERTLNPs** were administered into mice *via* tail vein injection, and 48 hours post-injection, splenic immune cell transfection was analyzed using flow cytometry (**Fig. 3a**). It was observed that **ERTLNPs** efficiently transfected T cells within the spleen (**Fig 3b, c, and Supplementary Fig. 11**), highlighting their potential in T cell–mediated immunotherapies, particularly for chimeric antigen receptor (CAR) T cell therapy. Moreover, there was approximately 5% GFP^+^ of peripheral blood T cells in mTmG transgene mice (**Fig. 3d, Supplementary Fig. 12**). The immunofluorescence images further confirmed efficient GFP expression in the white pulp region of the spleen, with clear co-localization of GFP fluorescence and CD3^+^ T cells (**Fig. 3e, f**), consistent with the efficient transfection of splenic T cells observed post-systemic administration.

**Fig. 3.**
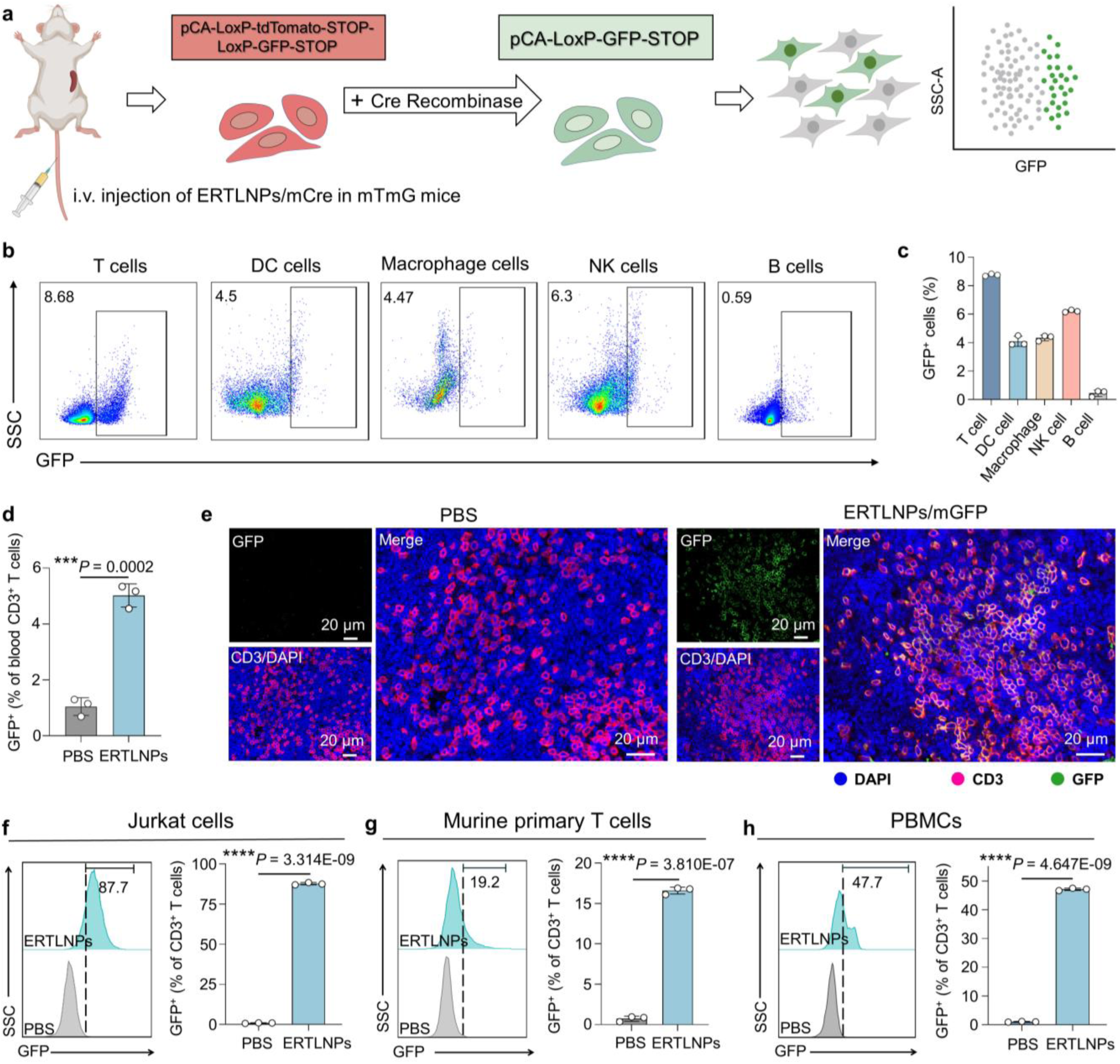
ERTLNPs enable highly efficient mRNA delivery to T cells *in vitro* and *in vivo*. **a**, Schematic of Cre mRNA (denoted as mCre) -loaded ERTLNP transfection in mTmG mice following systemic administration.**b**,**c**, Representative flow cytometry analysis (**b**) and relative quantification (**c**) of GFP^+^ cells in the splenic cells of mTmG mice 48 hours after the intravenous administration of **ERTLNPs**/mCre (1 mg kg^−1^ mCre). **d**, Relative quantification of GFP^+^ cells gating on CD45^+^ CD3^+^ cells in peripheral blood of mTmG mice 48 hours after the intravenous administration of **ERTLNPs**/mCre (1 mg kg^−1^ mCre) **e,** Representative immunofluorescence images of GFP^+^ CD3^+^cells in the spleen of mTmG mice 48 hours after the intravenous administration of **ERTLNPs**/mGFP (1 mg kg^−1^ mGFP). **f-h**, Representative flow cytometry analysis and relative quantification of GFP^+^ cells in Jurkat T cells (**f**), primary murine T cells (**g**), and PBMCs (**h**) after 24 hours of incubation with**ERTLNPs**/mGFP. Data in **c**, **d**, **f-h** are shown as mean ± SD, n = 3 biologically independent samples. *P* values were determined by a two-tailed Student’s *t*-test in **d**, **f**-**h** as indicated in the figures. \*\*\**P* < 0.001 and \*\*\*\**P* < 0.0001. A representative image of three independent samples from each group is shown in **b**, **e**, and **f-h**.

We further assessed the *in vitro* T cell transfection efficiency of **ERTLNPs**. It was observed that **ERTLNPs** could achieve nearly 90% cellular transfection in Jurkat T cells (**Fig. 3g**). Moreover, **ERTLNPs** transfected approximately 20% of primary T cells isolated from mouse spleens (**Fig. 3h**) and up to 50% of human peripheral blood mononuclear cells (PBMCs) (**Fig. 3i**). These results collectively demonstrated that **ERTLNPs** without targeting ligand modification, could selectively transfect T cells in the mouse spleen following systemic administration, and were also capable of transfecting primary T cells *in vitro*. This phenomenon highlighted the potential of **ERTLNPs** as a delivery platform for CAR T cell therapy.

### ERTLNPs can rapidly and efficiently activate primary T cells

Despite the exciting progress in immune cell transfection, studies utilizing non-targeted ligand-modified carriers for T cell transfection remain limited. Encouraged by the ability of **ERTLNPs** to efficiently transfect primary T cells, we further investigated the mechanism by which **ERTLNPs** mediated T cell transfection. During *in vitro* primary murine T cell culture, we were pleased to observe that primary T cells exhibited marked aggregation and proliferation after 24 hours of incubation with **ERTLNPs**, which was a characteristic hallmark of T cell activation (**Fig. 4a**). Generally, T cell activation was a prerequisite for effective mRNA transfection of primary T cells *in vitr*o^13,35–37^. T cell activation usually depended on treatment with anti-CD3 and anti-CD28 antibodies, which bunded to the CD3 and CD28 co-stimulatory receptors on the T cell surface to achieve robust activation. In this study, we further evaluated the activation and expansion capability of T cells treated with **ERTLNPs** *in vitro*. As shown in **Fig. 4b**, CD69 used as a marker of T cell early activation^36,38^, the proportion of CD69⁺ T cells significantly increased 24 hours after **ERTLNPs** treatment, reaching levels comparable to those induced by anti-CD3/CD28 stimulation. In contrast, the commercial transfection reagents such as MC3 LNP, SM102 LNP, and ALC0315 LNP barely induced the activation in primary T cells. Moreover, within the initial 6-hour post-treatment window, the percentage of CD69⁺ T cells in the **ERTLNPs** group rose more rapidly than that in the anti-CD3/CD28 group, and **ERTLNPs** were able to sustain a high activation phenotype even after 48 hours. These results clearly suggested that **ERTLNPs** could rapidly and efficiently activate primary T cells (**Fig. 4c**).

**Fig. 4.**
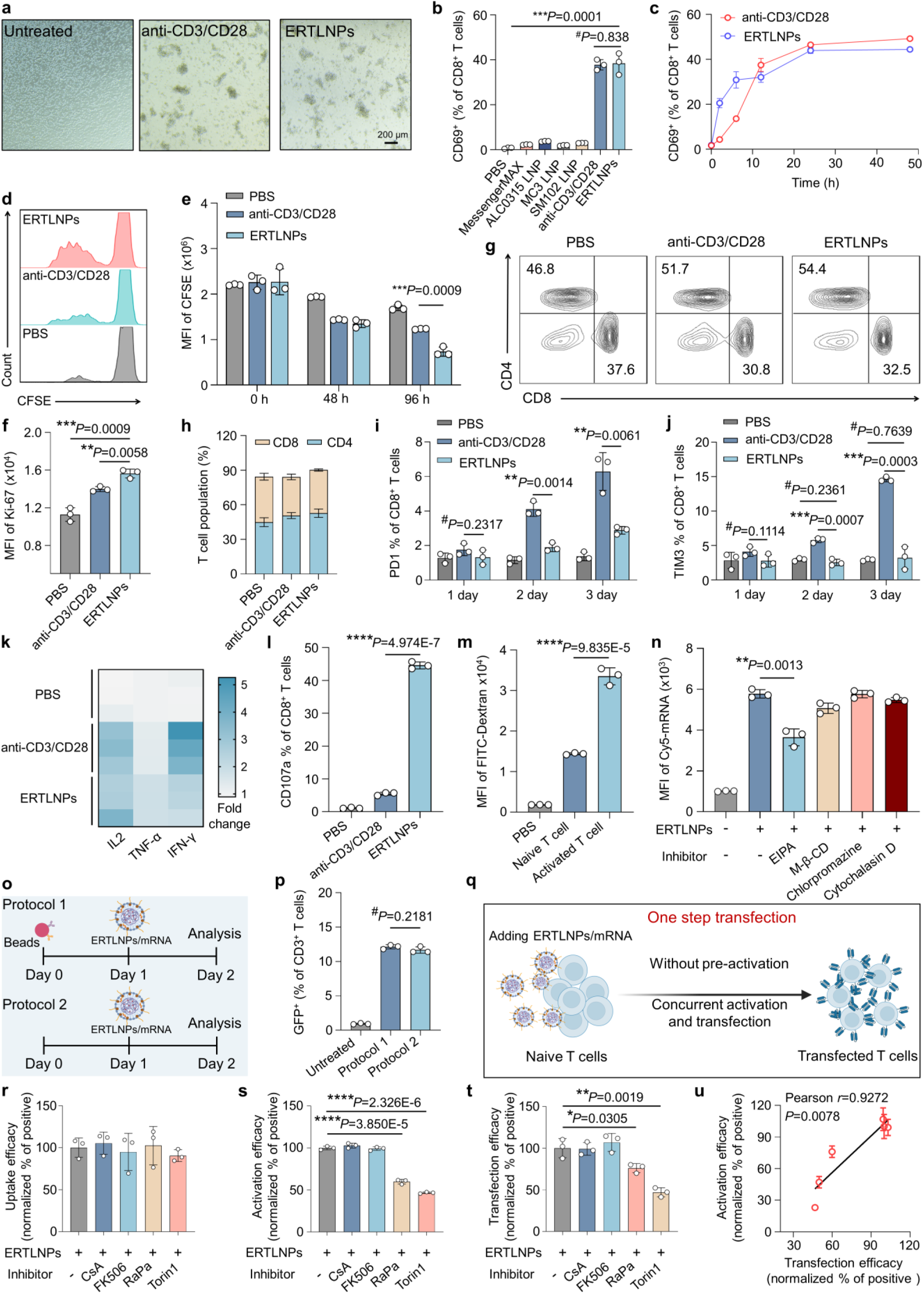
ERTLNPs significantly potentiate activation and effector functionality of primary T cells. **a**, Representative brightness images of primary murine T cells after 24 hours of incubation with anti-CD3/CD28 or **ERTLNPs**. Scale bar: 200 μm. **b**, Relative quantification of CD69^+^ cells in primary murine CD8^+^ T cells after 24 hours of incubation with **ERTLNPs**. Messenger MAX, ALC0315 LNP, MC3 LNP, SM102 LNP and antiCD3/CD28 were used as control. **c**, Relative quantification curves of CD69^+^ cells in primary murine CD8^+^ T cells after incubation with anti-CD3/CD28 or **ERTLNPs** for different durations (2, 6, 12, 24, and 48 hours). **d**,**e**, Representative flow cytometry analysis (**d**) and relative quantification of (**e**) of CFSE-labelled primary murine T cells after 0, 48, and 96 hours of incubation with anti-CD3/CD28 or **ERTLNPs**. **f**, Relative quantification of Ki-67 levels in primary murine T cells after 96 hours of incubation with **ERTLNPs**. **g**,**h**, Representative flow cytometry analysis (**g**) and relative quantification (**h**) of CD8^+^ and CD4^+^ cells in primary murine T cells after 72 hours of incubation with anti-CD3/CD28 or **ERTLNPs**.The higher CD4/CD8 ratio in primary T cells incubated with **ERTLNPs** than that incubated with anti-CD3/CD28 indicates that **ERTLNP**s can induce more robust and sustained T cell activation. **i**,**j**, Relative quantification of PD-1^+^ cells (**i**) and TIM3^+^ cells (**j**) in primary murine CD8^+^ T cells after incubation with anti-CD3/CD28 or **ERTLNPs** for different durations (1, 2, and 3 days). **k**, Heat map of relative levels of IL-2, TNF-α, and IFN-γ expression in primary murine T cells after incubation with anti-CD3/CD28 or **ERTLNPs** for 48 hours. **l**, Relative quantification of CD107a^+^ cells in CD8^+^ T cells after 24 hours of incubation with anti-CD3/CD28 or **ERTLNPs** using flow cytometry. **m**, Relative quantification of of FITC-dextran uptake by naive T cells and activated T cells. **n**, Cellular uptake efficiency of mRNA-loaded **ERTLNPs** in naive T cells pretreated with various endocytosis inhibitors by flow cytometry, mRNA was labelled with Cy5. **o**, Schematic of two different *in vitro* transfection protocols mediated by **ERTLNPs.** In one protocol, naive T cells are pretreated with anti-CD3/CD28 Dynabeads for 24 hours before the addition of **ERTLNPs**, while in the other protocol, **ERTLNPs** are added without prior Dynabeads pretreatment. **p**, Relative quantification of GFP^+^ cells in transfected T cells according to the protocols as indicated in **o**. **q**, Schematic of one-step transfection of primary T cells mediated **ERTLNPs** without Dynabeads stimulation. **r-t**, Uptake efficiency (**r**), activation efficiency (**s**) and transfection efficiency (**t**) of mRNA-loaded **ERTLNPs** in naive T cells pretreated with various activation inhibitors by flow cytometry. u, Correlation analysis between activation efficacy (CD69^+^) and transfection efficiency (GFP^+^ cells) of naive T cells mediated by **ERTLNPs**. Data in **b**, **c**, **e**, **f**, **i**, **j**, **l-n**, **p**, and **r-u** are shown as mean ± SD, n = 3 biologically independent samples. *P* values were determined by a two-tailed Student’s *t*-test in **b, c, e, f, h-n**, and **r**-**u** as indicated in the figures. *^#^P* > 0.05, \**P* < 0.05, \*\**P* < 0.01, \*\*\**P* < 0.001, and \*\*\*\**P* < 0.0001. A representative image of three independent samples from each group is shown in **a**, **d**, and **g**.

To further explore the mechanism of **ERTLNPs**-mediated T cell activation, we evaluated the ability of PEI-RT alone to activate T cells. It was found that at the same polymer dose, the T cell activation efficiency of **ERTLNPs** was significantly higher than that of PEI-RT alone (**Supplementary Fig. 13a**). Additionally, **ERTLNPs** exhibited markedly better mRNA transfection performance in primary T cells than PEI-RT alone (**Supplementary Fig. 13b**). When PEI-RT was removed from **ERTLNPs**, the remaining three components had almost no ability to activate T cells (**Supplementary Fig. 14**). Moreover, removal of PEG, DSPC, or cholesterol from **ERTLNPs** severely impaired T cell activation (**Supplementary Fig. 15**), suggesting that the structural assembly of **ERTLNPs** played an important role in T cell activation.

In addition, we also further evaluated the effect of different polymers and their corresponding polymer-lipid nanoparticles on T cell activation (**Supplementary Fig. 16**). It was observed that different polymer-lipid nanoparticles showed significantly different T cell activation capabilities. Among them, **ERTLNPs** exhibited the most robust T cell activation. Furthermore, the similar trend was obtained in T cell activation efficiency for these polymers alone, and the corresponding polymer-lipid nanoparticles had higher activation efficiencies than the polymers alone. This phenomenon was consistent with our findings in the **ERTLNPs** system. Interestingly, these polymer-lipid nanoparticles also exhibited similar trends in their transfection efficiency in primary T cells (**Supplementary Fig. 17**). This indirectly indicated that **ERTLNPs** could rapidly induce the activation of primary T cells, thereby promoting T cell transfection. Typically, incubation of primary T cells with anti-CD3/CD28 antibodies not only primed the activation of primary T cells but also contributed to their proliferation^39^. Given the excellent T cell activation capability of **ERTLNPs**, we further investigated whether **ERTLNPs** could also enhance T cell proliferation. The primary T cells were first labeled with carboxyfluorescein succinimidyl ester (CFSE), and then incubated with either anti-CD3/CD28 antibodies or **ERTLNPs** for 48 or 96 hours. It was observed that after 48 hours of incubation, both the anti-CD3/CD28 group and the **ERTLNPs** group significantly promoted the proliferation of primary T cells. Moreover, after 96 hours of incubation, the **ERTLNPs** group exhibited a higher levels of cell proliferation compared to the anti-CD3/CD28 group. In addition, the primary T cells incubated with **ERTLNPs** underwent more rounds of cell division (**Fig. 4d, e**). Similarly, the **ERTLNPs** group displayed a higher mean fluorescence intensity of Ki-67 than the anti-CD3/CD28 group (**Fig. 4f**). These results suggested that **ERTLNPs** promoted T cell proliferation more significantly than anti-CD3/CD28 antibodies.

Furthermore, the proportion of CD4⁺ cells in primary T cells treated with **ERTLNPs** (approximately 54.4% after 72 hours) was notably higher than that in the anti-CD3/CD28 group (**Fig. 4g, h**). Previous studies have reported that strong and sustained T cell activation could increase the percentage of CD4⁺ T cells^40,41^, which would be beneficial for *in vivo* chimeric CAR-T therapy. Despite promoting rapid T cell activation and proliferation, the **ERTLNPs** group induced lower expression levels of exhaustion markers such as programmed death-1 (PD-1) and T cell immunoglobulin and mucin-domain containing-3 (TIM-3) compared to the anti-CD3/CD28 group, indicating that **ERTLNPs** did not exacerbate T cell exhaustion (**Fig. 4i, j**). We further evaluated the *in vivo* T cell activation induced by systemic administration of **ERTLNPs**. it was observed that **ERTLNPs** could activate approximately 50% of splenic CD3⁺ T cells with negligible exhaustion observed (**Supplementary Fig. 18**), confirming that **ERTLNPs** contributed to the activation of T cells *in vivo*. Moreover, compared to the untreated group, the **ERTLNPs** group showed higher levels of pro-inflammatory cytokine secretion, such as interleukin-2 (IL-2), interferon-gamma (IFN-γ), and tumour necrosis factor-alpha (TNF-α) after *in vitro* culture of primary T cells (**Fig. 4k**). In addition, CD107a, a marker for degranulation of CD8⁺ T cells, was also highly expressed in CD8⁺ T cells after incubation with **ERTLNP**s, as indicated by a marked increase in the percentages of CD107a⁺ CD8⁺ T cells (**Fig. 4l**).

Activation was typically a prerequisite for efficient mRNA transfection in primary T cells. according to the above findings, it was concluded that **ERTLNPs** could efficiently promote T cell activation and facilitate highly efficient gene transfection. However, the cellular uptake was a critical process in gene transfection. It was observed that activated T cells showed significantly enhanced uptake capability (**Fig. 4m**), which would contribute to gene transfection. To further elucidate the mechanism of T cell activation by **ERTLNPs**, we evaluated the uptake efficiency of **ERTLNPs**/Cy5-mRNA in primary T cells. It was found that Cy5-mRNA-loaded **ERTLNPs** showed efficient cellular uptake efficacy (**Supplementary Fig. 19**). To investigate the cellular uptake mechanism of **ERTLNPs**, primary T cells were pretreated with various endocytosis inhibitors. It was found that treatment with 5-(N-ethyl-n-isopropyl) amiloride (EIPA, a macropinocytosis inhibitor) significantly reduced the uptake efficiency of Cy5-mRNA-loaded **ERTLNPs**. The Na⁺/H⁺ exchanger on the cell membrane usually regulated -mediated cellular uptake, and EIPA could block its function^42^. These results suggested that macropinocytosis was the primary pathway for **ERTLNPs** entry into T cells (**Fig. 4n**).

Given the superior activation performance of **ERTLNPs** over anti-CD3/CD28 antibodies, we wondered whether **ERTLNPs** could achieve efficient mRNA transfection in primary T cells without pre-activation using anti-CD3/CD28 antibodies. Thus, we designed the experimental groups as follows: in the first group, primary T cells were pretreated with anti-CD3/CD28 Dynabeads for 24 hours, followed by incubation with **ERTLNPs**/mGFP complexes for another 24 hours; in the second group, primary T cells were directly incubated with **ERTLNPs**/mGFP without any prior activation (**Fig. 4o**). As shown in **Fig. 4p**, **ERTLNPs** exhibited excellent mRNA transfection efficiency in primary T cells regardless of anti-CD3/CD28 Dynabeads pretreatment. This demonstrated that **ERTLNPs** could enable efficient one-step transfection in primary T cells, eliminating the complicated steps typically required by conventional transfection agents, such as antibody Dynabeads activation (**Fig. 4q**).

To further clarify the activation pathway of T cells by **ERTLNPs**, we investigated the effects of various inhibitors on primary T cell activation signaling. These inhibitors included cyclosporine A (CsA, a nuclear translocation inhibitor of nuclear factor of activated T cells), tacrolimus (FK506, a calcineurin inhibitor), rapamycin (RaPa, an inhibitor for mechanistic target of rapamycin (mTOR), and Torin1 (a potent inhibitor for mechanistic target of rapamycin complex 1 (mTORC1)). It was found that these inhibitors had negligible effects on the cellular uptake efficiency of **ERTLNPs** (**Fig. 4r**). However, both RaPa and Torin1 significantly suppressed T cell activation, indicating that the mTOR signaling pathway played a critical role in **ERTLNPs**-mediated T cell activation (**Fig. 4s**). We then further evaluated the effects of these inhibitors on the transfection performance of **ERTLNPs** in primary T cells. It was observed that both RaPa and Torin1 significantly reduced the transfection efficiency of **ERTLNPs** (**Fig. 4t**). Interestingly, when transfection efficiency was plotted as a function of T cell activation efficiency, a strong correlation was observed between **ERTLNPs**-mediated transfection efficiency and T cell activation efficiency following treatment with RaPa or Torin1 (**Fig. 4u**). Therefore, it was concluded that **ERTLNPs** could efficiently activate primary T cells and subsequently promote high-efficient mRNA transfection.

### ERTLNPs activate primary T cells *via* mTOR signaling

According to the results above, it was found that the mTOR signaling pathway played a critical role in **ERTLNPs**-mediated activation of primary T cells. mTOR was a conserved serine/threonine kinase that was essential for cell proliferation, growth, and metabolism. Previous results had shown that **ERTLNPs** encapsulated mRNA was internalized by primary T cells primarily *via* macropinocytosis (**Fig. 4n**). As shown in **Fig. 5a**, it was found that EIPA pretreatment impeded the activation of primary T cells. Studies have reported that activated T cells could uptake nutrients such as amino acids through macropinocytosis, which was essential for sustaining the activation of the mTOR signaling pathway^16^. This phenomenon highlighted the connection between the macropinocytic pathway and T cell activation.

**Fig. 5.**
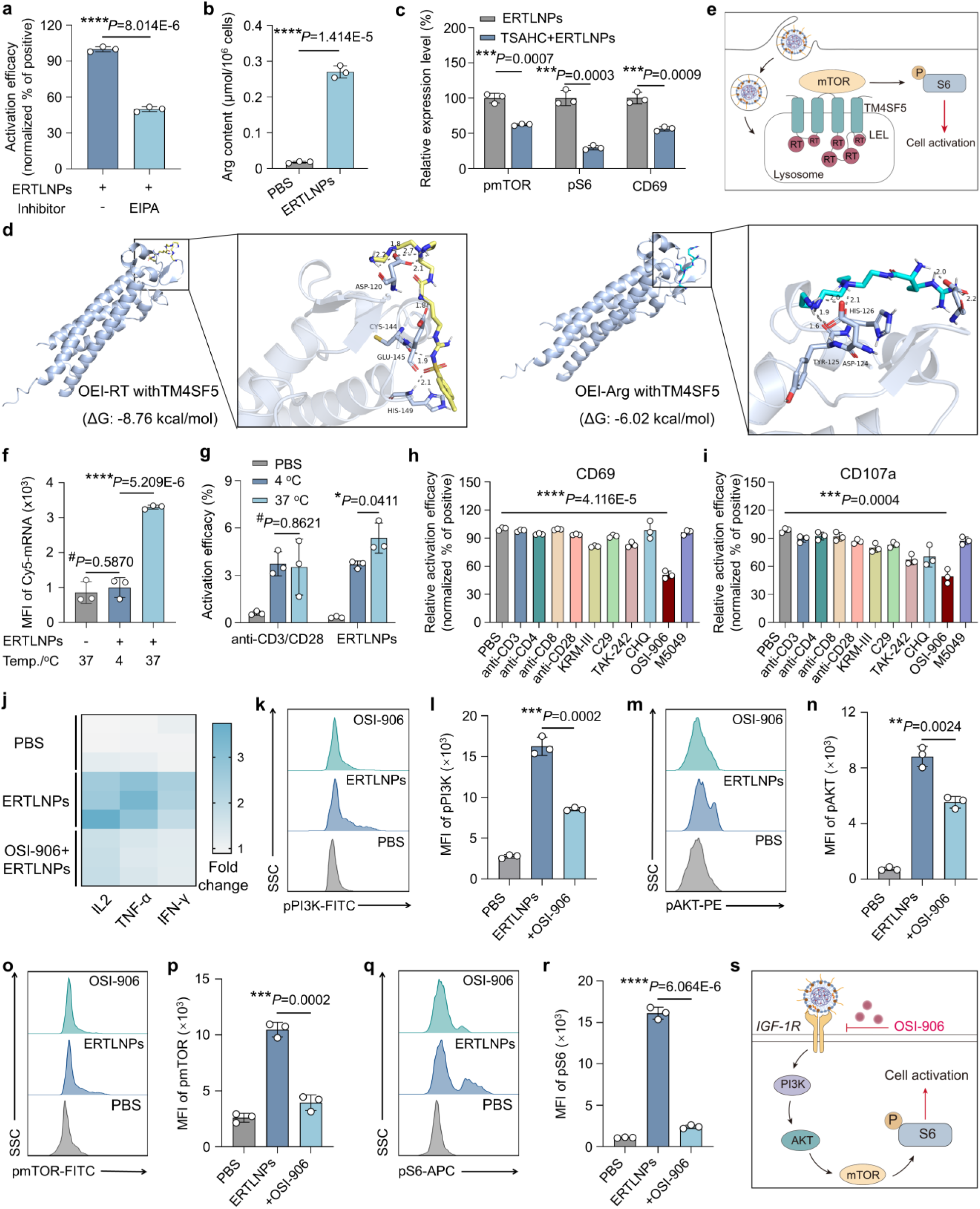
Mechanistic profiling of ERTLNPs-mediated T cell activation. **a**, Relative quantification of **ERTLNPs-**mediated activation efficiency (CD69 expression) of naive T cells with or without EIPA pretreatment (50 μM, macropinocytosis inhibitor). **b**, Content of arginine in naive T cells incubated with **ERTLNPs** for 24 hours. **c**, Relative contents of pmTOR, pS6 and CD69 expression in **ERTLNPs**-incubated naive T cells with or without TSAHC (100 μM) pre-treatment. **d**, Molecule docking analysis of PEI-RT (left) or PEI-Arg (right) with TM4SF5. **e**, Schematic of **ERTLNPs** internalized by primary T cells initiates TM4SF5/mTOR signaling pathway and promotes T cell activation. **f**,**g**, Uptake efficiency (**f**) and activation efficiency (**g**) of mRNA-loaded **ERTLNPs** in naive T cells incubated at 4 °C or 37 °C by flow cytometry. **h**,**i**, Relative quantification of **ERTLNPs**-mediated naive T cells activation (CD69 (**h**) and CD107a (**i**)) in which T cells are preincubated with antibodies or receptor inhibitors. anti-CD3 antibody, CD3 receptor inhibitor; anti-CD4 antibody, CD4 receptor inhibitor; anti-CD8 antibody: CD8 receptor inhibitor; anti-CD28 antibody, CD28 receptor; KRM-III (), TCR; C29, TCR2; Resatorvid (TAK-242), TLR4; Chloroquine (CHQ), TLRs; Linsitinib (OSI-906), dual inhibitor of the IR and IGF1R; Enpatoran (M5049), TLR7/8. **j**, Heatmap of relative levels of IL-2, IFN-γ, and TNF-α serectionin **ERTLNPs**-treated naive T cells with or without OSI-906 pretreatment (50 μM, IGF-1R inhibitor). **k**,**l**, Representative flow cytometric analysis (**k**) and relative quantification (**l**) of pPI3K expression in the **ERTLNPs**-treated naive T cells with or without OSI-906 pretreatment. **m**,**n**, Representative flow cytometric analysis (**m**) and relative quantification (**n**) of pAKT expression in the **ERTLNPs**-treated naive T cells with or without OSI-906 pretreatment. **o**,**p**, Representative flow cytometric analysis (**o**) and relative quantification (**p**) of pmTOR expression in the **ERTLNPs**-treated naive T cells with or without OSI-906 pretreatment. **q**,**r**, Representative flow cytometric analysis (**q**) and relative quantification (**r**) of pS6 expression in the **ERTLNPs**-treated naive T cells with or without OSI-906 pretreatment. **s**, Schematic of **ERTLNPs** engaging with IGF-1R on T cells to initiate PI3K/AKT/mTOR signaling pathway and activate naive T cells and effector functions of naive T cells. Data in **a-c**, **f-i**, **l**, **n**, **p**, and **r** are shown as mean ± SD, n = 3 biologically independent samples. *P* values were determined by a two-tailed Student’s *t*-test in **a-c**, **f-i**, **l**, **n**, **p**, and **r** as indicated in the figures. *^#^P* > 0.05, \*\**P* < 0.01, \*\*\**P* < 0.001, and \*\*\*\**P* < 0.0001. A representative image of three independent samples from each group is shown in **k**, **m**, **o,** and **q**.

The functional target of mTOR was a signaling hub located on the lysosomal surface, where amino acids transported into the lysosome triggered activation of the mTOR signaling pathway. For instance, lysosome-internalized arginine played a vital role in T cell proliferation, differentiation, and effector functions by modulating the mTORC1 pathway, thereby regulating metabolic reprogramming and immune responses of T cells^43^. Studies had shown that transmembrane 4 L six family member 5 (TM4SF5) acted as an arginine sensor on the lysosomal membrane and facilitated arginine-mediated priming of the mTOR pathway to activate primary T cells^44^. Considering that **ERTLNPs** were rich in arginine residues, we hypothesized that upon uptake by T cells, the arginine residues delivered into the lysosome would interact with TM4SF5, thereby promoting activation of the mTOR/S6 signaling pathway. To validate this hypothesis, the levels of intracellular arginine residues were first assessed in primary T cells using a detection kit. After incubating primary T cells with **ERTLNPs** for 24 hours, significant arginine residue signals were detected in the cell lysates (**Fig. 5b**). Then we prepared 4’-(p-Toluenesulfonylamide)-4-hydroxychalcone (TSAHC), an antagonist of TM4SF5^45^ (**Supplementary Fig. 20**), to block the interaction between **ERTLNPs**-derived arginine residues and TM4SF5 for inhibiting mTOR pathway activation. It was found that TSAHC pretreatment significantly reduced the phosphorylation levels of mTOR/S6 in **ERTLNPs**-treated primary T cells and markedly suppressed T cell activation (**Fig. 5c**). Based on these findings, we concluded that **ERTLNPs** were entered into lysosome *via* macropinocytosis, where their arginine residues interacted with TM4SF5, triggering the mTOR/S6 signaling pathway to activate primary T cells.

To further explore the interaction between arginine residues and TM4SF5, we performed molecular docking simulations using PEI-RT and PEI-Arg oligomers to evaluate their binding affinities to TM4SF5. The results showed that the binding free energy (ΔG) between PEI-RT oligomers and TM4SF5 was −8.76 kcal/mol, significantly lower than that of PEI-Arg oligomers (ΔG = −6.02 kcal/mol) (**Fig. 5d**), suggesting that a strong binding affinity formed between PEI-RT oligomers and TM4SF5. Further analysis revealed that these interactions between PEI-RT oligomers and TM4SF5 involved hydrophobic interactions, hydrogen bonding interactions, π-π interactions, and van der Waals forces, with van der Waals forces playing a prominent role (**Supplementary Fig. 21**). In addition, PEI-RT oligomers formed hydrogen bonds with amino acid residues such as Asp120, Cys144, Glu145, and His149 of TM4SF5. In contrast, PEI-Arg oligomers mainly formed hydrogen bonds and salt bridges with amino acid residues such as His126, Thr125, and Asp120, with weaker van der Waals forces (**Supplementary Fig. 22**). These observations suggested that **ERTLNPs**, once internalized by primary T cells, formed strong multiple non-covalent interactions between their arginine residues and TM4SF5 within lysosomes, priming the mTOR signaling pathway and promoting primary T cell activation.

In addition, in primary T cells, the mTOR/S6 signaling pathway can also respond to various surface receptor signals such as co-stimulatory receptors, Toll-like receptors (TLR2, TLR4), and growth factor receptors like IGF-1R^46,47^. These receptors, when stimulated by external agonists, could activate the intracellular mTOR signaling pathway to promote T cell activation. Therefore, we further investigated whether cell surface receptors were involved in T cell activation following incubation of primary T cells with **ERTLNPs**. When incubated at 4 °C, the uptake efficiency of **ERTLNPs** loaded mRNA by primary T cells was negligible, indicating that the **ERTLNPs**-mediated mRNA uptake process was energy-dependent (**Fig. 5f**). We then evaluated the activation efficiency of primary T cells mediated by anti-CD3/CD28 Dynabeads or **ERTLNPs** at both 4 °C and 37 °C. The results showed that anti-CD3/CD28 Dynabeads effectively activated primary T cells at both temperatures, indicating that low temperature treatment did not affect T cell activation response (**Fig. 5g**). In contrast, although **ERTLNPs** exhibited lower T cell activation efficiency at 4 °C compared to 37 °C, they still induced a noticeable activation effect at 4 °C. This phenomenon suggested that **ERTLNPs** not only activated primary T cells through intracellular signaling but also through the interactions with surface receptors.

To identify which cell surface receptor was involved in **ERTLNPs**-mediated T cell activation, primary T cells were pretreated with various cell receptor inhibitors to block the binding of **ERTLNPs** to the corresponding receptors and assessed the changes in T cell activation. It was found that OSI-906 (i.e., linsitinib), an IGF-1R inhibitor, significantly reduced CD69 and CD107a expression in primary T cells treated with **ERTLNPs**, indicating that OSI-906 effectively blocked **ERTLNPs**-induced T cell activation. In contrast, this effect was not observed with other receptor inhibitors (**Fig. 5h, i**). Additionally, OSI-906 pretreatment significantly reduced the levels of pro-inflammatory cytokines secreted by T cells (**Fig. 5j**). Furthermore, **ERTLNPs** incubation markedly increased the phosphorylation levels of the PI3K/AKT/mTOR signaling axis in primary T cells, while OSI-906 pretreatment significantly inhibited their phosphorylation (**Fig. 5k-r**). It was reported that external stimulation of IGF-1R axis would prime the downstream PI3K/AKT signaling pathway, which in turn activated the mTOR pathway^48^. These results suggested that **ERTLNPs** stimulated the IGF-1R receptor on T cell surfaces, activating the PI3K/AKT/mTOR signaling axis and thereby promoting the activation of primary T cells (**Fig. 5s**).

Taken together, it was concluded that **ERTLNPs** activated primary T cells *via* a dual mechanism: (1) interaction of **ERTLNPs** with IGF-1R receptor on T cells, and (2) interaction of lysosomal internalized **ERTLNPs**-derived arginine residues with TM4SF5. Both mechanisms converged to prime the downstream mTOR signaling pathway, ultimately promoting the activation of primary T cells.

### ERTLNPs drive inherent T cell activation and metabolic reprogramming

Based on the above results, **ERTLNPs** were capable of dual-mode activation of primary T cells *via* the mTOR signaling pathway. We next performed RNA sequencing to assess the differential gene expression profiles of primary T cells before and after incubation with **ERTLNPs** (**Fig. 6a**). The volcano plot revealed substantial transcriptomic differences in primary T cells pre- and post-incubation with **ERTLNPs** or anti-CD3/CD28 Dynabeads (**Fig. 6b, c**). Volcano plot analysis of representative gene expression indicated that compared to the untreated group, 2597 genes were significantly upregulated and 2964 genes were downregulated in the **ERTLNPs**-treated group. Moreover, in the anti-CD3/CD28 group, 3397 genes were significantly upregulated, and 3112 were downregulated compared to the untreated group. When comparing the **ERTLNPs** group to the anti-CD3/CD28 group, 1169 genes were uniquely upregulated, and 1216 genes were uniquely downregulated (**Fig. 6d**). The gene expression heatmap further revealed notable upregulation of genes associated with T cell activation, T cell metabolism, and the mTOR signaling pathway (**Fig. 6e**). Notably, the genes involved in glycolysis, such as hexokinase 2 (HK2) and glucose transporter 1 (GLUT1), were markedly upregulated in the **ERTLNPs** group, suggesting that upon activation, T cells relied on glycolytic pathways to meet their energy demands.

**Fig. 6.**
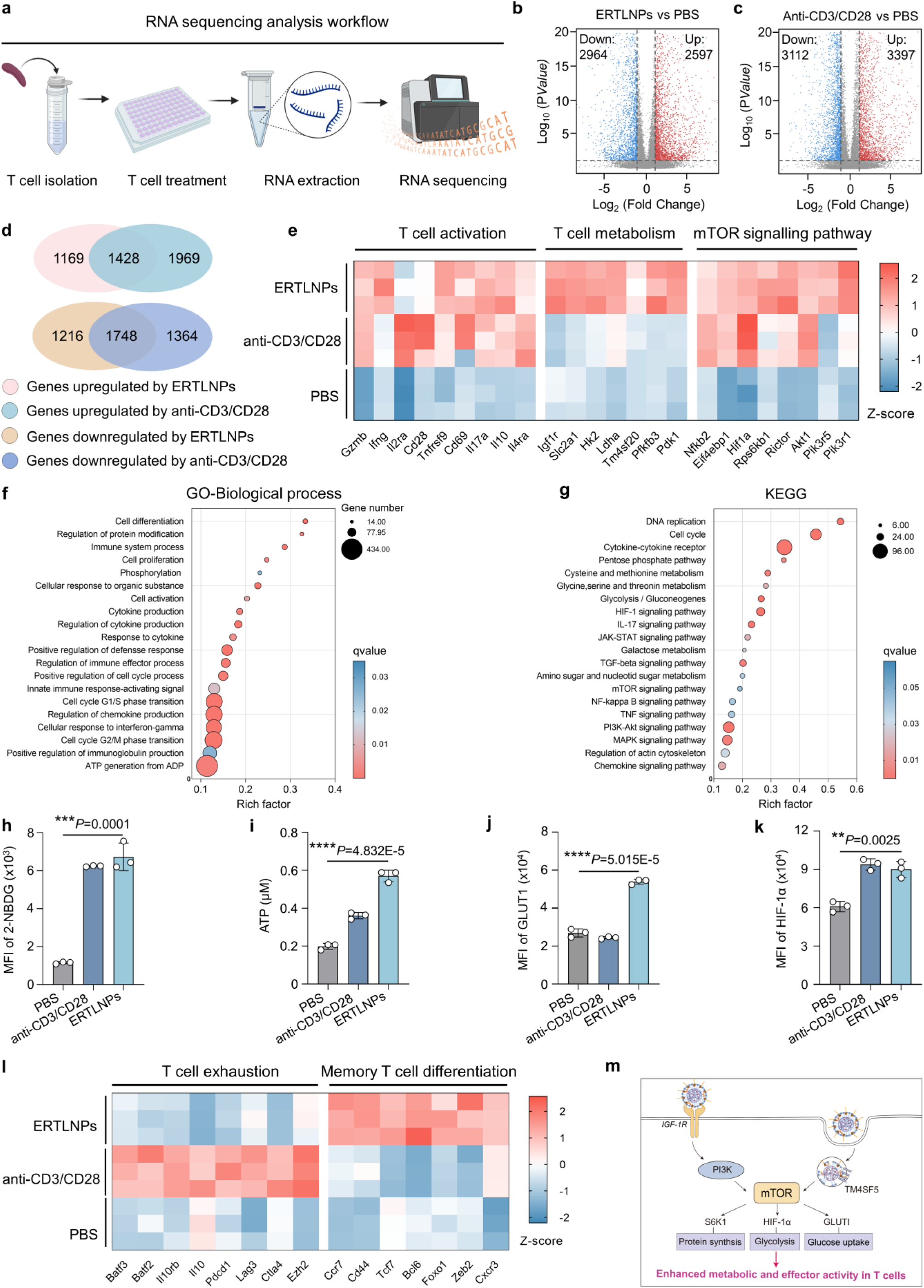
ERTLNPs reprogram cell metabolism to promote T cell activation. **a**, Schematic of RNA sequencing analysis in primary T cells treated with **ERTLNPs** or anti-CD3/CD28. **b-g**, Transcriptomic analysis of T cells after 24 hours of incubation with PBS, anti-CD3/CD28 or **ERTLNPs**: (**b**) Volcano plot of differentially expressed genes (DEGs) in T cells between the E**RTLNPs** and PBS groups, including 2597 upregulated and 2964 downregulated genes (|log_2_FC| >1, \**P* <0.05), FC: fold change; (**c**) Volcano plot of DEGs in T cells between the anti-CD3/CD28 and PBS groups, including 3397 upregulated and 3112 downregulated genes (|log_2_FC| >1, \**P* <0.05); (**d**) Venn diagrams showing the overlap of significantly upregulated (top) and downregulated (bottom) genes in T cells after treatment with **ERTLNPs** or anti-CD3/CD28, as compared to untreated controls. (**e**) A heat map of DEGs showing relative levels of genes related to T cell activation, T cell metabolism and the mTOR signaling pathway. Genes shown were selected from top differentially expressed genes between PBS, anti-CD3/CD28 and **ERTLNPs** in each pathway; (**f**) Gene Ontology (GO) enrichment analysis of the upregulated genes of the pathways of **ERTLNPs**-treated naive T cells; (**g**) Kyoto Encyclopedia of Genes and Genomes (KEGG) enrichment analysis of the upregulated genes of the pathways of **ERTLNPs**-treated T cells. **h**, Flow cytometry analysis quantifying 2-NBDG uptake efficiency in T cells pre-treated with anti-CD3/CD28 or **ERTLNPs** for 12 hours. **i**, Content of ATP of T cellss incubated with anti-CD3/CD28 or **ERTLNPs** incubation for 24 hours. **j**, Mean fluorescence intensity (MFI) of glucose transporter type 1 (GLUT1) expression on T cells incubated with anti-CD3/CD28 or **ERTLNPs for** 24 hours. **k**, MFI of HIF-α in T cells incubated with anti-CD3/CD28 or **ERTLNPs** for 24 hours. **l,** A heat map of DEGs showing the expression of genes related to T cell exhaustion and memory T cell differentiation incubated with PBS, anti-CD3/CD28, or **ERTLNPs**. **m**, Schematic of **ERTLNPs**-mediated dual-pathway initiation of mTOR signaling pathway promotes T cell metabolic and functional activation. **ERTLNPs** activate the mTOR signaling axis *via* two distinct mechanisms: engagement of IGF-1R on the T cell surface and lysosomal arginine sensing through TM4SF5. These pathways converge to upregulate downstream mTOR effectors, leading to enhanced protein synthesis and metabolic reprogramming. Taken together, these processes promote T cell metabolic fitness and effector activation. Data in **h-k** are shown as mean ±SD, n = 3 biologically independent samples in **b-l**. *P* values were determined by a two-tailed Student’s *t*-test in **h-k** as indicated in the figures. \*\**P* < 0.01, \*\*\**P* < 0.001, and \*\*\*\**P* < 0.0001.

To further evaluate the biological processes modulated by **ERTLNPs**, Gene Ontology (GO) enrichment analysis was performed on the differentially expressed genes (DEGs) in **ERTLNPs**-treated primary T cells. The results indicated significant enrichment in biological processes related to cell proliferation, protein phosphorylation, cell activation, and metabolic pathways (**Fig. 6f**). Kyoto Encyclopedia of Genes and Genomes (KEGG) pathway analysis revealed that the upregulated genes including Glycolysis/Gluconeogenes, HIF-1 signaling pathway, Amino sugar and nucleotid sugar metabolism, mTOR signaling pathway, and PI3K-Akt signaling pathway, were mainly involved in the mTOR signaling pathway, consistent with our earlier *in vitro* findings. This phenomenon supported the notion that **ERTLNPs** could prime the mTOR pathway to promote T cell activation and metabolic reprogramming, thereby sustaining T cell proliferation, differentiation, and effector functions (**Fig. 6g**). The RNA sequencing analysis thus further confirmed the T cell activation mechanism mediated by **ERTLNPs**.

To validate the RNA sequencing results, 2-(N-(7-nitrobenz-2-oxa-1,3-diazol-4-yl) amino)-2-deoxyglucose (2-NBDG) was employed as a fluorescent glucose analog to assess glucose uptake by T cells following **ERTLNPs** or anti-CD3/CD28 treatment. As shown in **Fig. 6h**, both the **ERTLNPs** and anti-CD3/CD28 groups exhibited significantly enhanced 2-NBDG uptake compared to the PBS group, indicating increased glucose uptake efficiency and elevated metabolic activity in T cells. Furthermore, the level of adenosine triphosphate (ATP) in primary T cells incubated with **ERTLNPs** were significantly increased (**Fig. 6i**), suggesting a heightened state of glucose uptake and energy metabolism upon T cell activation. In addition, the levels of glucose transporter type 1 (**GLUT1)** and hypoxia-inducible factor 1α (HIF-1α) were markedly elevated in the **ERTLNPs**-treated group relative to the PBS group (**Fig. 6j, k**). Notably, upon treatment with mTOR inhibitors such as Rapa and Torin1, the level of GLUT1 was substantially reduced (**Supplementary Fig. 23**). Based on the above results, we found that **ERTLNPs** induced significantly lower expression of T cell exhaustion markers compared to conventional anti-CD3/CD28 Dynabeads stimulation (**Fig. 4i, j**). This observation was further supported by transcriptomic profiling. Heatmap analysis revealed a marked upregulation of key exhaustion-associated genes, including Pdcd1, Ctla4, Lag3, and Batf, in T cells treated with anti-CD3/CD28, whereas such upregulation was absent in **ERTLNPs**-treated cells (**Fig. 6l**). In parallel, **ERTLNPs**-treated T cells exhibited significantly elevated expression of multiple memory-associated genes, including Ccr7, Cd44, Tcf7, Foxo1, Zeb2, and Cxcr3. These genes are known to play critical roles in regulating memory T cell differentiation, homing, and long-term persistence. These findings suggest that **ERTLNPs** promote T cell differentiation toward a memory-like phenotype and support the emergence of a longer-lived, functionally poised T cell subset. (**Fig. 6l**). These results highlight the unique metabolic programming and reduced exhaustion associated with **ERTLNPs**-mediated T cell activation, underscoring its potential as a superior ligand-free activation strategy.

Collectively, these findings suggested that **ERTLNPs**-mediated activation of primary T cells triggered metabolic reprogramming through the mTOR signaling pathway, promoting increased glucose uptake and glycolysis to meet the elevated energy and metabolic demands required for T cell activation and effector function (**Fig. 6m**).

### ERTLNPs generate FAP CAR T cells *in vitro* and *in vivo*

Upon systemic administration, **ERTLNPs** showed efficient T cell transfection and activation in the spleen, highlighting their potential for *in vivo* T cell gene editing such as CAR T cell therapy. Currently, the CAR T cell therapies are predominantly focused on targeting and eliminating cancer cells, the application of *in vivo*–generated CAR T cells to eliminate non-cancerous cell types for the treatment of cancer or other pathological conditions has been rarely reported^18,49,50^. In pathological microenvironments of the diseases such as pancreatic cancer and fibrosis, fibroblasts typically showed elevated surface expression of fibroblast activation protein (FAP). In a recent report, CAR T cells were engineered to target FAP, enabling the selective depletion of fibroblasts in the pathological microenvironment for the treatment of cardiac fibrosis^51,52^. Given the rapid progress of adoptive T cell therapies, targeting FAP offered a promising approach and CAR T cells could be engineered both *in vitro* and *in vivo* to specifically target fibroblasts within diseased tissues (**Fig. 7a**). The components constituting the fibroblast activation protein chimeric antigen receptor (FAP CAR) included anti-mouse FAP single-chain variable fragment (scFv, clone 73.3), CD8 hinge region, and intracellular domains (ICD) of CD28 and CD3ζ. These components were cloned into a plasmid template for *in vitro* transcription (IVT) to generate mRNA encoding the FAP CAR (mFAP CAR, denoted as mFC), which was subsequently used to transfect primary murine T cells (**Fig. 7b**). To evaluate *in vitro* delivery efficiency of mFC, **ERTLNPs** were used to encapsulate mFC and incubated in primary murine T cells for 24 hours. Flow cytometry analysis revealed that approximately 25% of CD3⁺ T cells incubated with **ERTLNPs** were FAP CAR-positive (**Fig. 7c**). We then investigated whether systemic administration of **ERTLNPs** loaded with mFC could efficiently engineer T cells *in vivo* in C57BL/6 mice (**Fig. 7d**). Notably, 9% of FAP CAR^+^ T cells were obtained in the spleen following systemic administration of **ERTLNPs** encapsulting mFC (**Fig. 7e, f**). These findings were consistent with the previous results demonstrating high transfection efficiency of splenic T cells after systemic delivery of **ERTLNPs** encapsulating the reporter gene such as Cre mRNA (**Fig. 3c**). These results collectively indicate that **ERTLNPs** enable efficient *in vivo* T cell engineering to generate FAP CAR T cells upon systemic administration.

**Fig. 7.**
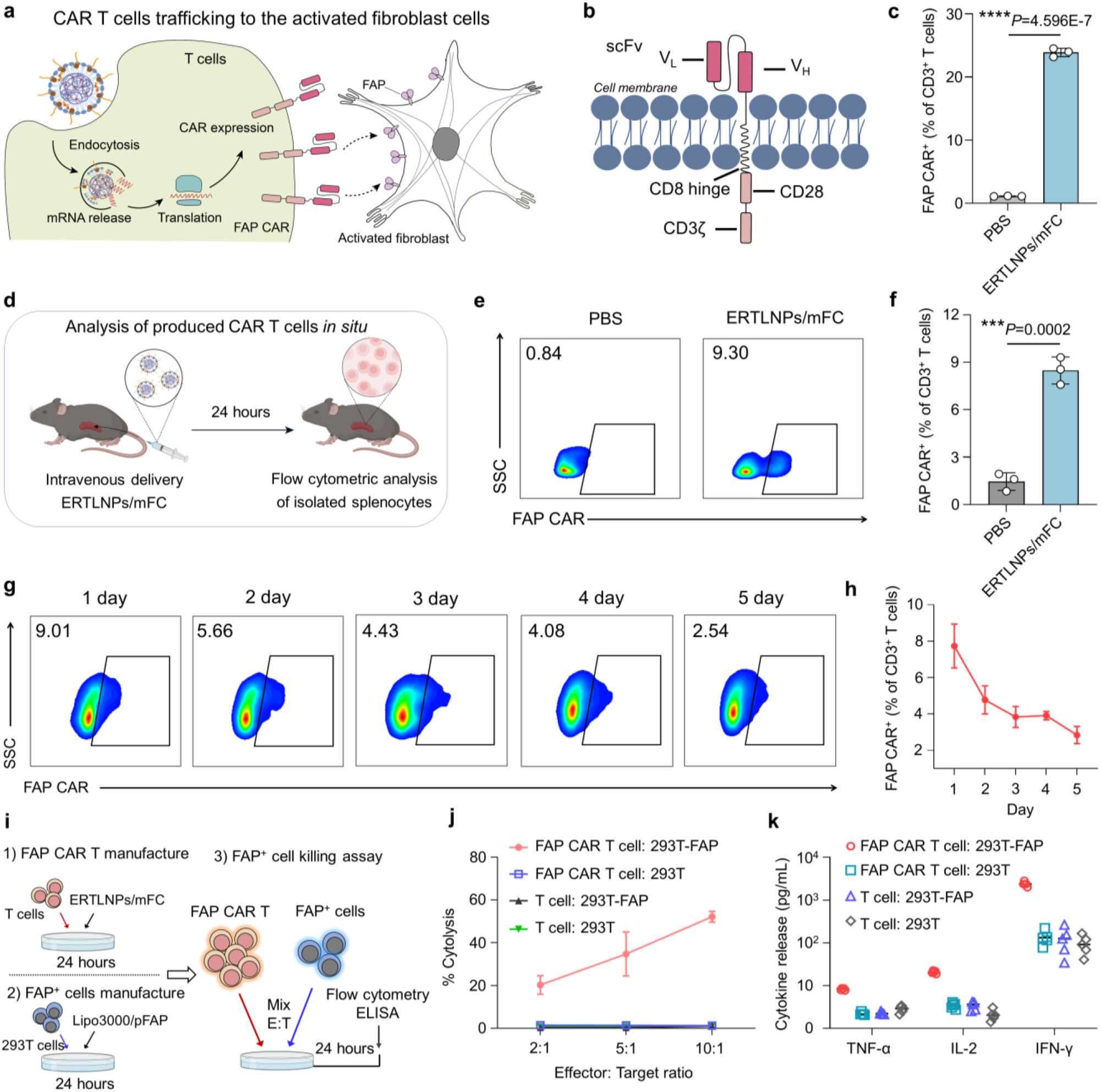
ERTLNPs efficiently mediate the generation of FAP CAR-T cells *in vitro* and *in vivo*. **a**, Illustration of **ERTLNPs/**mFC-transduced T cells trafficking to the activated fibroblast cells. **b**, Schematic of the composition of FAP CAR molecules. **c**, The percentages of FAP CAR^+^ cells in CD3^+^ T cells after **ERTLNPs/**mFC transduction at 24 hours *in vitro*. **d**, Schematic of **ERTLNPs**-mediating mRNA delivery *in vivo* for FAP CAR T cells production. **e**,**f**, Representative flow cytometry analysis (**e**) and relative quantification (**f**) of FAP CAR^+^ cells gating on CD3^+^ T cell in the spleen of C57 mice 24 hours after intravenous administration **ERTLNPs/**mFC (1 mg kg^−1^ mFC). **g**,**h**, Representative flow cytometry analysis. (**g**) and relative quantification of FAP CAR^+^ cells gating on CD3⁺ T in the spleen of C57 mice after systemic administration **ERTLNPs/**mFC at various time points (Day 1, 2, 3, 4, and 5). **i**, Schematic diagram illustrating the *in vitro* generation of FAP CAR T cells and FAP⁺ cells, as well as the cytotoxicity of FAP CAR T cells against FAP⁺ cells. FAP CAR T cell generation: primary T cells transfected with **ERTLNPs**/mFC for 24 hours; FAP^+^ cell generation: 293T cells transfected with lipofectamine 3000/*p*FAP-GFP for 24 hours; Functional validation: FAP CAR T cells were co-cultured with FAP^+^ 293T cells (denoted as 293T-FAP) or parental 293T cells at varying effector-to-target (E:T) ratios (2:1–10:1) for 24 hours, and the supernatants were collected and performed quantitative analysis of cytokines (IL-2, IFN-γ, and TNF-α), and cytotoxicity was assessed by flow cytometry. **j**, Verification of the *in vitro* specific cytotoxicity of FAP CAR T cells against 293T-FAP cells at various effector-to-target (E:T) ratios. **k**, Cytokine analysis (TNF-α, IL-2, and IFN-γ) of the cell supernatant after co-culture of T cells and 293T cells. Data in **c**, **f**, **h**, **j**, and **k** are shown as mean ±SD, n = 3 biologically independent samples in **c**, **f**, and **h**, n = 5 biologically independent samples in **j** and **k.** *P* values were determined by a two-tailed Student’s *t*-test in **c**, **f**, **h**, **j**, and **k** as indicated in the figures. \*\*\**P* < 0.001 and \*\*\*\**P* < 0.0001. A representative image of three independent samples from each group is shown in **e** and **g**.

As **ERTLNPs**-mediated mRNA transfection primarily occurred in the cytoplasm, the expressed FAP CAR gene could not be integrated into the T cell genome. Thus, **ERTLNPs**-mediated *in vivo* production of FAP CAR T cells was transient, with the proportions of FAP CAR⁺ T cells diminishing progressively over time due to T cell proliferation (**Fig. 7g, h**). Given that FAP was also highly expressed during wound healing^52,53^, the presence of persistent FAP CAR T cells might pose safety concerns for patients who develop wounds in the future. Overall, the use of **ERTLNPs** enabled real-time and controllable CAR T cell generation *in vitro* and *in vivo*, providing a highly efficient and safe alternative that circumvented risks associated with permanent genomic integration and long-term CAR expression.

To assess the killing efficacy of **ERTLNPs**-engineered FAP CAR T cells, we established FAP⁺ 293T cells and co-cultured them with FAP CAR T cells alongside FAP⁻ 293T controls for 24 hours (**Fig. 7i**). Our results demonstrated that FAP CAR T cells selectively killed FAP⁺ 293T cells in a dose-dependent manner, whereas their cytotoxicity toward FAP⁻ 293T cells was negligible. Likewise, unmodified T cells exerted minimal killing effects on either FAP⁺ or FAP⁻ 293T cells (**Fig. 7j**). Moreover, the co-culture of FAP CAR T cells with FAP⁺ 293T cells resulted in robust secretion of pro-inflammatory cytokines, including interleukin-2 (IL-2), tumour necrosis factor-α (TNF-α), and interferon-γ (IFN-γ), which was primarily attributed to the selective targeting and lysis of FAP⁺ 293T cells by FAP CAR T cells (**Fig. 7k**)

### ERTLNPs-mediated FAP CAR T cell therapy for idiopathic pulmonary fibrosis

Based on the above results, **ERTLNPs** encapsulating mFC could efficiently generate FAP CAR T cells both *in vitro* and *in vivo* for the clearance of FAP^+^ cells. According to RNA sequencing results derived from patient samples, the levels of FAP were significantly upregulated in fibrotic lung tissues compared to normal lung tissues (**Fig. 8a**), indicating that FAP was a viable target for **ERTLNPs**-mediated *in vivo* generation of FAP CAR T cells for the treatment of pulmonary fibrosis. To evaluate the therapeutic effect of *in vivo*–generated FAP CAR T cells by **ERTLNPs** on pulmonary fibrosis, we established a bleomycin-induced idiopathic pulmonary fibrosis (IPF) mouse model. Fluorescence images of lung tissue sections showed that fibroblasts overexpressed FAP, whereas FAP expression was nearly undetectable in healthy lung tissues (**Fig. 8b**). Moreover, FAP CAR T cells generated in the spleen could effectively infiltrate the fibrotic lung lesions and interacted with FAP^+^ pulmonary fibroblasts. This phenomenon confirmed that FAP CAR T cells were capable of migrating specifically from the lymphoid reservoir to the fibrotic microenvironment and targeting FAP^+^ fibroblasts (**Fig. 8b**).

**Fig. 8.**
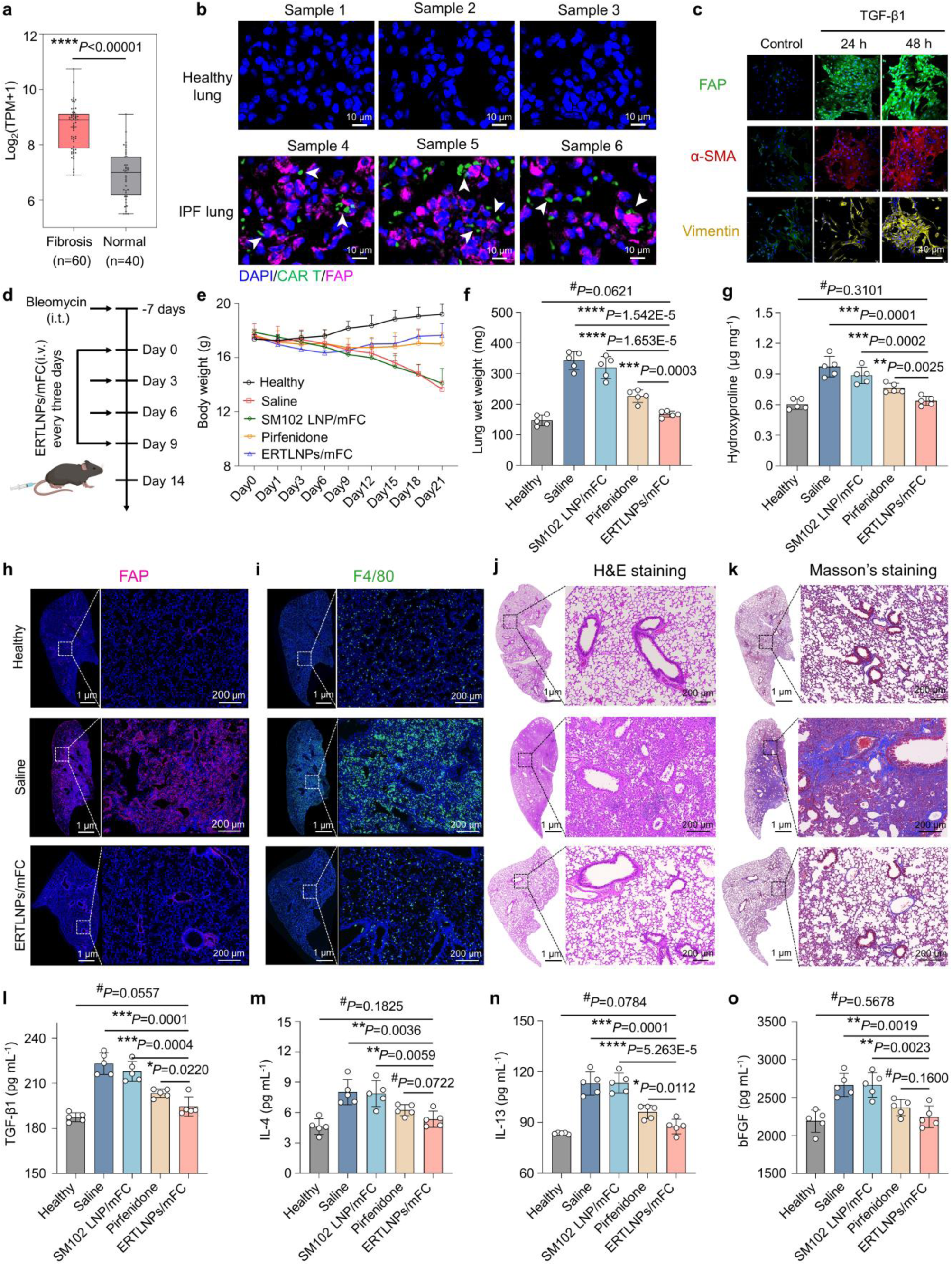
ERTLNPs-mediated *in vivo* generation of FAP CAR T cells efficiently alleviates the progression of idiopathic pulmonary fibrosis (IPF). **a**, Relative levels of FAP expression in lung tissues of IPF patients (fibrosis group, n = 60) and healthy individuals (normal group, n = 40).**b**, Representative immunofluorescence images of FAP^+^ cells (red) and FAP CAR T (green) in the lung tissues of IPF and healthy mice. The white arrow indicated FAP CAR T cells attacking FAP⁺ cells. **c**, Representative fluorescence images of FAP, α-smooth muscle actin (α-SMA) and vimentin expression in primary murine lung fibroblasts treated with TGF-β1 (10 ng mL^−1^) for 24 hours or 48 hours). Scale bar: 40 μm. **d**, Schematic of **ERTLNPs/**mFC treatment in a mouse model of IPF. **e**, Average body weight changes of C57 mice with pulmonary fibrosis receiving different treatments. **f**, Lung wet weight and hydroxyproline levels of C57 mice with pulmonary fibrosis after treatment. Pirfenidone (25 mg kg^−1^, oral gavage, every other day) was used as positive control. **h**,**i**, Representative immunofluorescence images of FAP^+^ cells (**h**) and F4/80^+^ cells (**i**) in lung tissues of C57 mice with pulmonary fibrosis after treatment. Age- and weight-matched healthy mice were used control. **j**,**k**, Representative H&E (**j**) and Masson’s trichrome staining images (**k**) in lung tissues of C57 mice with pulmonary fibrosis after treatment. Age- and weight-matched healthy mice were used control. **l-o**, Levels of cytokines including TGF-β1 (**l**), IL-4 (**m**), IL-13 (**n**), and bFGF (**o**) in lung tissues of C57 mice with pulmonary fibrosis after treatment. Age- and weight-matched healthy mice were used control. Data in **e-g** and **l-o** are shown as mean ± SD, n = 5 biologically independent samples in **e-g** and **l-o**. *P* values were determined by a two-tailed Student’s *t*-test in **a**, **f**, **g**, and **l-o** as indicated in the figures. *^#^P* > 0.05, \**P* < 0.05, \*\**P* < 0.01, \*\*\**P* < 0.001, and \*\*\*\**P* < 0.0001. A representative image of three independent samples from each group is shown in **b**, **c**, and **h-k**.

In addition, transforming growth factor β1 (TGF-β1) was typically overexpressed in fibrotic tissues^54^. Thus, we isolated primary lung fibroblasts from healthy mice and incubated them with TGF-β1 for 24 and 48 hours. In both conditions, the level of FAP was markedly improved, indicating that TGF-β1 could induce a transition from the quiescent phenotype to fibrotic phenotype in lung fibroblasts (**Fig. 8c**). In contrast, FAP was nearly undetectable in fibroblasts cultured without TGF-β1. In addition, fibrosis markers such as α-smooth muscle actin (α-SMA) and vimentin were significantly upregulated in TGF-β1-treated fibroblasts (**Fig. 8c**). These data collectively demonstrated that FAP was a reliable marker of fibroblasts in fibrotic lung tissue, which encouraged us to further investigate whether **ERTLNPs**-mediated *in vivo* generation of FAP CAR T cells could be directly used to treat IPF in mice.

Therefore, the bleomycin-induced IPF mouse model was established for preclinical evaluation of *in vivo*-generated FAP CAR T cells by **ERTLNPs** (**Fig. 8d**). Pirfenidone, a clinically approved antifibrotic drug, were selected as a positive control. Given that *in vivo* generation of FAP CAR T cells *via* **ERTLNPs** was a transient process, **ERTLNPs/**mFC were administered intravenously once every three days, for a total of four doses. In the IPF mouse model, the saline and SM102-LNP groups exhibited significant body weight loss during the treatment. In contrast, the pirfenidone and **ERTLNPs/**mFC groups showed negligible body weight loss, indicating that **ERTLNPs/**mFC therapy effectively alleviated the progression of IPF and exhibited good *in vivo* safety (**Fig. 8e**). At the end of the treatment, lung tissues were collected and weighed. Compared to healthy mice, the lung wet weights in the saline and SM102-LNP/mFC groups increased significantly, mainly due to congestion and edema triggered by interstitial inflammation during IPF progression (**Fig. 8f**). In contrast, the lung wet weights in the **ERTLNPs/**mFC group showed negligible difference from healthy mice and were comparable to that of the pirfenidone-treated group. Additionally, the hydroxyproline contents in the lung tissue of the **ERTLNPs/**mFC group were much lower than those of the saline and SM102 LNP/mFC groups, and comparable to that of healthy mice (**Fig. 8g**). These results strongly suggested that **ERTLNPs/**mFC treatment could substantially mitigate the progression of idiopathic pulmonary fibrosis in mice.

Subsequently, we evaluated FAP expression in lung tissues of IPF mice after treatment. It was observed that following **ERTLNPs/**mFC treatment, the proportion of FAP⁺ cells in the lung tissues of IPF mice was drastically reduced, reaching levels comparable to those in healthy mice (**Fig. 8h**). In contrast, large numbers of FAP⁺ cells (i.e., activated fibroblasts) were observed in the lung tissues of the saline and SM102 LNP/mFC groups (**Supplementary Fig. 24**). Moreover, a large number of macrophages were present in the lung tissue of the saline-treated group (**Fig. 8i, and Supplementary Fig. 25**). Macrophages were critically involved in the exacerbation of idiopathic pulmonary fibrosis, as they recruited other inflammatory effector cells, such as neutrophils, and contributed to pulmonary tissue damage^55,56^. By comparison, the proportion of macrophages in the lung tissues of the **ERTLNPs/**mFC-treated group was significantly downregulated and was comparable to that of healthy mice. These results indicated that **ERTLNPs/**mFC treatment effectively eliminated activated fibroblasts and macrophages in fibrotic lung tissues. Furthermore, the H&E and Masson’s trichrome staining images showed that compared with healthy mice, lung tissues of IPF mice displayed extensive collagen deposition and severely disrupted alveolar structures (**Fig. 8j, k** and **Supplementary Figs. 26, 27**). Notably, **ERTLNPs/**mFC treatment effectively cleared collagen and extracellular matrix from the lung tissue, while leading to a marked restoration of alveolar structure and function. Moreover, TGF-β1 of lung tissues, as a key profibrotic cytokine, was significantly downregulated after **ERTLNPs/**mFC treatment (**Fig. 8l**). IL-4 and IL-13 of lung tissues which drove Th2 immune responses and exacerbated fibrosis were also reduced in the **ERTLNPs/**mFC group (**Fig. 8m, n**). In addition, the level of basic fibroblast growth factor (bFGF) contributing to abnormal angiogenesis and fibrotic tissue expansion was decreased in lung tissues of the **ERTLNPs/**mFC group (**Fig. 8o**). Furthermore, liver function markers including aspartate aminotransferase (AST), alanine aminotransferase (ALT), alkaline phosphatase (ALP) and kidney function markers including creatinine (CRE), blood urea nitrogen (BUN), uric acid (UA) were also measured after treatment (**Supplementary Fig. 28, 29**). The saline, pirfenidone, and SM102 LNP groups showed clear signs of liver dysfunction, and the pirfenidone group also showed elevated serum creatinine, suggesting systemic toxicity. In contrast, liver and kidney function markers in the **ERTLNPs/**mFC-treated group were comparable to those in healthy mice. H&E staining images of major organs (heart, liver, spleen, kidneys) also revealed normal histopathological morphology in the **ERTLNPs/**mFC group (**Supplementary Fig. 30**).

These findings collectively demonstrated that systemically administered **ERTLNPs/**mFC could generate FAP CAR T cells *in vivo*, which selectively target and eliminate the activated fibroblasts in the lungs of IPF mice. The **ERTLNPs/**mFC platform contributed to significant therapeutic improvement without causing the systemic side effects observed with pirfenidone or SM102 LNPs.

### ERTLNPs-mediated *in vivo* CAR T cell therapy for liver fibrosis

In normal liver tissue, hepatic stellate cells (HSCs) remained in a quiescent state. However, in chronically injured liver tissue, these cells became activated into myofibroblasts and produced large amounts of collagen and extracellular matrix (ECM)^57^. In addition, Kupffer cells, the liver-resident macrophages, were activated in chronic liver injury and secreted inflammatory cytokines (e.g., IFN-α, IL-1β) and TGF-β1^58^, while recruiting other immune cells to participate in the inflammatory response, ultimately inducing the formation of a fibrotic liver microenvironment. Recent studies have reported that in fibrotic liver tissues, the activated fibroblasts overexpressing FAP contributed to liver fibrosis progression by activating HSCs and modulating the phenotype of Kupffer cells^59^. Thus, activated fibroblasts represented a viable therapeutic target for liver fibrosis, and the specific elimination of FAP^+^ fibroblasts could be considered as an effective strategy to alleviate inflammation and fibrosis progression in the liver tissues. Here, we established a carbon tetrachloride (CCl₄)-induced liver fibrosis model (**Fig. 9a**) and evaluated the therapeutic efficacy of **ERTLNPs/**mFC following systemic administration. Magnesium isoglycyrrhizinate (MgIG), a clinically used antifibrotic agent, was used as a positive control. As fibrosis progressed, liver tissues accumulated a large amount of collagen and ECM, resulting in significantly increased liver wet weight (**Fig. 9b**). Compared with the saline and MgIG groups, mice treated with **ERTLNPs/**mFC showed significantly reduced liver wet weight, approaching that of normal liver tissues (**Fig. 9b**). Moreover, the hydroxyproline content in the livers of mice treated with **ERTLNPs/**mFC was 2.2-fold lower than that of the saline group (**Fig. 9c**). Masson’s trichrome staining images further revealed that fibrotic areas (blue regions) in the **ERTLNPs/**mFC -treated group were markedly smaller than those in the saline and MgIG groups (**Fig. 9d**). H&E staining images showed that the liver tissues in the saline and MgIG groups exhibited disorganized cellular architecture and severely disrupted hepatic sinusoidal structures, whereas the liver tissues in the **ERTLNPs/**mFC group exhibited regular morphology and compact cellular arrangement (**Fig. 9d**).

**Fig. 9.**
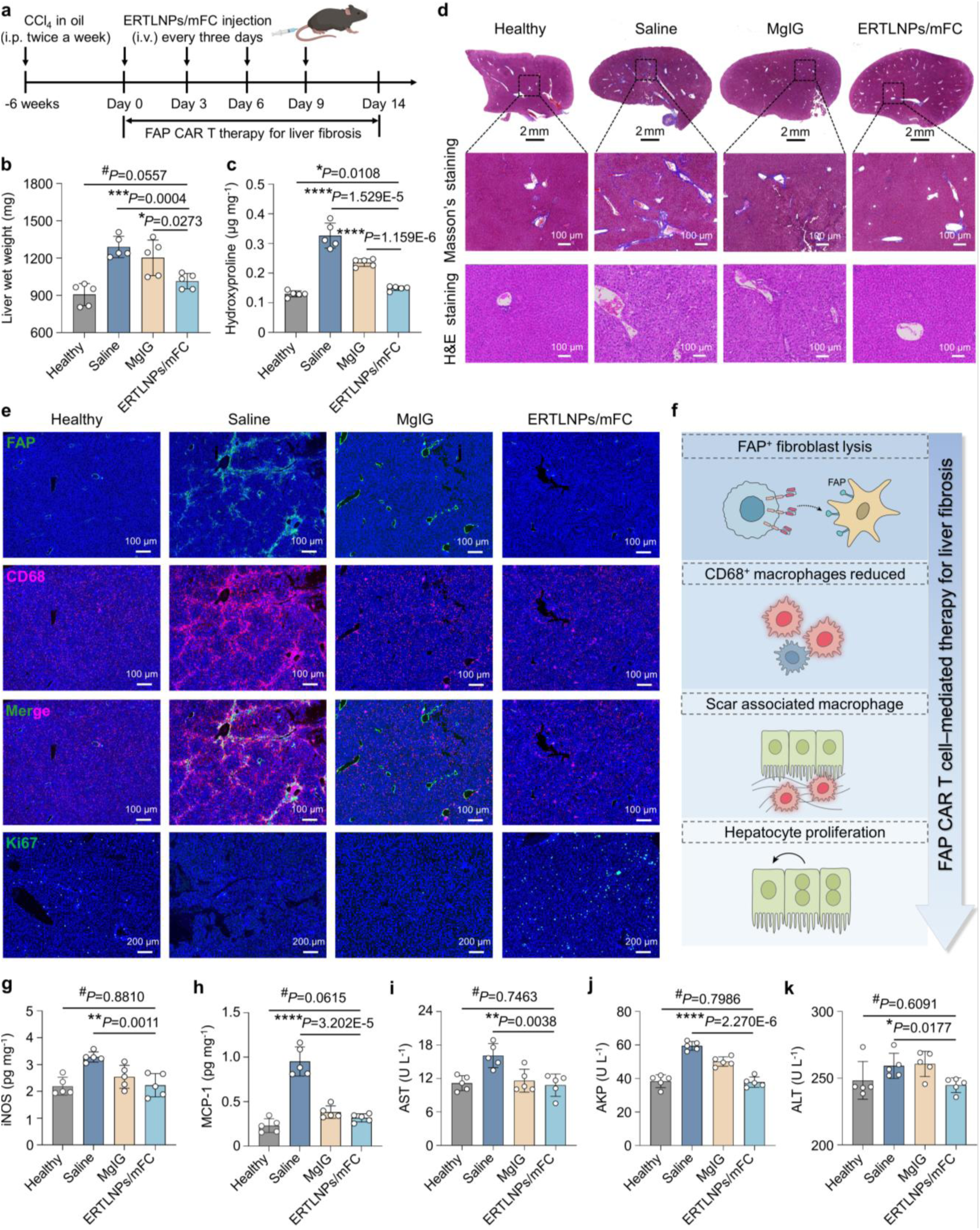
ERTLNPs-mediated *in vivo* generation of FAP CAR T cells significantly relieves liver fibrosis. **a**, Schematic of **ERTLNPs/**mFC treatment in a mouse model of liver fibrosis. **b**,**c**, Liver wet weight (**b**) and hydroxyproline content (**c**) in different treatments were analyzed. Magnesium isoglycyrrhizinate (MgIG) (50 mg kg^−1^, intraperitoneal injection, once every two day) was used as positive control. **d**, Representative Masson’s trichrome and H&E staining images of liver tissues in a mouse model of liver fibrosis after treatment. Age- and weight-matched healthy mice were used control. **e**, Representative immunofluorescence images of FAP^+^, CD68^+^, and Ki67^+^ cells in liver tissues of C57 mice with liver fibrosis after treatment. **f**, Schematic of the action mechanism of FAP CAR T cells in alleviating liver fibrosis progression. **g**,**h**, Levels of iNOS (**g**) and MCP-1 (**h**)in liver tissues of C57 mice with liver fibrosis after treatment. **i-k**, Levels of liver function markers including aspartate aminotransferase (AST) (**i**), alkaline phosphatase (AKP) (**j**), and alanine aminotransferase (ALT) (**k**) in liver tissues of C57 mice with liver fibrosis after treatment. Data in **b**, **c** and **g-k** are shown as mean ± SD, n = 5 biologically independent samples in **b**, **c** and **g-k**. *P* values were determined by a two-tailed Student’s *t*-test in **b**, **c** and **g-k** as indicated in the figures. *^#^P* > 0.05, \**P* < 0.05, \*\**P* < 0.01, \*\*\**P* < 0.001, and \*\*\*\**P* < 0.0001. A representative image of three independent samples from each group is shown in **d** and **e**.

Immunofluorescence images showed that, the liver tissues of the saline-treated group exhibited a significantly higher percentages of FAP⁺ fibroblasts and macrophages than those of healthy mice (**Fig. 9e**). Macrophage infiltration was closely linked to chronic inflammation and liver fibrosis severity^59^. In normal livers, CD68⁺ macrophages were sparsely distributed, whereas their infiltration significantly increased in the fibrous septa of fibrotic livers, with a subset of CD68⁺ macrophages closely colocalized with FAP⁺ fibroblasts (**Fig. 9e**). This spatial interaction may further stimulate CD68⁺ macrophages to secrete inflammatory mediators (e.g., TNF-α, IL-1β, and TGF-β1), thereby exacerbating fibrosis. In addition, CD68⁺ macrophages contributed to tissue repair and scar formation within fibrotic areas, potentially promoting fibrosis progression through crosstalk with fibroblasts^60^.

Compared to the saline group, the **ERTLNPs/**mFC group exhibited a significant reduction in both FAP⁺ fibroblasts and CD68⁺ macrophages. Notably, Ki67, a proliferation marker, was markedly upregulated after **ERTLNPs/**mFC treatment, suggesting that **ERTLNPs**-mediated *in vivo* generation of FAP CAR T cells effectively eliminated FAP⁺ fibroblasts and promoted hepatocyte proliferation and liver tissue regeneration (**Fig. 9e, f**). Furthermore, the levels of inducible nitric oxide synthase (iNOS, a marker of M1φ macrophages), monocyte chemoattractant protein-1 (MCP-1), arginase-1 (Arg1, a marker of M2φ macrophages), and bFGF were significantly downregulated in the liver tissues after **ERTLNPs/**mFC treatment (**Fig. 9g, h, and Supplementary Fig. 31, 32**). Moreover, the levels of liver function markers including AST, ALT, ALP were elevated in saline-treated fibrotic mice compared to Age- and weight-matched healthy mice. In contrast, liver function values in the **ERTLNPs/**mFC group were within the normal range (**Fig. 9i–k**). Moreover, the major organs (heart, spleen, lung, kidney) displayed normal histological morphology after treatment (**Supplementary Fig. 33**), supporting the *in vivo* safety of **ERTLNPs**-mediated CAR T cell therapy.

Taken together, **ERTLNPs**-mediated *in vivo* CAR T cell therapy effectively eliminated activated fibroblasts, macrophages, and excess ECM of fibrotic liver tissues. The **ERTLNPs/**mFC treatment significantly reversed hepatic inflammation, promoted hepatocyte proliferation and regeneration, recovered to normal liver function, and substantially alleviated disease progression in the liver fibrosis mouse model.

### ERTLNPs-mediated *in vivo* CAR T cell therapy for orthotopic pancreatic cancer

Pancreatic cancer is one of the most lethal solid tumours despite the use of multi-agent conventional chemotherapy regimens^61,62^. This poor prognosis was mainly due to two factors: First, pancreatic tumour tissues contained a high abundance of activated fibroblasts (FAP^+^ fibroblasts), which promoted tissue fibrosis, hindering drug penetration and reducing efficacy against cancer cells. Second, the fibrotic and immunosuppressive microenvironment impaired the infiltration and functionality of cytotoxic T lymphocytes (CTLs), further accelerating disease progression^63,64^. A desirable therapeutic strategy involved the *in vivo* generation of FAP CAR T cells to specifically target and eliminate activated fibroblasts, thereby alleviating pancreatic tumour fibrosis. Simultaneously, this approach could be combined with immune checkpoint blockade to enhance the infiltration and function of CTLs. Therefore, we proposed to utilize **ERTLNPs** to mediate the *in vivo* generation of FAP CAR T cells for the specific targeting and elimination of fibroblasts, aiming to alleviate pancreatic tumour fibrosis and enhance CTLs infiltration. Meanwhile, such strategy was combined with the immune checkpoint inhibitor anti–programmed cell death protein 1 (anti–PD-1) treatment to relieve the immunosuppressive effects of cancer cells on T cells and enhance their cytotoxic function.

By analyzing the gene expression in pancreatic ductal adenocarcinoma (PDAC) and normal pancreatic tissues by The Cancer Genome Atlas (TCGA), it was observed that levels of FAP in tumour tissues from pancreatic cancer patients were significantly higher than those in normal pancreatic tissue samples (**Fig. 10a**). We then established an orthotopic pancreatic cancer mouse model and evaluated the therapeutic efficacy of *in vivo*-generated FAP CAR T cell therapy *via* **ERTLNPs/**mFC in combination with anti–PD-1 (Ab) treatment (**Fig. 10b**). The first-line clinical drug gemcitabine (GEM) and Ab were used as controls. Compared to the saline group, both Ab and GEM showed only modest therapeutic effects, and tumour overgrowth during the treatment period leaded to the death of some mice in these groups (**Fig. 10c, d**). In contrast, the **ERTLNPs/**mFC group could efficiently suppress tumour growth, and the **ERTLNPs/**mFC plus Ab group achieved the most significant anti-tumour efficacy. Tumour images and tumour weight data also demonstrated that the **ERTLNPs/**mFC plus Ab group had the most effective tumour suppression (**Fig. 10e, f**). Moreover, the **ERTLNPs/**mFC plus Ab treatment significantly prolonged the survival of pancreatic cancer-bearing mice, with no deaths observed even sixty days after the combination therapy (**Fig. 10g**). Moreover, the tumour-bearing mice did not experience any body weight loss during the treatment period, indicating that the combination of **ERTLNPs/**mFC plus Ab demonstrated good *in vivo* safety (**Supplementary Fig. 34**). These results sufficiently demonstrated that **ERTLNPs**-mediated *in vivo* CAR T cell therapy significantly enhanced the efficacy of anti-PD-1–mediated immunotherapy against pancreatic cancer, markedly prolonged the survival of pancreatic tumour–bearing mice, and exhibitd superior therapeutic outcomes compared to the first-line chemotherapeutic drug GEM.

**Fig. 10.**
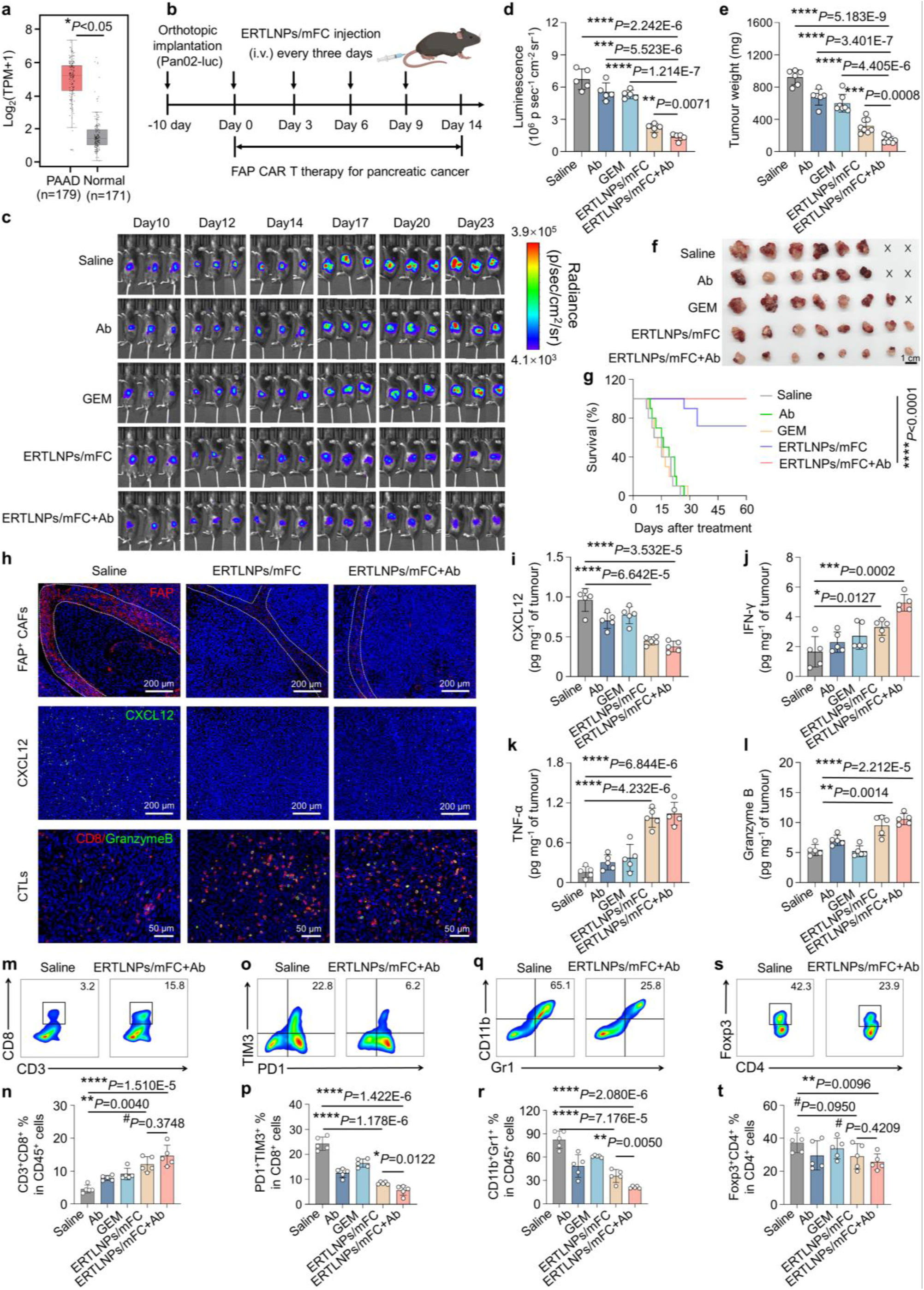
ERTLNPs-mediated *in vivo* generation of FAP CAR T cells significantly suppresses the growth of orthotopic pancreatic cancer in mice. **a**, Relative levels of FAP expression in tumor tissues (n=179) and adjacent tissues (n = 171) of pancreatic cancer patients. PAAD: pancreatic adenocarcinoma. Quantitative analysis of FAP expression obtained from the TCGA database. **b**, Schematic of **ERTLNPs**/mFC treatment in a mouse model of orthotopic pancreatic cancer. **c,** Representative *in vivo* bioluminescence images of Pan02 tumor-bearing mice receiving different treatments. **d**, Bioluminescence intensity of tumor tissues of C57 mice with orthotopic pancreatic cancer after treatment. **e**,**f**, Tumour weights (**e**) and tumour images (**f**) of C57 mice with orthotopic pancreatic cancer after treatment. **g**, Survival curves of Pan02 tumor-bearing mice receiving different treatments. **h**, Representative immunofluorescence images of FAP^+^ cells (red), CXCL12 expression (green), and activated cytotoxic T cells (Granzyme B^+^/CD8^+^, orange) in the tumour tissues of C57 mice with orthotopic pancreatic cancer after treatment with **ERTLNPs**/mFC or **ERTLNPs**/mFC plus Ab. The area outlined by the white dashed line indicates the fibroblast-rich region within the tumour tissues of orthotopic pancreatic cancer in mice. Scale bar: 200 μm or 50 μm. **i**-**l**, Levels of CXCL12 (**i**), IFN-γ (**j**), TNF-α (**k**), and Granzyme B (**l**) in tumour tissues of C57 mice with orthotopic pancreatic cancer after treatment. **m,n**, Representative flow cytometric analysis (**m**) and relative quantification (**n**) of CD3^+^CD8^+^ T cells gating on CD45^+^ cells in the tumours of C57 mice with orthotopic pancreatic cancer at the end of treatment. **o,p,** Representative flow cytometric analysis (**o**) and relative quantification (**p**) of exhausted T cells (TIM-3^+^PD1^+^CD8^+^ T) gating on CD45^+^ cells in the tumours of C57 mice with orthotopic pancreatic cancer at the end of treatment. **q,r,** Representative flow cytometric analysis (**q**) and relative quantification (**r**) of MDSCs (CD11b^+^ Gr-1^+^cells gating on CD45^+^ cells) in the tumours of C57 mice with orthotopic pancreatic cancer at the end of treatment. **s,t,** Representative flow cytometric analysis (**s**) and relative quantification (**t**) of Tregs (Foxp3^+^CD4^+^) gating on CD45^+^ CD3^+^ cells in the tumours of C57 mice with orthotopic pancreatic cancer at the end of treatment. Data in **d**, **e**, **i-l, n, p, s,** and **t** are shown as mean ±SD, n = 5 biologically independent samples in **c-e**, **i-l**, **n**, **p**, **r**, and **t**, n = 8 biologically independent samples in **f** and **g**. *P* values were determined by a two-tailed Student’s *t*-test in **a**, **d**, **e**, **g**, **i-l**, **n**, **p**, **s**, and **t** as indicated in the figures. *^#^P* > 0.05, \**P* < 0.05, \*\**P* < 0.01, \*\*\**P* < 0.001, and \*\*\*\**P* < 0.0001. A representative image of three independent samples from each group are shown in **h**. A representative image of five independent samples from each group are shown in **m**, **o**, **q**, and **s.**

Furthermore, immunofluorescence images revealed that both the **ERTLNPs/**mFC and **ERTLNPs/**mFC plus Ab groups had significantly fewer FAP^+^ cells than the saline group in tumour tissues, and the fibroblast-enriched stroma around cancer cells was nearly absent in these treatment groups (**Fig. 10h and Supplementary Fig. 35**). Moreover, CXCL12 secreted by FAP⁺ fibroblasts could bind to CXCR4 on cancer cells or T cells, activating the CXCL12–CXCR4 signaling axis, which promoted tumour stromal fibrosis and triggered the downstream PI3K/Akt pathway, thereby promote cancer cell proliferation and invasion, while suppressing the cytotoxic function of CTLs^63^. Notably, **ERTLNPs/**mFC treatment significantly suppressed CXCL12 expression in tumour tissues (**Fig. 10h, i**). In addition, following treatment with **ERTLNPs/**mFC and **ERTLNPs/**mFC plus Ab, there was a markedly increased infiltration of CD8⁺ T cells and Granzyme B⁺ CD8⁺ T cells (**Fig. 10h, m, and Supplementary Fig. 36, 37**). Moreover, following **ERTLNPs/**mFC treatment, pro-inflammatory cytokines (IFN-γ, TNF-α, and Granzyme B) in the tumour tissue were significantly upregulated (**Fig. 10j-l**), while anti-inflammatory cytokines (TGF-β1 and IL-10) were markedly downregulated (**Supplementary Fig. 38, 39**), and the proportions of exhausted T cells (PD1^+^ TIM3^+^ T cells) and immunosuppressive cells such as myeloid-derived suppressor cells (MDSCs) and regulatory T cells (Tregs) were significantly reduced. (**Fig. 10m-t, and Supplementary Fig. 40-45**). These findings indicated that **ERTLNPs**-mediated CAR T cell therapy effectively eliminated activated fibroblasts within pancreatic tumour tissues, alleviated tumour-associated fibrosis, significantly enhanced both the infiltration and function of CTLs cells, and efficiently reversed the immunosuppressive tumour microenvironment of pancreatic cancer.

In addition, pancreatic cancer usually disrupted the metabolic functions of major organs such as liver and kidney during its progression^65^. The serum liver function makers (AST, ALT, ALP) were significantly elevated in the saline, Ab, and GEM groups compared to healthy controls, with increased ALP value suggesting abnormal bile acid metabolism^66^. Likewise, the renal function markers (UA, BUN, CRE) also showed significant abnormalities in the saline, Ab, and GEM groups compared to healthy mice. Among them, UA serves as a parameter of oxidative stress^67^, indicating enhanced oxidative stress *in vivo* (**Supplementary Fig. 46**, **47**). H&E staining images showed disorganized splenic germinal centers and fibrotic lesions in lung tissues in the saline and GEM groups. Furthermore, mice in the saline group exhibited obvious lung metastases (**Supplementary Fig. 48**). In contrast, **ERTLNPs/**mFC treatment restored the metabolic functions of major organs in pancreatic cancer-bearing mice and effectively inhibited the lung metastasis of pancreatic cancer cells.

## Discussion

In this work, we designed and developed an inherent T cell-activating polymer–lipid nanoparticle (**ERTLNPs**) system for *in vivo* CAR T cell therapy in various diseases. Upon systemic administration, **ERTLNPs** exhibited selective and efficient T cell transfection specifically in the spleen. The **ERTLNPs** rapidly and efficiently activated primary T cells through the mTOR pathway *via* two distinct mechanisms: (1) interactions between **ERTLNPs** and IGF-1R receptors on T cells, and (2) interactions between arginine residues of internalized **ERTLNPs** and transmembrane protein TM4SF5 within lysosomes. Taken together, these mechanisms synergistically triggered downstream mTOR signaling pathway, ultimately promoting primary T cell activation. Notably, such T cell-activating capability of **ERTLNPs** also facilitated enhanced mRNA transfection efficiency in primary T cells. *In vitro* RNA sequencing further revealed that **ERTLNPs** reprogrammed the metabolism of primary T cells and upregulated mTOR signaling, thereby contributing to T cell activation. Given that activated fibroblasts overexpressing FAP were implicated in disease progression in various pathological tissues, including pancreatic cancer and fibrotic lesions, mRNA encoding FAP-specific chimeric antigen receptor (mFAP CAR) was selected as the therapeutic gene. Both *in vitro* and *in vivo* studies confirmed that **ERTLNPs** loaded with mFAP CAR could efficiently generate functional FAP CAR T cells with potent cytotoxic activity. Moreover, we established various disease models including pulmonary fibrosis, hepatic fibrosis, and orthotopic pancreatic cancer. In all cases, **ERTLNPs/**mFC treatment efficiently eliminated activated fibroblasts within the diseased tissue, effectively reshaped the diseased tissue microenvironment and achieved superior therapeutic outcomes compared to frontline clinical drugs, without inducing systemic immune-related adverse events. Given the potent T cell-activating capability of **ERTLNPs**, potential concerns regarding systemic inflammatory responses were carefully further evaluated. To assess cytokine-associated toxicity, the serum levels of pro-inflammatory cytokines including interleukin-6 (IL-6), granulocyte-macrophage colony-stimulating factor (GM-CSF), IL-2, IFN-γ, and TNF-α were monitored 24 and 72 hours following systemic administration of **ERTLNPs**. The levels of these cytokines of **ERTLNPs-**treated mice remained comparable to those in healthy mice, with no significant elevation observed at either time point (**Supplementary Fig. 49**). These results indicate that **ERTLNPs** effectively avoid excessive cytokine release and mitigate the risk of cytokine release syndrome (CRS), supporting their potential as a safe and well-tolerated mRNA delivery platform for *in vivo* T cell engineering. In future work, we will collaborate closely with clinicians to thoroughly evaluate the potential of **ERTLNPs** loaded with mFAP CAR formulations for clinical trials. Additionally, we will further investigate target cell markers for various diseases and construct corresponding CAR mRNAs to be delivered *via* **ERTLNPs** for both *in vitro* and *in vivo* CAR T cell therapies. Furthermore, we also plan to utilize artificial intelligence (AI) and other technologies to optimize carrier material design, exploring nanocarriers that can specifically target and activate other immune cell types such as macrophages or natural killer (NK) cells for the treatment of cancers, autoimmune diseases, and other conditions. In conclusion, the **ERTLNPs** platform represents a proof-of-concept for an inherent T cell-activating nanoparticle-based mRNA delivery system, offering a possible avenue for *in vivo* CAR T cell therapy in various diseases.

## Supporting information

All data in this study are available within the article and its supplementary information.

## Methods

Polyethylenimine (PEI) was purchased from Aladdin (Shanghai, China). Boc-Orn(Boc)-OH, Boc-Orn(Tos)-OH, Boc-Arg(Pbf)-OH, and Boc-Arg(Tos)-OH were purchased from GL Biochem. Ltd (Shanghai, China). Boc-Arg(NO_2_)-OH was purchased from Enlai Biological Technology Co., Ltd (Chengdu, China). Trifluoroacetic acid (TFA) was obtained from Xiya Reagent Co., Ltd. N, N-dimethyl formamide (DMF) was obtained from Beijing Chemical Factory. *p*-Toluene sulfonyl chloride, diisopropylethylamine (DIPEA), and 1-hydroxy benzotriazoleanhydrous (HOBT) were obtained from Dibai Chemical Technology Co., Ltd (Shanghai, China). EDC hydrochloride (EDCI) was obtained from J&K Scientific Ltd (Beijing, China). N-hydroxy succinimide (NHS) was obtained from TCI Co., Ltd (Shanghai, China). 2-Dimyristoyl-rac-glycero-3-methoxypolyethylene glycol-2000 (DMG-PEG), SM-102 lipid, cholesterol, and 1,2-distearoyl-sn-glycero-3-phosphocholine (DSPC) were purchased from Pharmaceutical Tech. Co., Ltd. (Shanghai, China). Bleomycin sulfate, CCl_4_, and corn oil were purchased from Aladdin (Shanghai, China). Anti-FAP antibody, anti-α-SMA antibody, and anti-vimentin antibody were purchased from Abcam (UK). DAPI was purchased from Sigma (USA). Blood biochemical assay kits were purchased from Jiancheng Biotechnology Co., Ltd (Nanjing, China). All enzyme-linked immunosorbent assay (ELISA) kits were purchased from Invitrogen. The inhibitors were purchased from MedChemexpress (USA). anti-CD3/CD28 Dynabeads were purchased from Thermo Fisher Scientific (USA). The fluorescence-labelled antibodies used for flow cytometry analysis were purchased from Cell Signaling Technology (USA) or Biolegend (USA).

## Statistical analysis

All statistical analyses were obtained using GraphPad Prism (Prism 9.0). All experiments in this work were repeated at least three times (n ≥ 3) to ensure their reliability and accuracy. All data are expressed as mean ±standard deviation (mean ±SD). Two-tailed Student’s t-test was used for two-group comparisons, and one-way analysis of variance (ANOVA) with Tukey’s test were used for multiple comparisons. Thresholds for statistically significant differences were \**P* < 0.05, \*\**P* < 0.01, \*\*\**P* <0.001, and \*\*\*\**P* < 0.0001. Threshold for no significant difference was ^#^*P >* 0.05. All graphic illustrations were created by BioRender.com. Flow cytometry data were analyzed by FlowJo software (v10.8.1).

## Reporting summary

Further information on research design is available in the Nature Portfolio Reporting Summary linked to this article.

## Data availability

All data in this study are available within the article and its supplementary information. The raw and analyzed datasets generated in the study are available for research purposes from the corresponding author on request. Source data are provided with this paper.

## Acknowledgements

This work was acknowledged to National Natural Science Foundation of China (52433006, 52473150, 52495010, 52203183, 51925305), Natural Science Foundation of Xiamen, China (3502Z202371004), National Key Research and Development Program of China (2021YFB3800900), Shenzhen Science and Technology Program (JCYJ20240813145515020), Fundamental Research Funds for the Central Universities (20720230004), the talent cultivation project Funds for the Innovation Laboratory for Sciences and Technologies of Energy Materials of Fujian Province (HRTP-[2022] 52), National University of Singapore (NUHSRO/2020/133/Startup/08, NUHSRO/2023/008/NUSMed/TCE/LOA, NUHSRO/2021/034/TRP/09/Nanomedicine, 23-0173-A0001), National Medical Research Council (MOH-001388-00, CG21APR1005, MOH-001500-00, MOH-001609-00, MOH-001740-01), Singapore Ministry of Education (MOE-000387-00, MOE-MOET32023-004), and National Research Foundation (NRF-000352-00).

## Author contributions

Q.-N.C., X.-Y.C., and H.-P.F., H.-Y.T. conceptualized and designed this study. Y.-L.Y. and W.-M.Z. performed the chemical synthesis and characterization. H.-Q.L., M.-X.J., and D.-Y.X. performed the *in vitro* and *in vivo* experiments. S.-T.Z., P.-J.S., and H.-L. Y. analyzed data. Q.-N.C., X.-Y.C., and H.-P.F., H.-Y.T. co-wrote the manuscript. All authors discussed the results and commented on the manuscript.

## Table of Contents

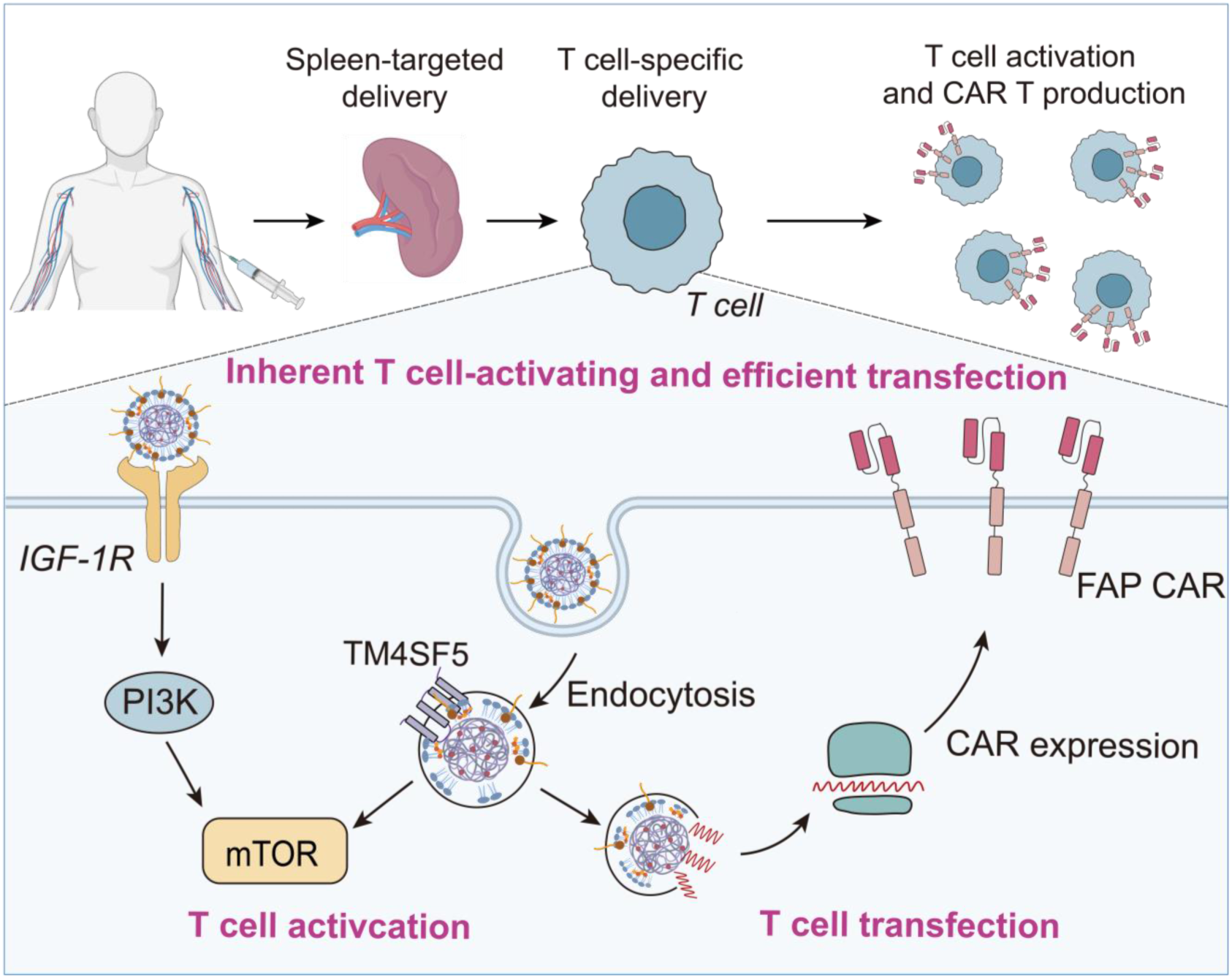

