## Supplementary material for "An inherent T cell-activating mRNA delivery platform for *in vivo* CAR T generation": All data in this study are available within the article and its supplementary information.

**Figures**

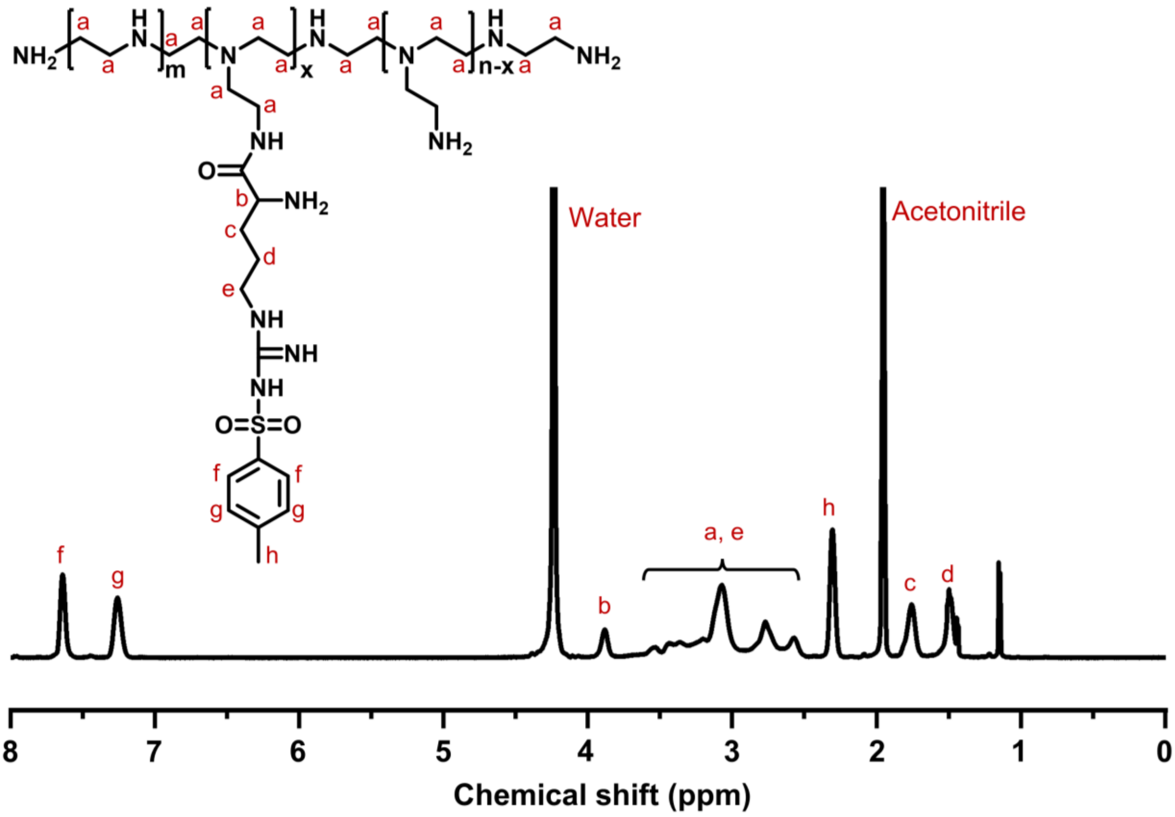


**Supplementary Fig. 1** $^{1}$H NMR spectrum of PEI-RT (500 MHz, $D_2O/CD_3CN$ (v/v=1/1)).

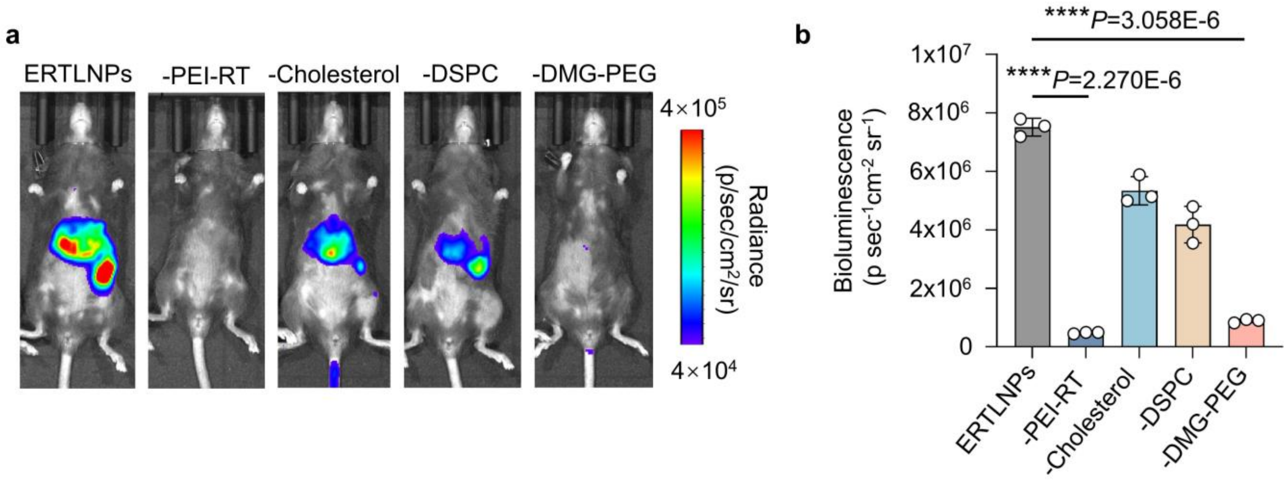


**Supplementary Fig. 2** Representative *in vivo* luciferase (Luc) bioluminescence images (**a**) and quantification data (**b**) of C57 mice 12 hours post-intravenous administration of different mRNA formations (0.5 mg $kg^{-1}$ mLuc). Data in **b** are shown as mean ± SD, n = 3 biologically independent samples. *P* values were determined by a two-tailed Student's *t*-test as indicated in the figures. ****$P < 0.0001$. A representative image of three independent samples from each group are shown in **a**.

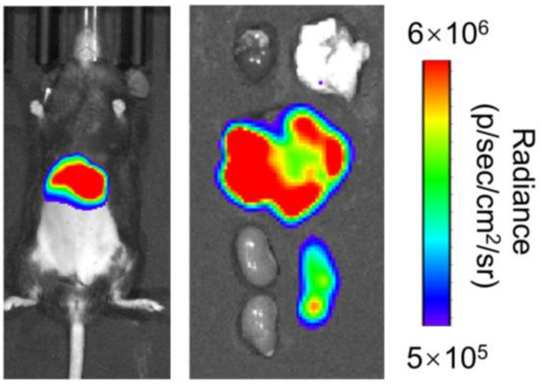


**Supplementary Fig. 3** Representative *in vivo* and *ex vivo* bioluminescence images of C57 mice 12 hours post-intravenous administration of mLuc-loaded SM102 LNPs (0.5 mg $kg^{-1}$ mLuc). A representative image of three independent samples is shown in the figure.

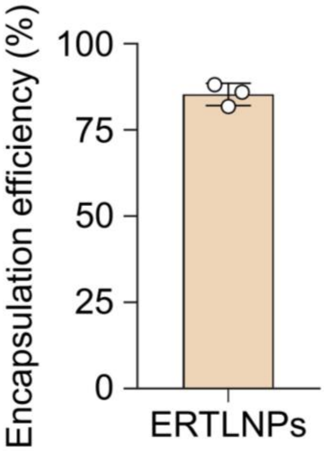


**Supplementary Fig. 4** The mRNA encapsulation efficiencies of **ERTLNPs** determined by Ribogreen assay. Data are shown as mean ± SD, n = 3 biologically independent samples.

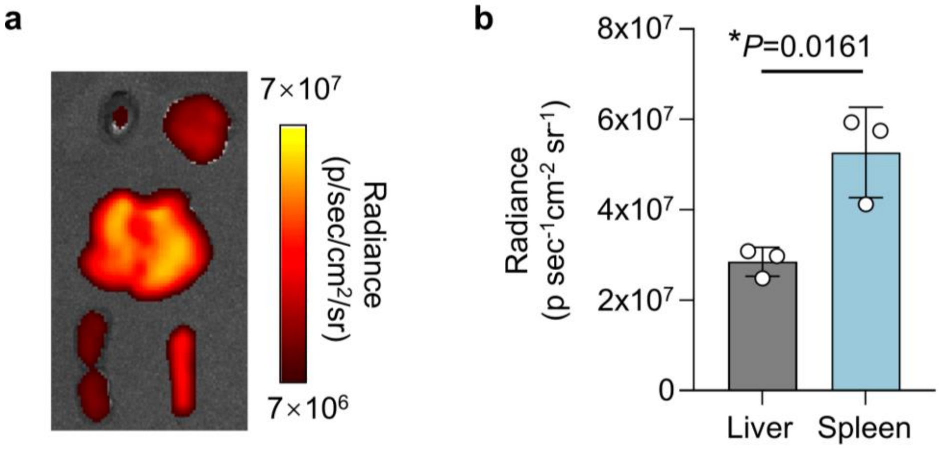


**Supplementary Fig. 5** Representative *ex vivo* biodistribution image (**a**) of C57 mice 12 hours post-intravenous administration of Cy5-mRNA loaded **ERTLNPs** (0.5 mg $kg^{-1}$ Cy5-mRNA) and quantification data (**b**) of spleen or liver. Data in **b** are shown as mean ± SD, n = 3 biologically independent samples. *P* values were determined by a two-tailed Student's *t*-test as indicated in the figures. *$P < 0.05$. A representative image of three independent samples from each group is shown in **a**.

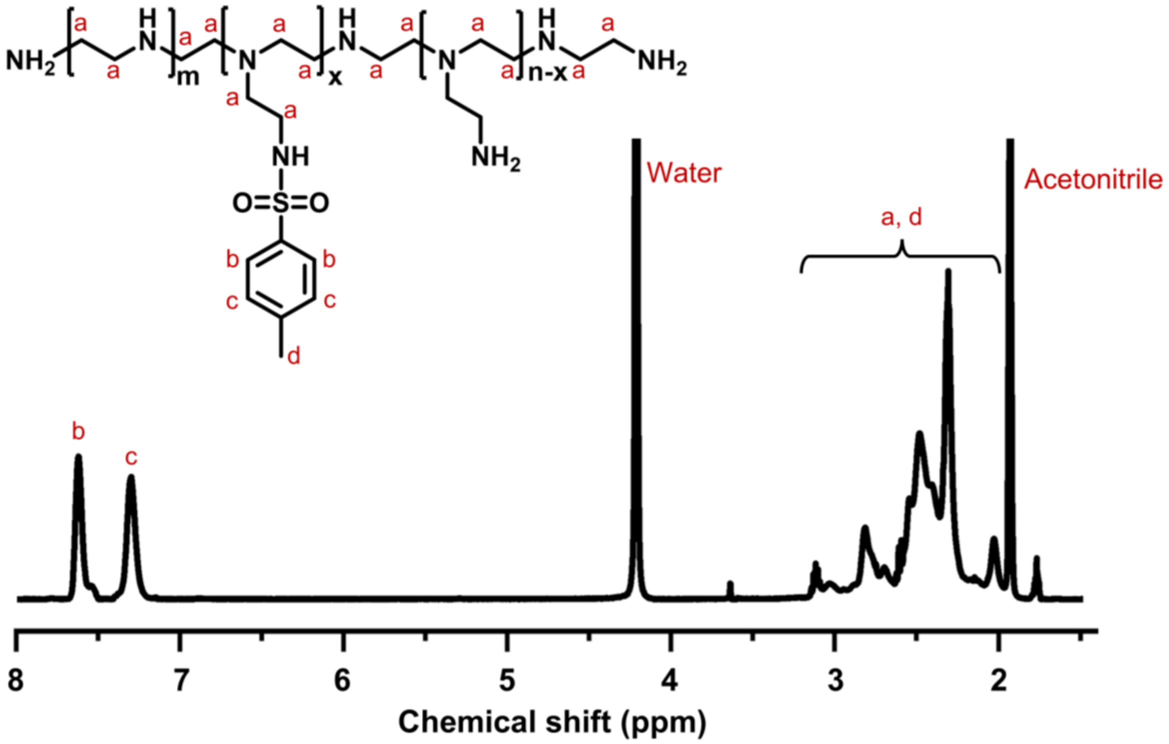


**Supplementary Fig. 6** $^{1}$H NMR spectrum of PEI-Tos (500 MHz, $D_2O/CD_3CN$ (v/v=1/1)).

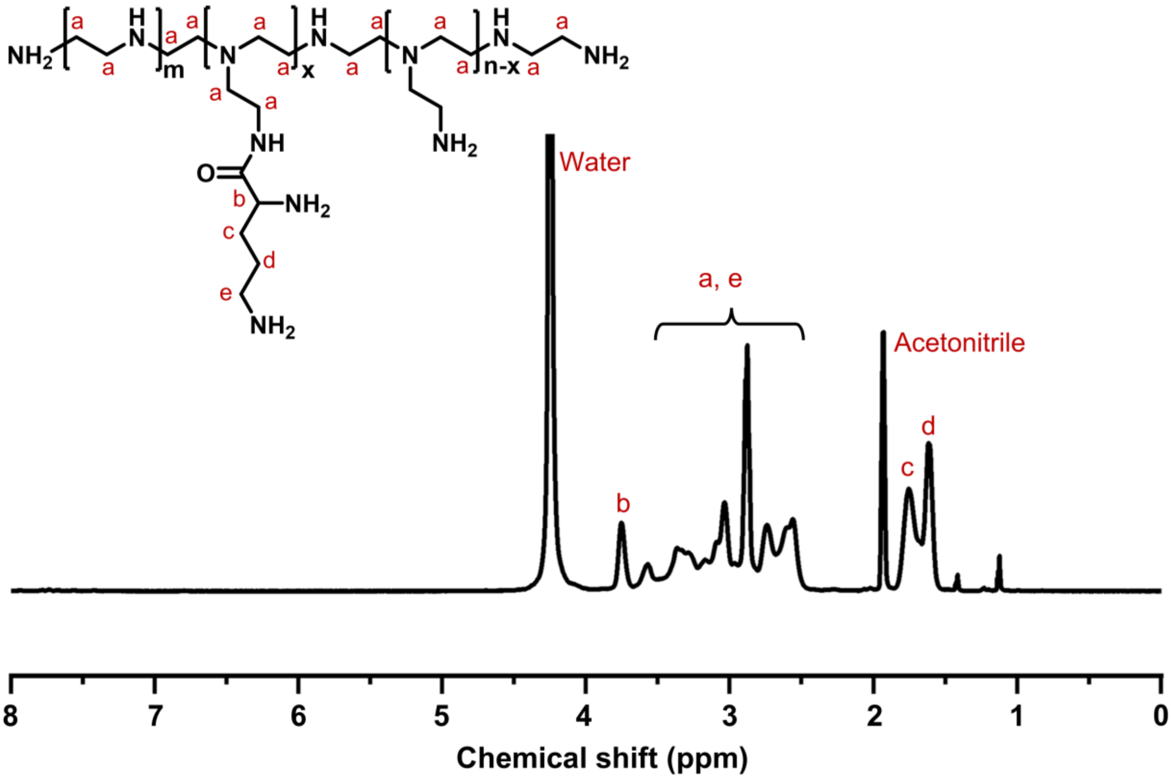


**Supplementary Fig. 7** $^{1}$H NMR spectrum of PEI-Orn (500 MHz, $D_2O/CD_3CN$ (v/v=1/1)).

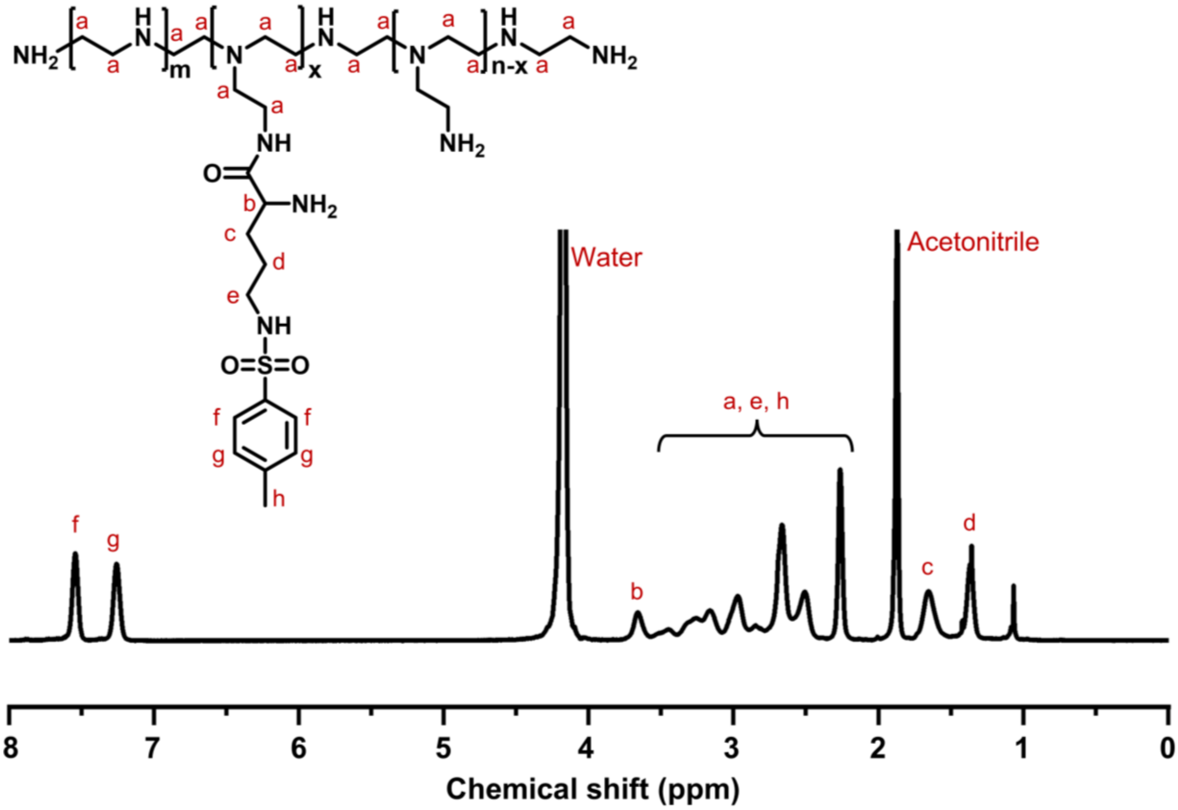


**Supplementary Fig. 8** $^{1}$H NMR spectrum of PEI-Orn(Tos) (500 MHz, $D_2O/CD_3CN$ (v/v=1/1)).

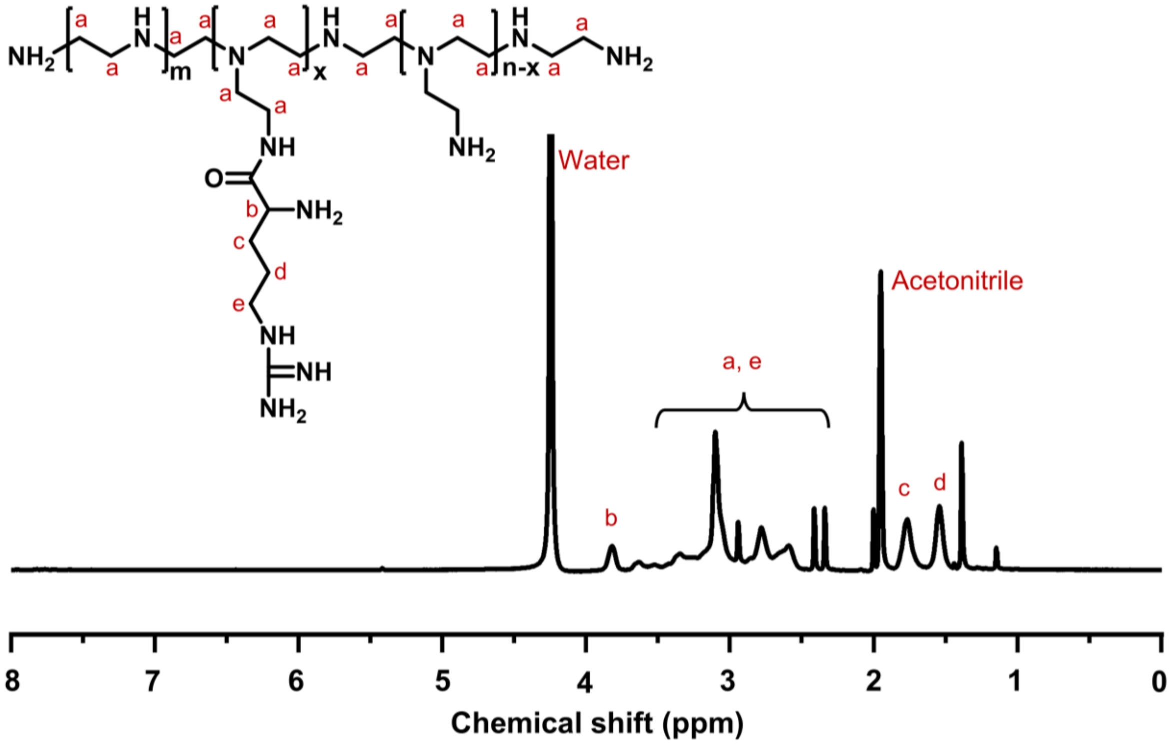


**Supplementary Fig. 9** $^{1}$H NMR spectrum of PEI-Arg (500 MHz, $D_2O/CD_3CN$ (v/v=1/1)).

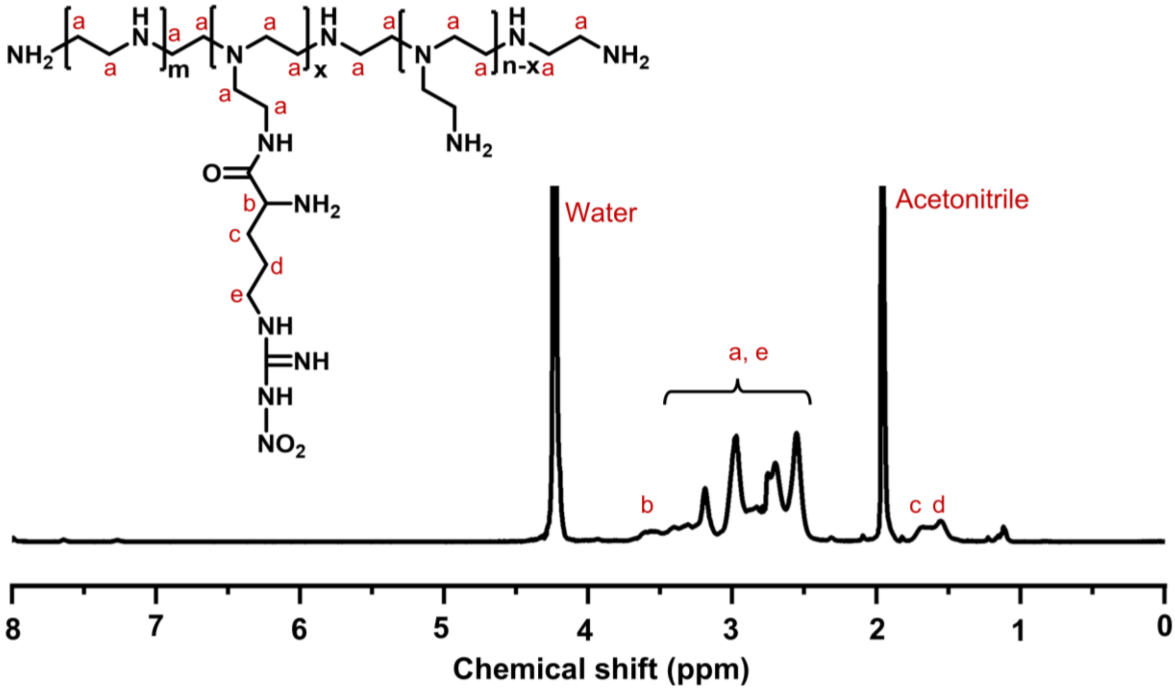


**Supplementary Fig. 10** $^{1}$H NMR spectrum of PEI-Arg ($NO_2$) (500 MHz, $D_2O/CD_3CN$ (v/v=1/1)).

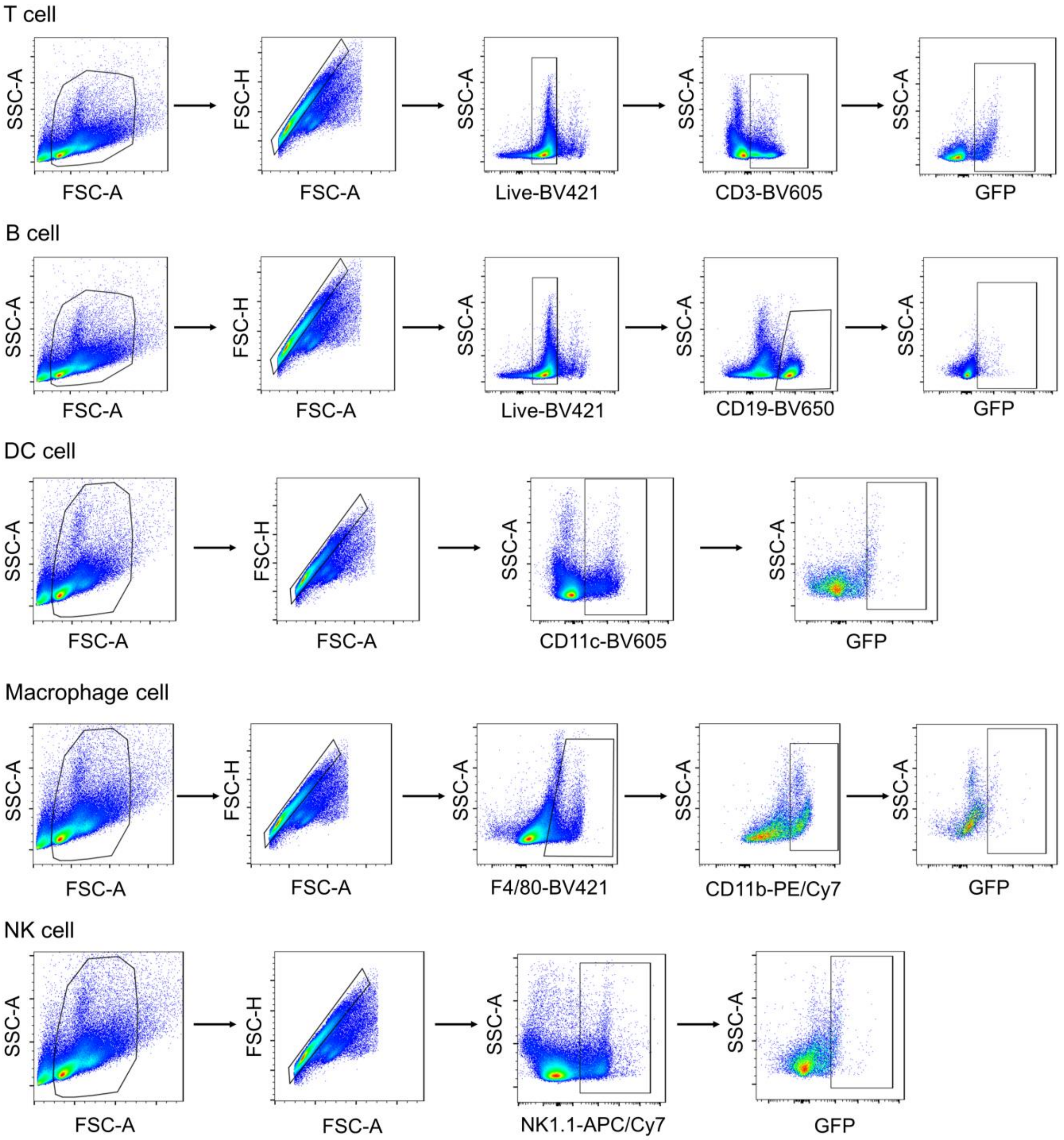


**Supplementary Fig. 11** Flow cytometry gating strategies for the analysis of GFP$^+$ cells in splenic immune cells including T cells, B cells, DCs, macrophage cells, and NK cells in mTmG mice.

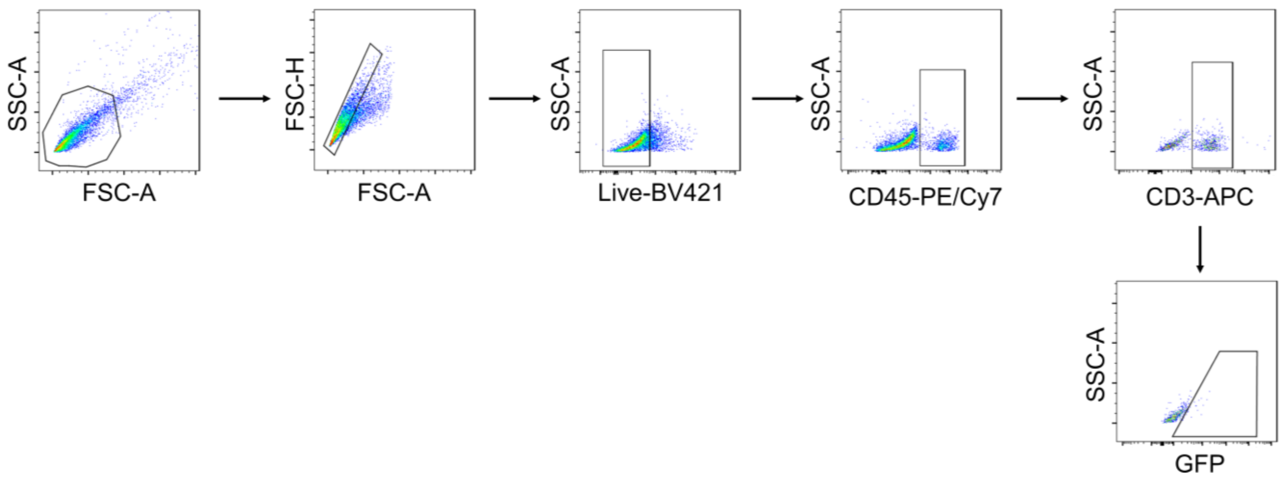


**Supplementary Fig. 12** Flow cytometry gating strategy for the analysis of $GFP^+$ cells in peripheral blood in mTmG mice.

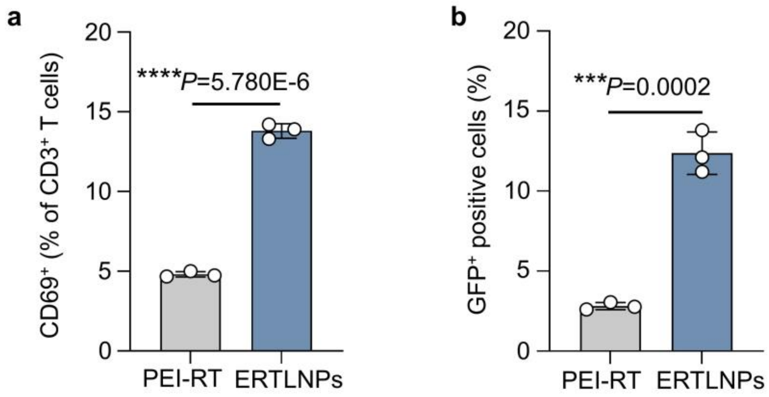


**Supplementary Fig. 13 a,b,** Relative quantification of activation efficiency ($CD69^+$ cells) (**a**) and transfection efficiency ($GFP^+$ cells) (**b**) of primary murine T cells incubated with mGFP-loaded PEI-RT or mGFP-loaded **ERTLNPs**. Data in **a** and **b** are shown as mean ± SD, n = 3 biologically independent samples. *P* values were determined by a two-tailed Student's *t*-test as indicated in the figures. $***P < 0.001$ and $****P < 0.0001$.

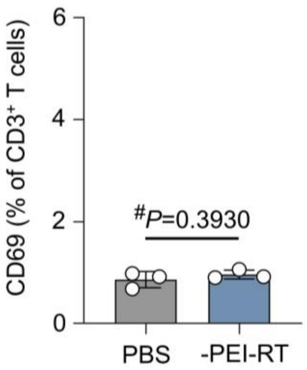


**Supplementary Fig. 14** Relative quantification of $CD69^+$ cells in primary murine T cells incubated with PEI-RT-removed **ERTLNPs**. Data are shown as mean ± SD, n = 3 biologically independent samples. *P* values were determined by a two-tailed Student's *t*-test as indicated in the figures.

$^{\#}P > 0.05$.

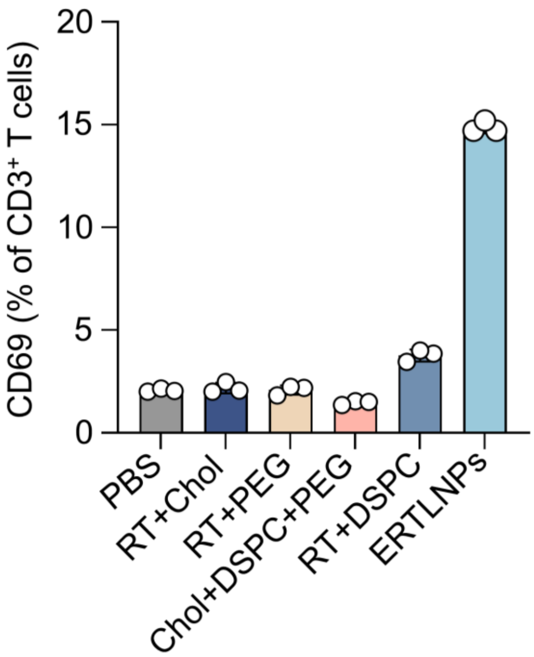


**Supplementary Fig. 15** Relative quantification of activation efficiency (CD69$^{+}$ cells) of primary murine T cells incubated with different complexes. Data are shown as mean ± SD, n = 3 biologically independent samples.

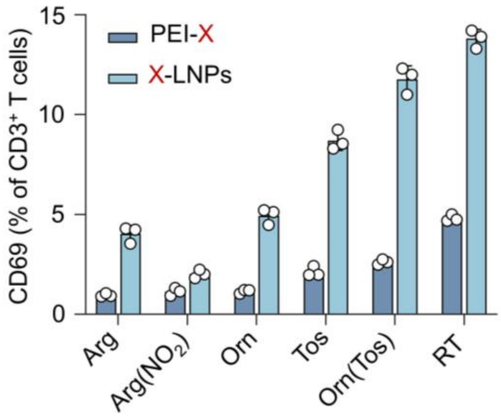


**Supplementary Fig. 16** Relative quantification of activation efficiency (CD69$^{+}$ cells) of primary murine T cells incubated with non-functional mRNA-loaded PEI-X and X-LNP (X= Arg, Arg ($NO_2$), Orn, Tos, Orn(Tos), or RT). Data are shown as mean ± SD, n = 3 biologically independent samples.

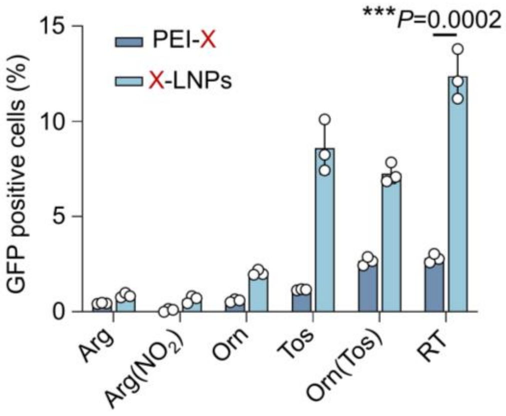


**Supplementary Fig. 17** Relative quantification of $GFP^+$ cells in primary murine T cells incubated with mGFP-loaded PEI-X and X-LNP (X= Arg, Arg ($NO_2$), Orn, Tos, Orn(Tos), or RT). Data are shown as mean ± SD, n = 3 biologically independent samples. *P* values were determined by a two-tailed Student's *t*-test as indicated in the figures. ***$P < 0.001$.

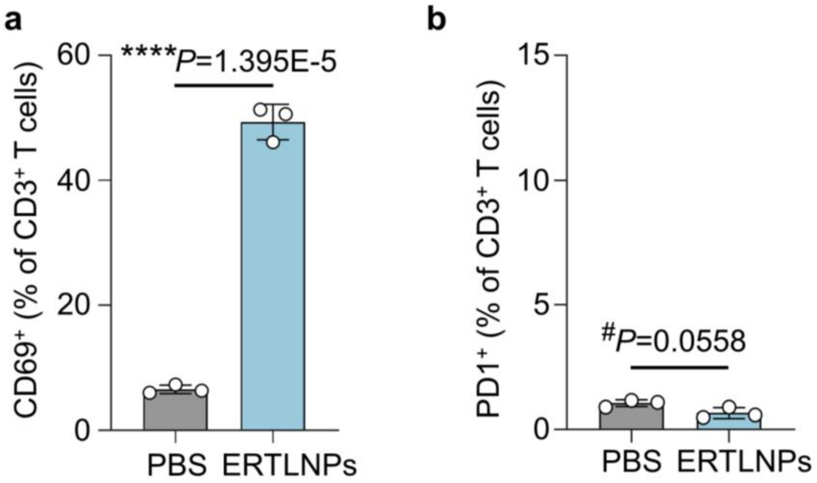


**Supplementary Fig. 18 a**,**b**, Relative quantification of $CD69^+$ cells (**a**) and PD-$1^+$ cells (**b**) in the splenic T cells of C57 mice 24 hours post-intravenously administrated with non-functional mRNA-loaded **ERTLNPs** (0.5 mg $kg^{-1}$ mRNA). Data are shown as mean ± SD, n = 3 biologically independent samples. *P* values were determined by a two-tailed Student's *t*-test as indicated in **a** and **b**. $^{\#}P > 0.05$ and ****$P < 0.0001$.

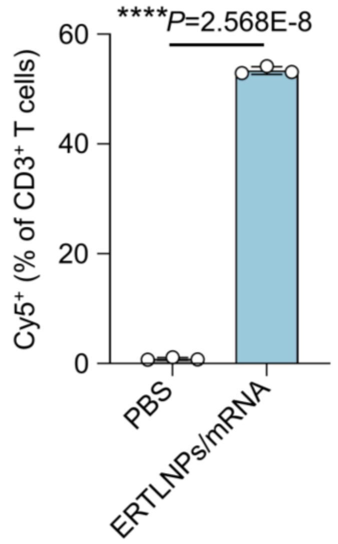


**Supplementary Fig. 19** Relative quantification of $Cy5^+$ cells in primary T cells incubated with Cy5-mRNA-loaded **ERTLNPs**. Data are shown as mean ± SD, n = 3 biologically independent samples. *P* values were determined by a two-tailed Student's *t*-test as indicated in the figure. ****$P < 0.0001$.

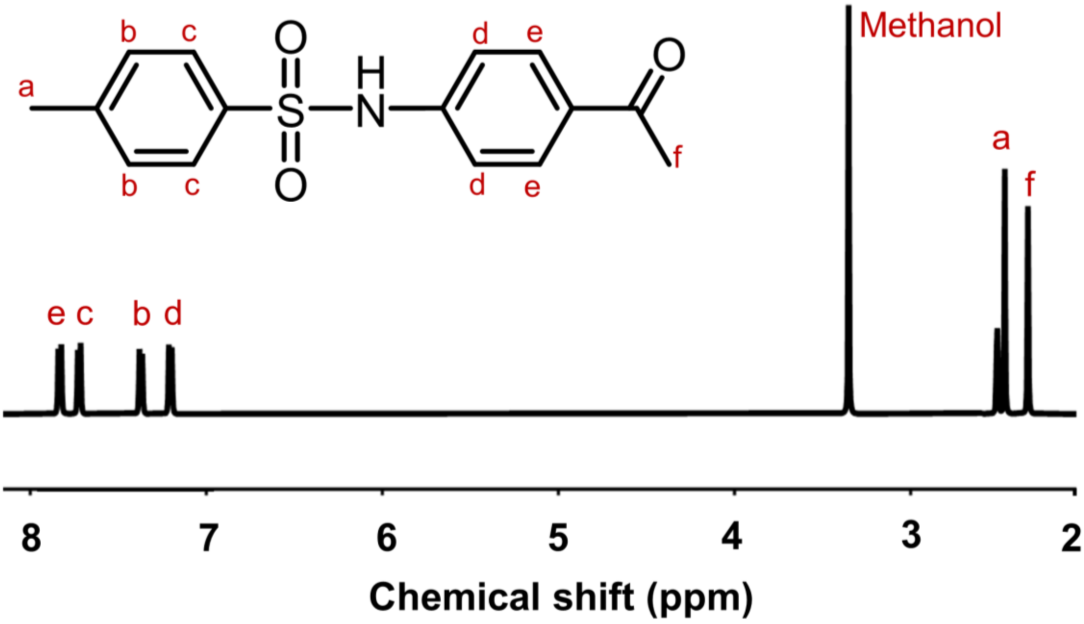


**Supplementary Fig. 20** $^1H$ NMR spectrum of TSAHC (500 MHz, $CD_3OD$).

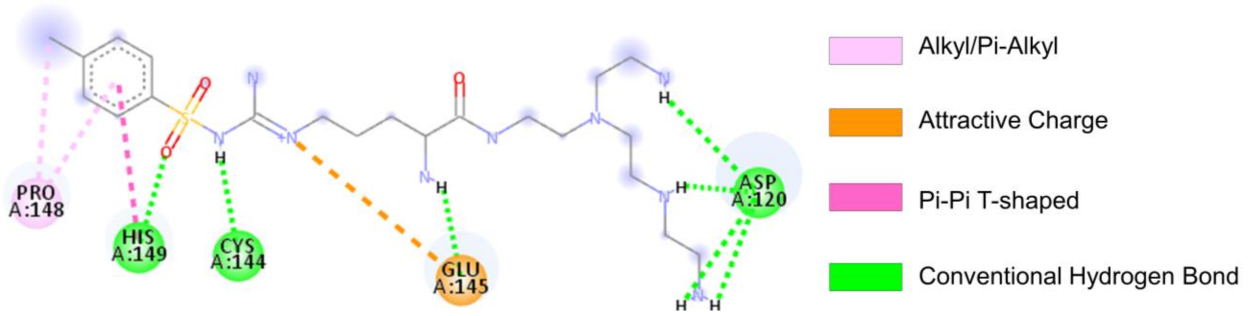


**Supplementary Fig. 21** 2D interaction map showing the predicted binding between the PEI-RT molecule and main amino acid residues within the extracellular domain of TM4SF5.

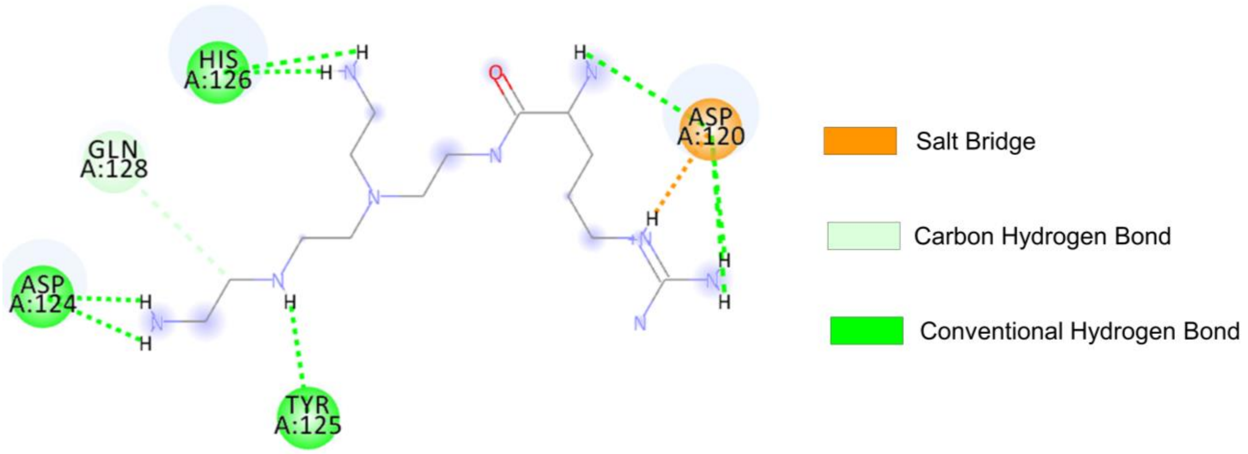


**Supplementary Fig. 22** 2D interaction map showing the predicted binding between the PEI-Arg molecule and main amino acid residues within the extracellular domain of TM4SF5.

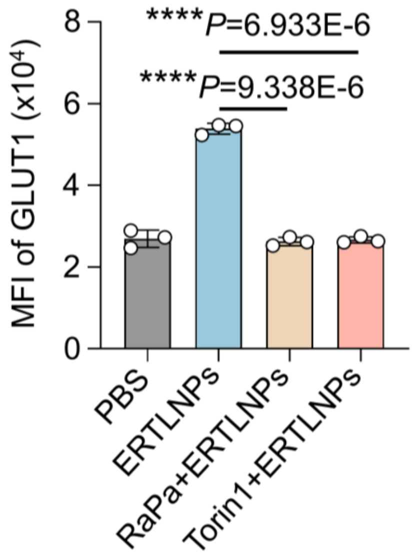


**Supplementary Fig. 23** MFI of GLUT1 expression on primary T cell incubated with **ERTLNPs-** with or without inhibitors (Rapamycin and Torin1). Data are shown as mean ± SD, n = 3 biologically independent samples. *P* values were determined by a two-tailed Student's *t*-test as indicated in the figure. ****$P$< 0.0001.

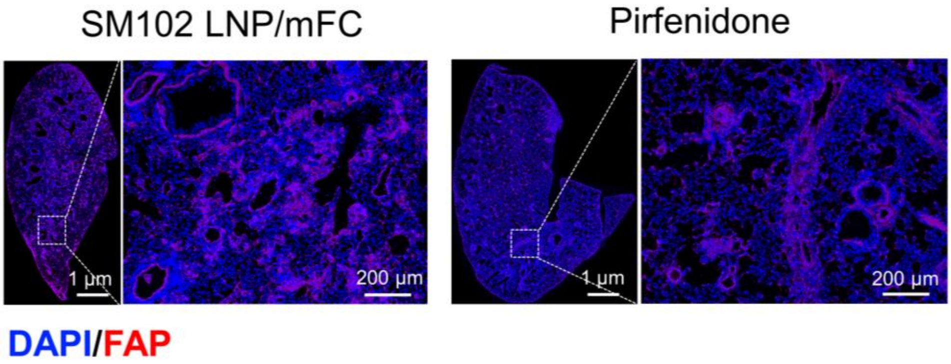


**Supplementary Fig. 24** Representative images of FAP$^+$ cells in lung tissues of IPF mice after treatment with SM102 LNP/mFC or Pirfenidone. A representative image of three independent samples from each group is shown in the figure.

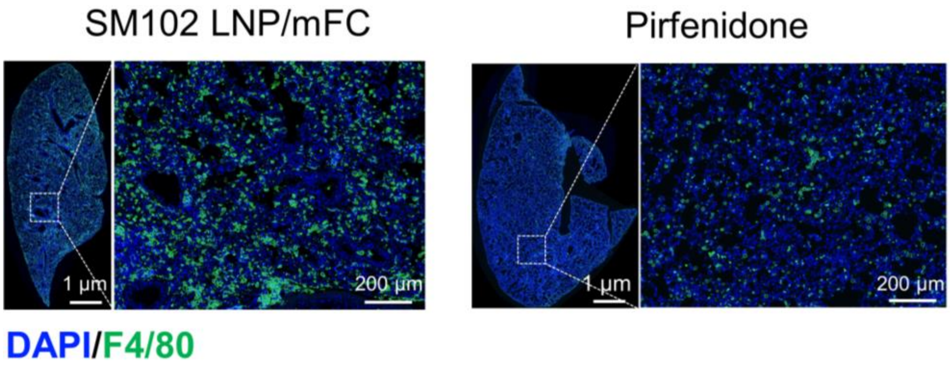


**Supplementary Fig. 25** Representative images of F4/80$^+$ cells in lung tissues of IPF mice after treatment with SM102 LNP/mFC or Pirfenidone. A representative image of three independent samples from each group is shown in the figure.

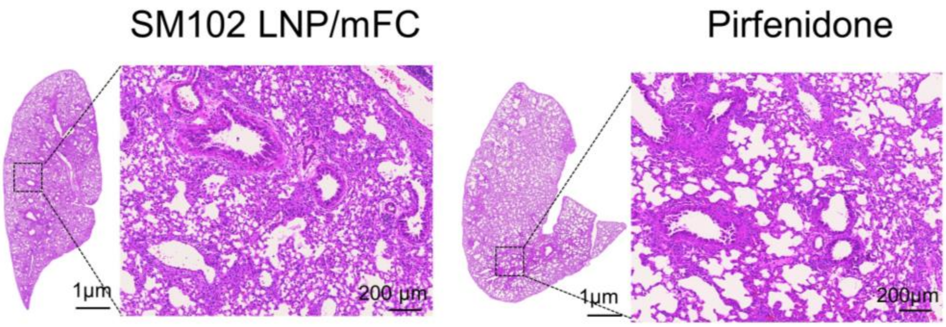


**Supplementary Fig. 26** Representative H&E staining images in lung tissue of IPF mice after treatment with SM102 LNP/mFC or Pirfenidone. A representative image of three independent samples from each group is shown in the figure.

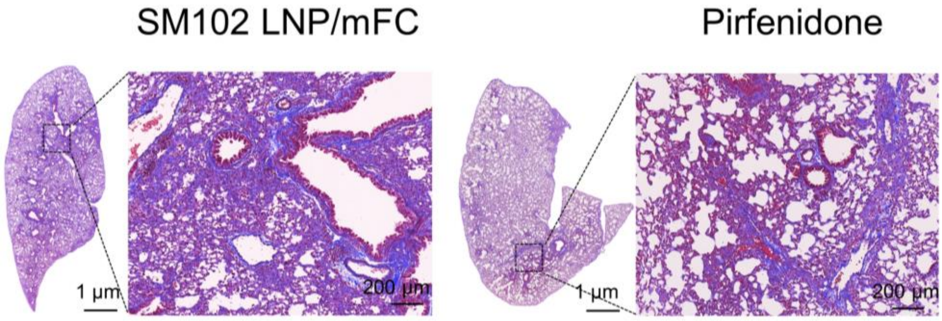


**Supplementary Fig. 27** Representative Masson's trichrome images in lung tissue of IPF mice after treatment with SM102 LNP/mFC or Pirfenidone. A representative image of three independent samples from each group is shown in the figure.

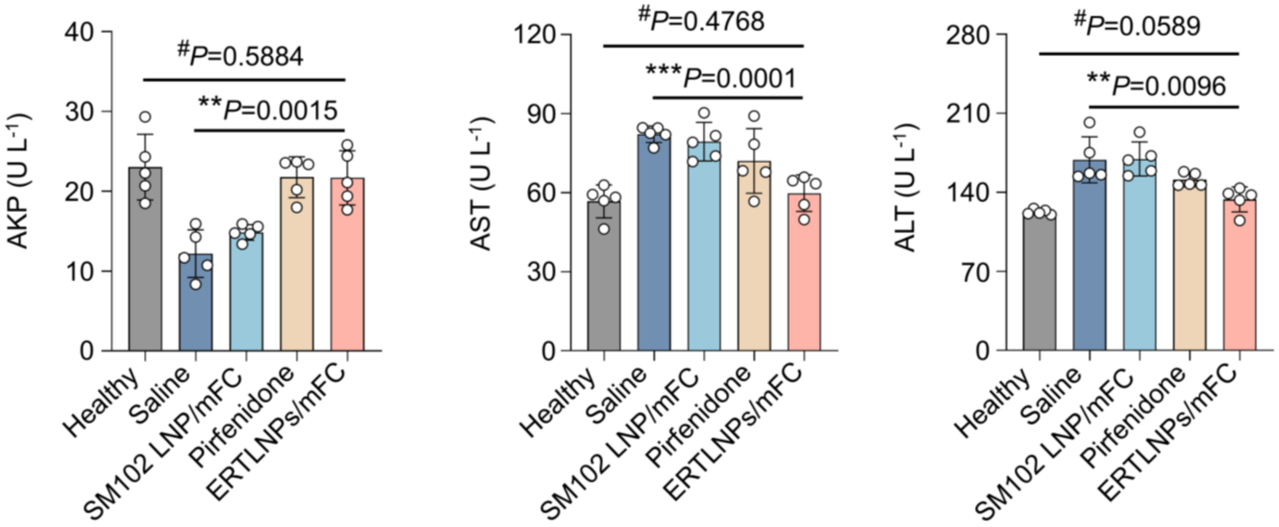


**Supplementary Fig. 28** Levels of liver function markers including aspartate aminotransferase (AST), alkaline phosphatase (AKP), and alanine aminotransferase (ALT) in serum of IPF mice after treatment with different formulations. Data are shown as mean ± SD, n = 5 biologically independent samples. *P* values were determined by a two-tailed Student's *t*-test as indicated in the figures. $^{\#}P$ > 0.05, $^{**}P < 0.01$, and $^{***}P < 0.001$.

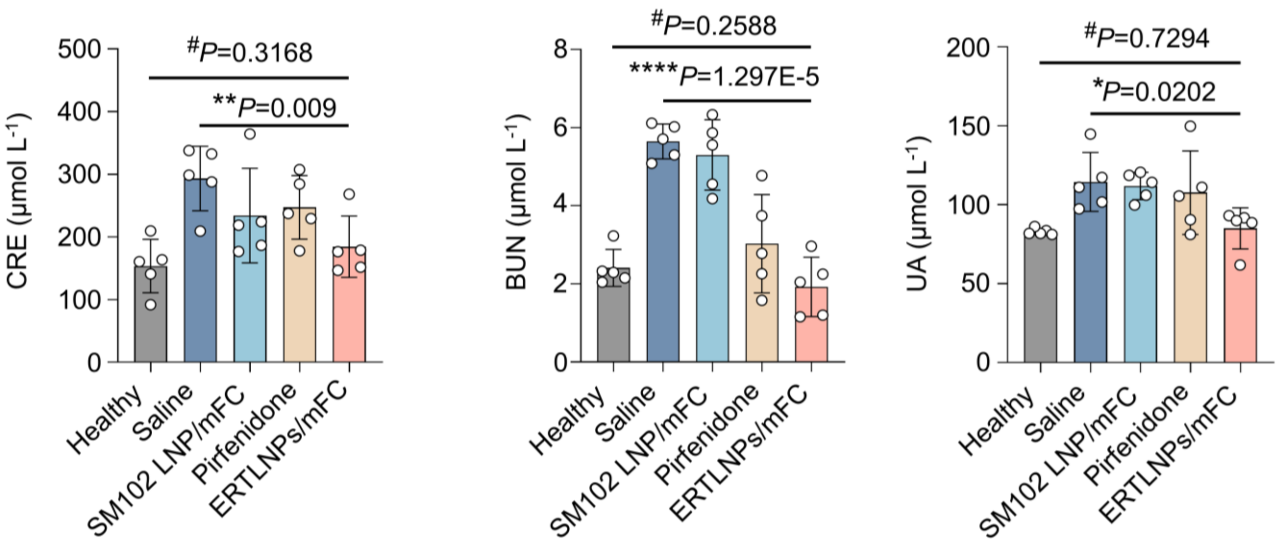


**Supplementary Fig. 29** Levels of renal function markers including uric acid (UA), urea nitrogen (BUN), and creatinine (CRE) in serum of IPF mice after treatment with different formulations. Data are shown as mean ± SD, n = 5 biologically independent samples. *P* values were determined by a two-tailed Student's *t*-test as indicated in the figures. #*P* > 0.05, **P* < 0.05, ***P* < 0.01, and ****P* < 0.001.

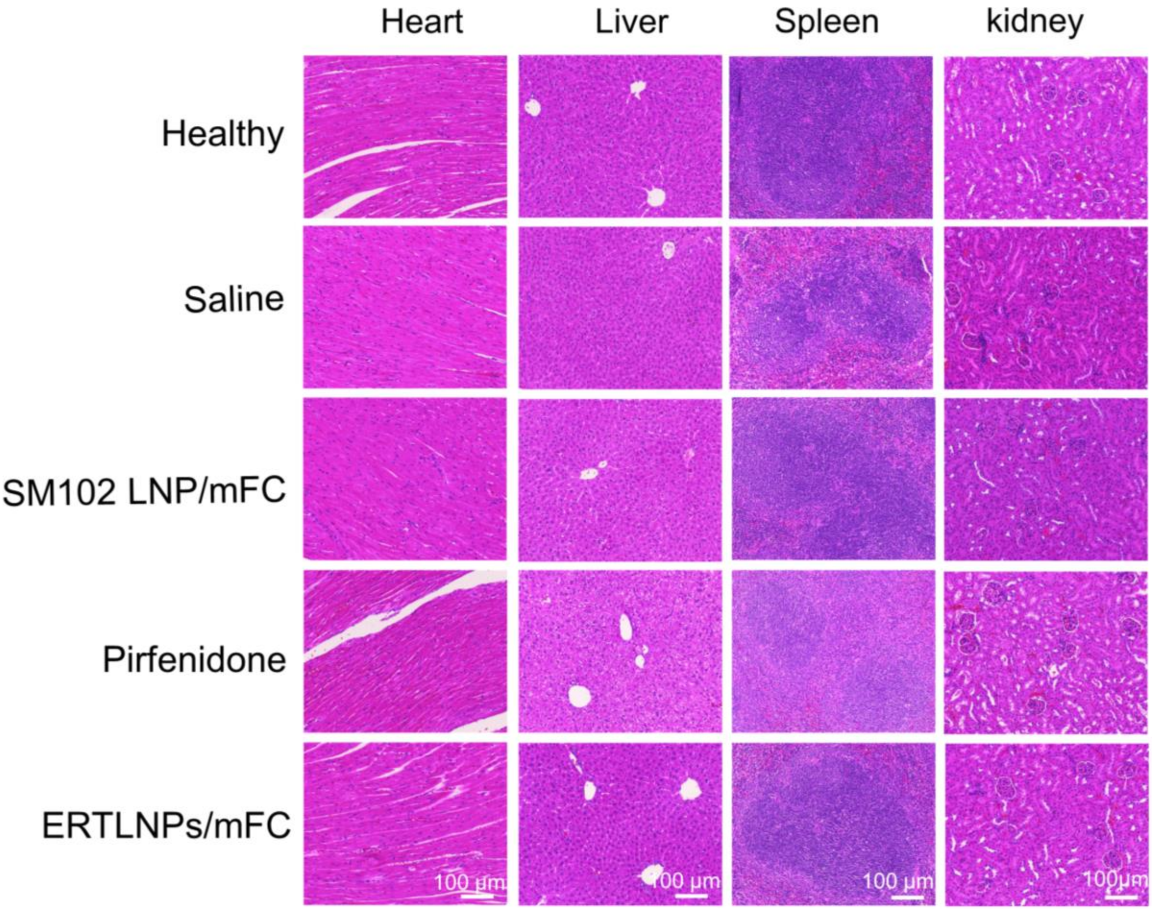


**Supplementary Fig. 30** Representative H&E staining images of main organs (heart, liver, spleen, and kidney) in IPF mice after treatment with different formulations. Scale bar: 100 μm. A representative image of five independent samples from each group is shown in the figure.

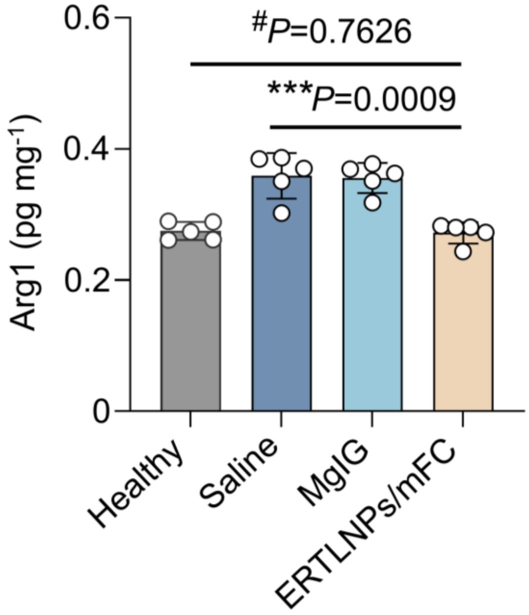


**Supplementary Fig. 31** Levels of Arg1 in liver tissues of liver fibrotic mice after treatment with different formulations. Data are shown as mean ± SD, n = 5 biologically independent samples. *P* values were determined by a two-tailed Student's *t*-test as indicated in the figures. #*P* > 0.05 and ****P* < 0.001.

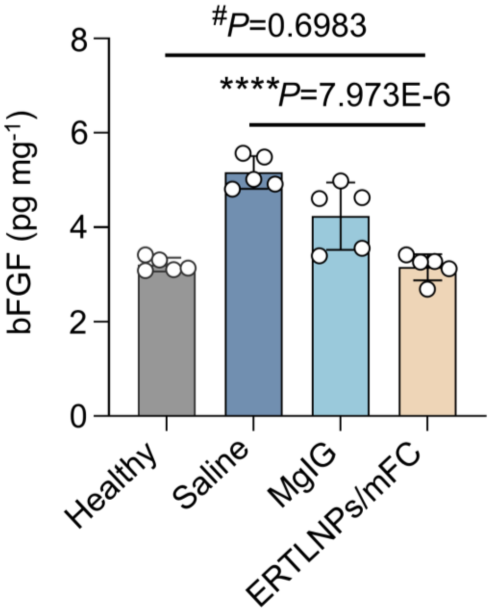


**Supplementary Fig. 32** Levels of bFGF in liver tissues of liver fibrotic mice after treatment with different formulations. Data are shown as mean ± SD, n = 5 biologically independent samples. *P* values were determined by a two-tailed Student's *t*-test as indicated in the figures. #*P* > 0.05 and ****P* < 0.001.

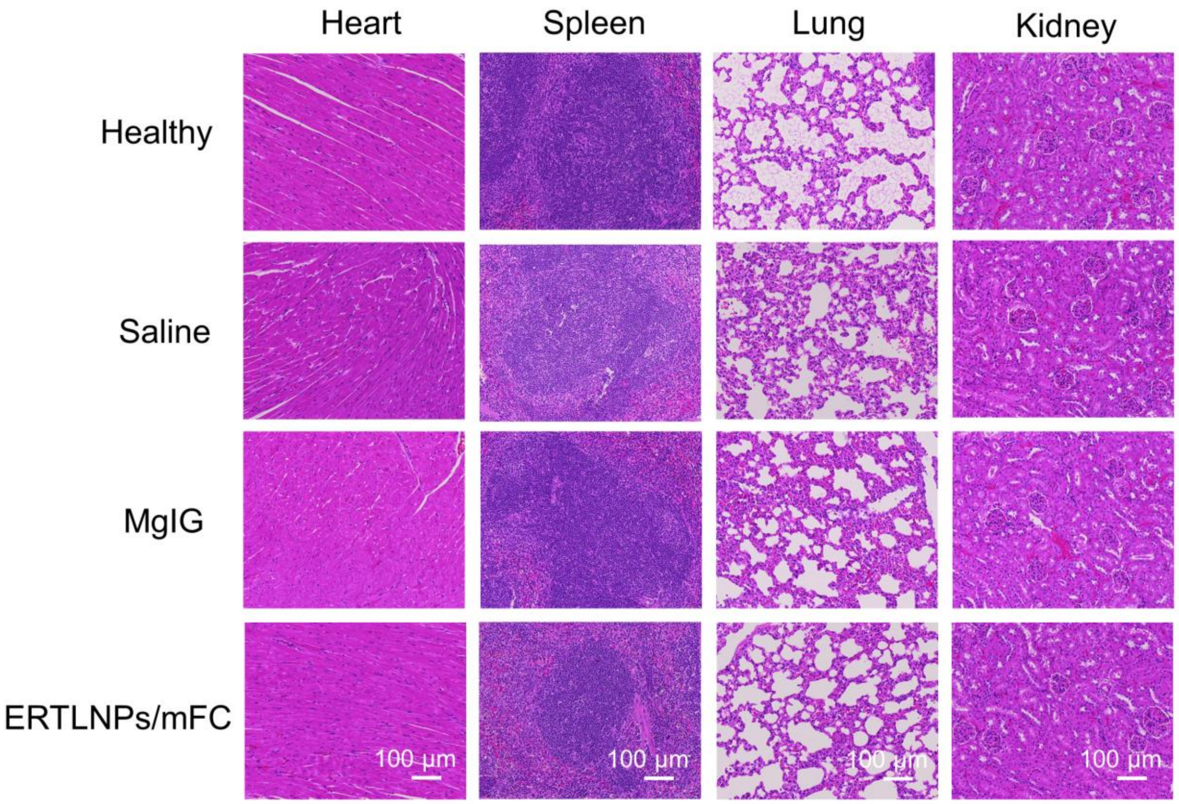


**Supplementary Fig. 33** Representative H&E staining images of main organs (heart, spleen, lung, and kidney) in liver fibrotic mice after treatment with different formulations. Scale bar: 100 μm. A representative image of five independent samples from each group is shown in the figure.

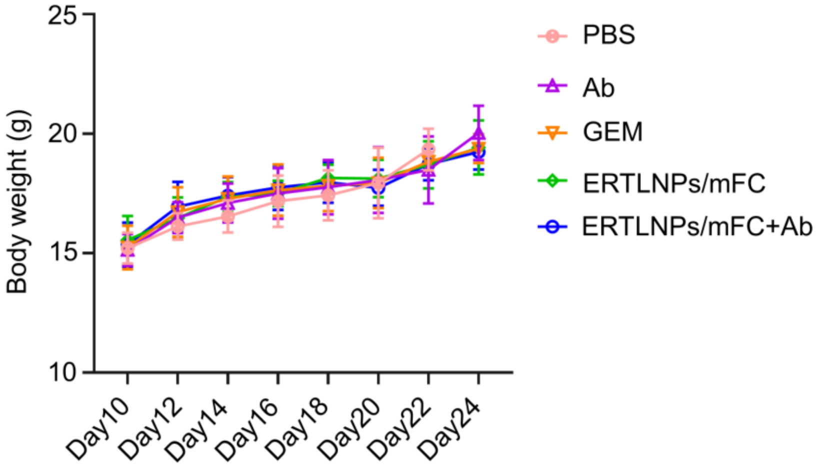


**Supplementary Fig. 34** The curves of body weight changes curves of liver fibrotic mice during treatment with different formulations. Data are shown as mean ± SD, n = 5 biologically independent samples.

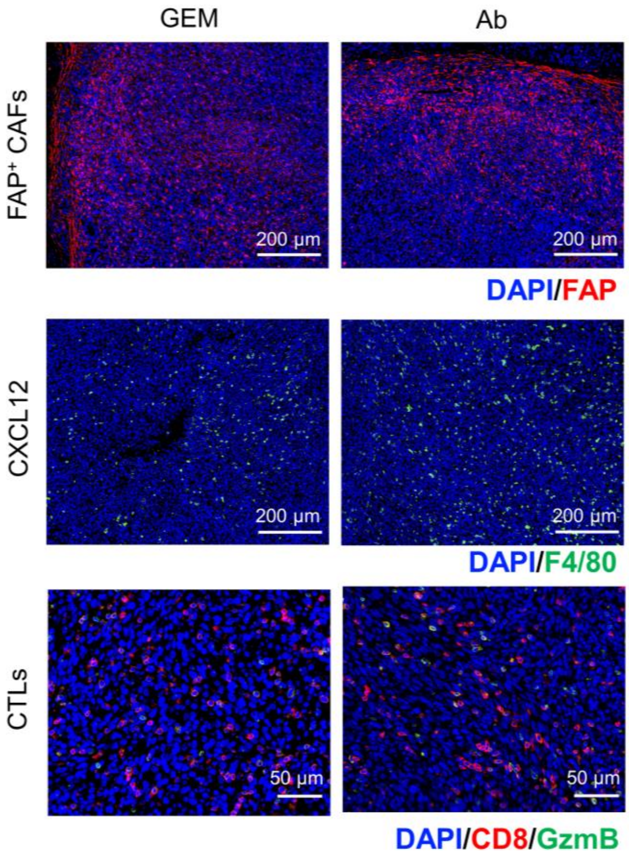


**Supplementary Fig. 35** Representative immunofluorescence images of FAP$^+$ cells (red), CXCL12 expression (green), and CTLs (Granzyme B$^+$/CD8$^+$, orange) in the tumour tissues of C57 mice with orthotopic pancreatic cancer after treatment with GEM or Ab. A representative image of three independent samples from each group is shown in the figure.

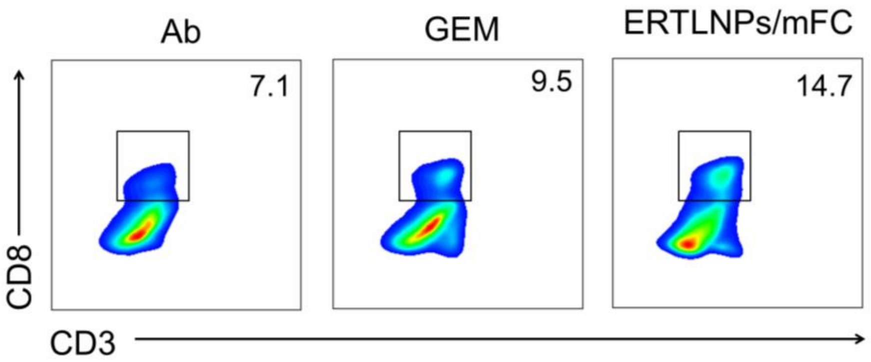


**Supplementary Fig. 36** Representative flow cytometric analysis of CD3$^+$CD8$^+$ T cells gating on CD45$^+$ cells in the tumours of C57 mice with orthotopic pancreatic cancer at the end of treatment. A representative image of five independent samples from each group is shown in the figure.

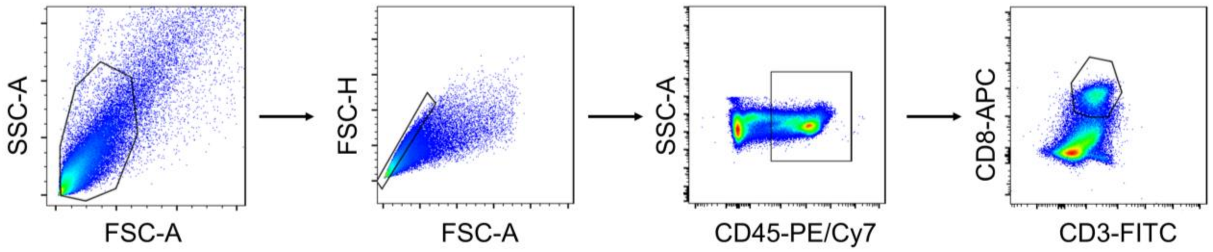


**Supplementary Fig. 37** Flow cytometry gating strategy for the analysis of CD3$^+$ CD8$^+$ T cells in tumour tissues.

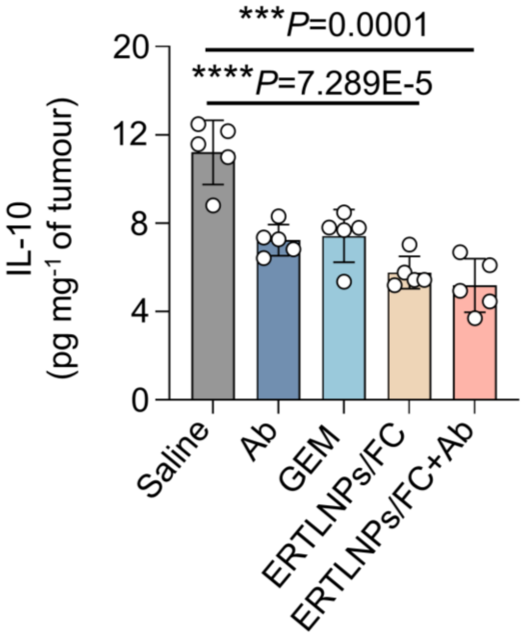


**Supplementary Fig. 38** Levels of IL-10 in tumour tissues of C57 mice with orthotopic pancreatic cancer at the end of treatment. Data are shown as mean ± SD, n = 5 biologically independent samples. *P* values were determined by a two-tailed Student's *t*-test as indicated in the figures. ***$P < 0.001$ and ****$P < 0.0001$.

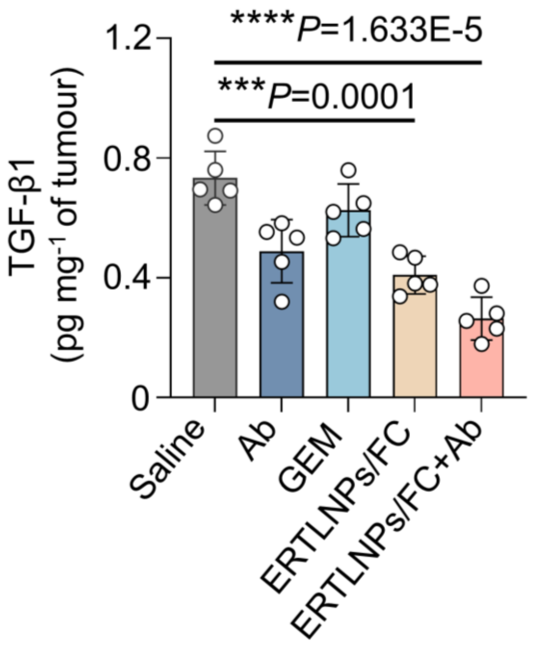


**Supplementary Fig. 39** Levels of TGF-β1 in tumour tissues of C57 mice with orthotopic pancreatic cancer at the end of treatment. Data are shown as mean ± SD, n = 5 biologically independent samples. *P* values were determined by a two-tailed Student's *t*-test as indicated in the figures. $**P < 0.01$ and $****P < 0.001$.

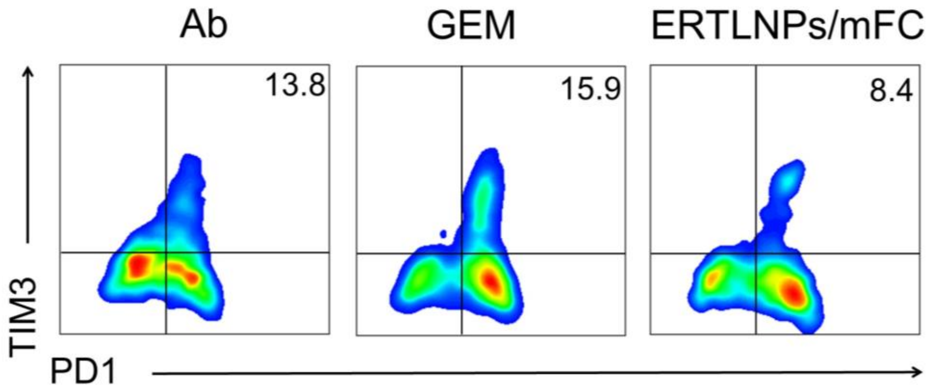


**Supplementary Fig. 40** Representative flow cytometric analysis of exhausted T cells (TIM-$3^+$PD$1^+$ CD$8^+$ T) gating on CD$45^+$ cells in the tumours of C57 mice with orthotopic pancreatic cancer at the end of treatment. A representative image of five independent samples from each group is shown in the figure.

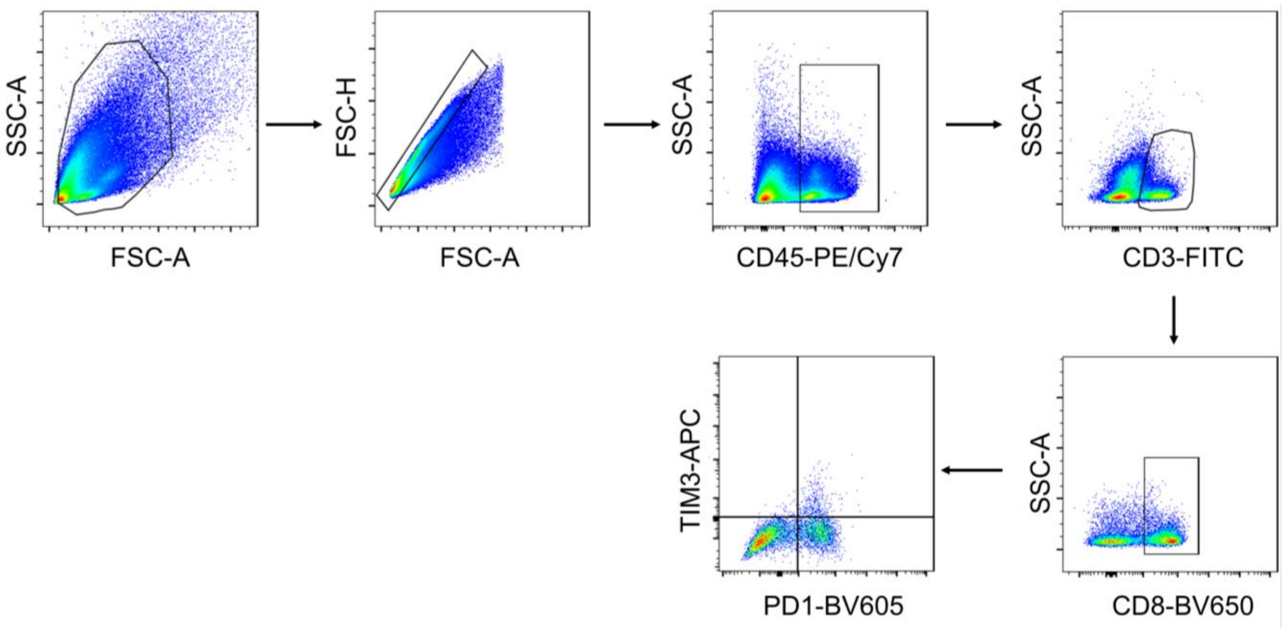


**Supplementary Fig. 41** Flow cytometry gating strategy for the analysis of exhausted T cells (TIM-3$^+$PD1$^+$CD8$^+$ T cells gating on CD45$^+$ cells) in tumour tissues.

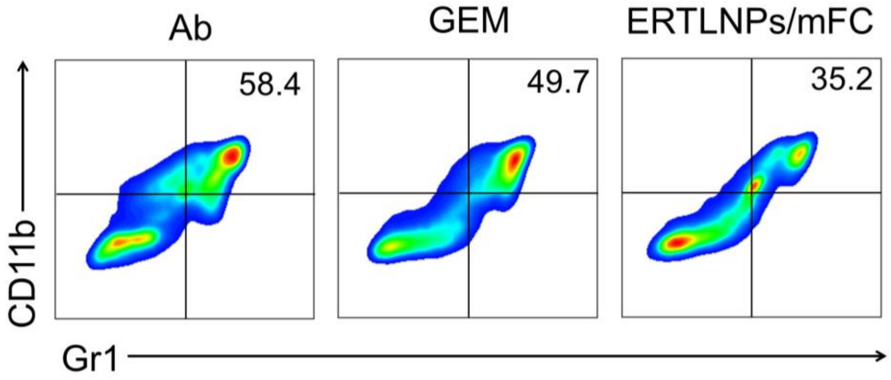


**Supplementary Fig. 42** Representative flow cytometric analysis of MDSCs (CD11b$^+$Gr-1$^+$ gating on CD45$^+$ cells) in tumour tissues of C57 mice with orthotopic pancreatic cancer at the end of treatment. A representative image of five independent samples from each group is shown in the figure.

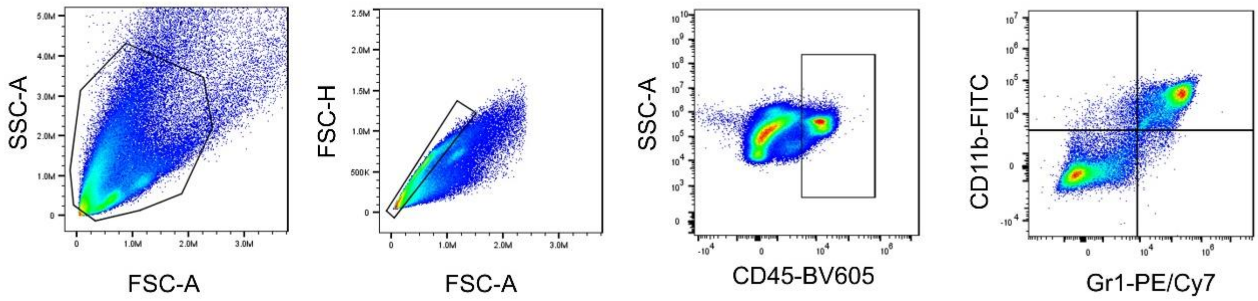


**Supplementary Fig. 43** Flow cytometry gating strategy for the analysis of MDSCs (CD11b$^+$Gr-1$^+$ gating on CD45$^+$ cells) in tumour tissues.

.

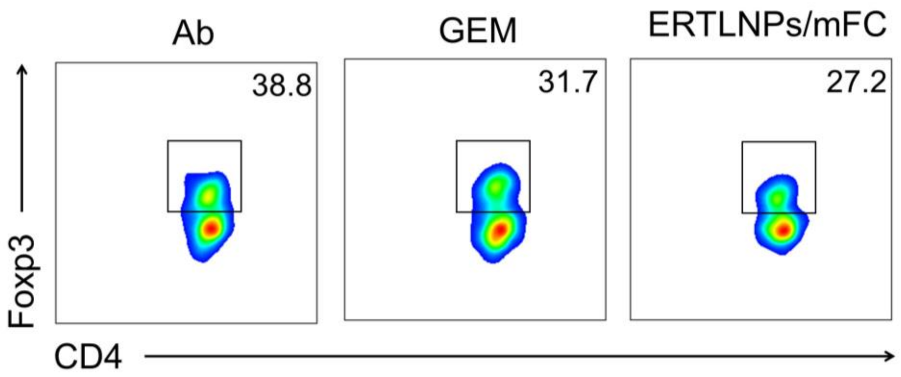


**Supplementary Fig. 44** Representative flow cytometric analysis of Tregs (Foxp3$^+$CD4$^+$ gating on CD45$^+$ CD3$^+$ cells) in tumour tissues of C57 mice with orthotopic pancreatic cancer at the end of treatment. A representative image of five independent samples from each group is shown in the figure.

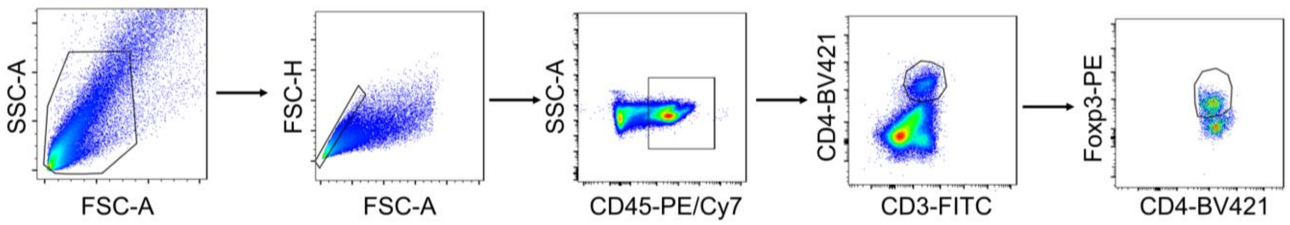


**Supplementary Fig. 45** Flow cytometry gating strategy for the analysis of Tregs (Foxp3$^+$CD4$^+$ gating on CD45$^+$ CD3$^+$ cells) in tumor tissues.

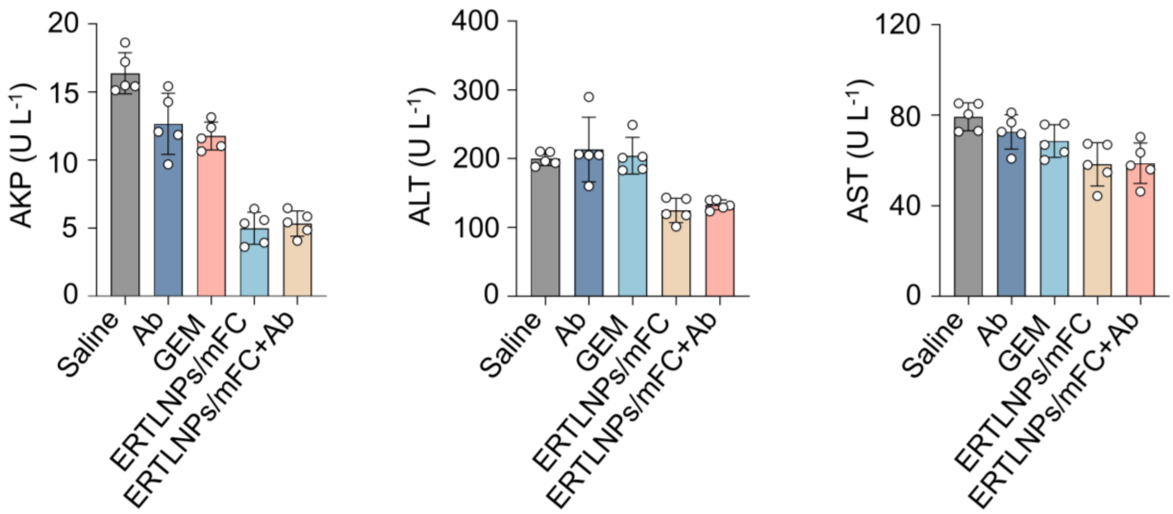


**Supplementary Fig. 46** Levels of Liver function markers including levels of AKP, ALT, and AST in serum of C57 mice with orthotopic pancreatic cancer at the end of treatment. Data are shown as mean ± SD, n = 5 biologically independent samples.

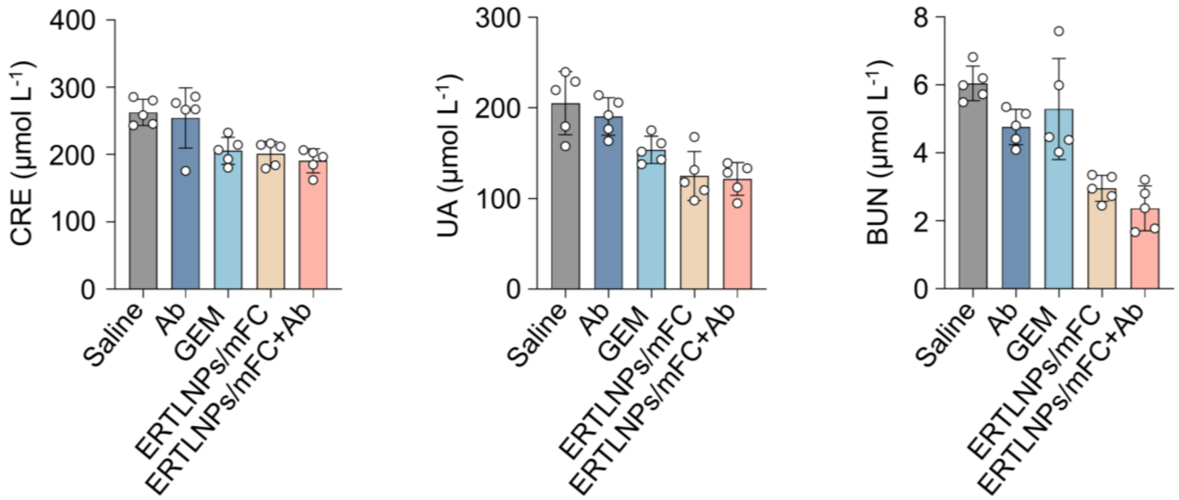


**Supplementary Fig. 47** Levels of Liver function markers including levels of CRE, UA, and BUN in serum of C57 mice with orthotopic pancreatic cancer at the end of treatment. Data are shown as mean ± SD, n = 5 biologically independent samples.

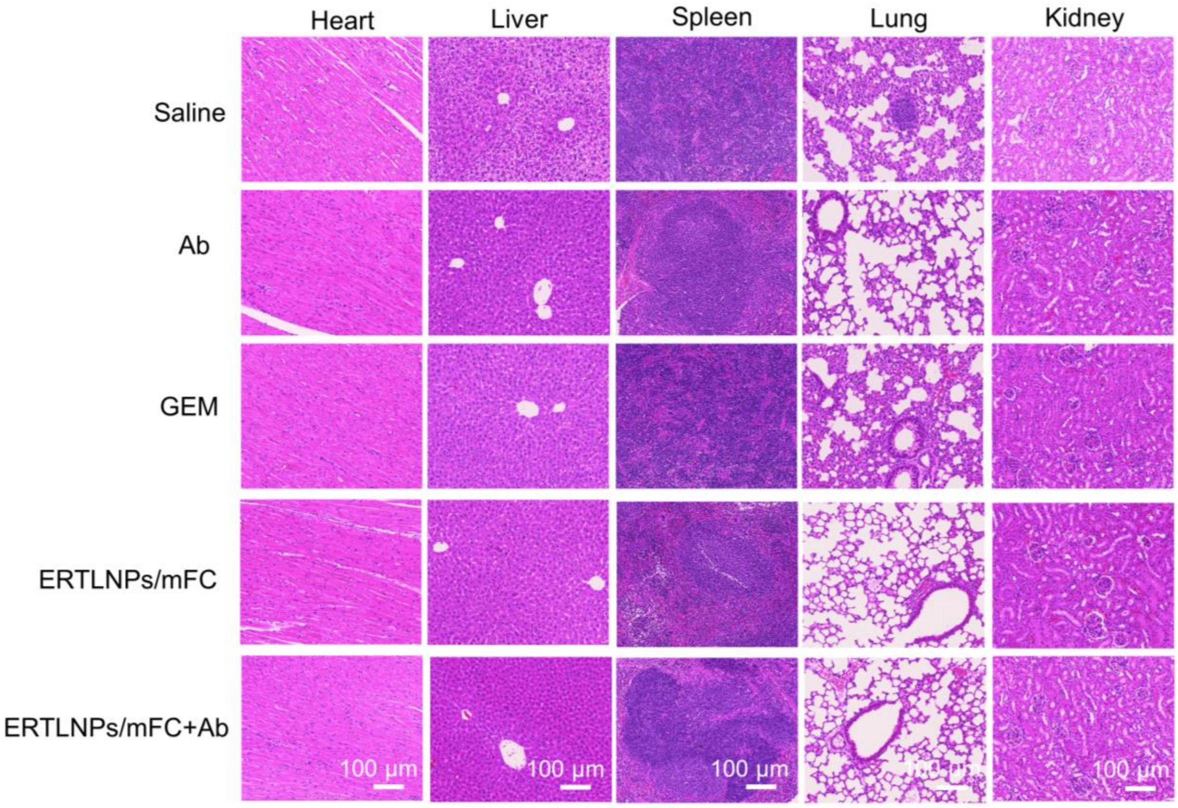


**Supplementary Fig. 48** Representative H&E staining images of main organs (heart, liver, spleen, lung, and kidney) in each group after receiving different treatments. Scale bar: 100 μm. A representative image of five independent samples from each group is shown in the figure.

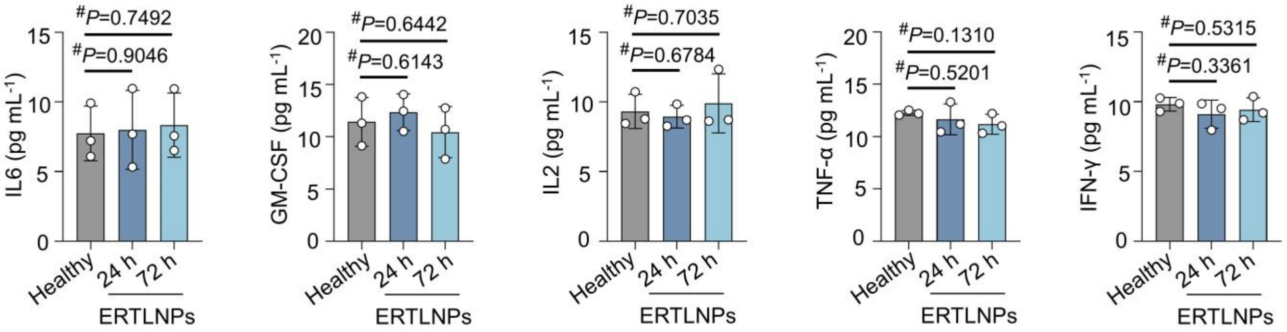


**Supplementary Fig. 49** Levels of pro-inflammatory cytokines including IL-6, GM-CSF, IL-2, IFN-γ, and TNF-α in serum at 24 and 72 hours post-intravenous injection of **ERTLNPs** to evaluate *in vivo* immunotoxicity. Data are shown as mean ± SD, n = 3 biologically independent samples. *P* values were determined by a two-tailed Student's *t*-test as indicated in the figures. $^{\#}P > 0.05$.
